# Antarctic cryptoendolithic bacterial lineages of pre-Cambrian origin as proxy for Mars colonization

**DOI:** 10.1101/2020.02.27.967604

**Authors:** Davide Albanese, Claudia Coleine, Omar Rota-Stabelli, Silvano Onofri, Susannah G. Tringe, Jason E. Stajich, Laura Selbmann, Claudio Donati

## Abstract

Cryptoendolithic communities are microbial ecosystems dwelling inside porous rocks. They are able to persist at the edge of the biological potential for life in the ice-free areas of continental Antarctica. These areas include the McMurdo Dry Valleys, often cited as a Terrestrial analog of the Martian environment. Despite their interest as a model for the early colonization by living organisms of terrestrial ecosystems and for adaptation to extreme conditions of stress, little is known about the evolution, diversity and genetic makeup of bacterial species that reside in these environments. We performed metagenomic sequencing of 18 communities from rocks collected in Antarctic desert areas over a distance of about 350 km. A total of 469 draft bacterial genome sequences were assembled, and clustered into 269 candidate species that lack a representative genome in public databases. The majority of these new species belong to monophyletic bacterial clades that diverged from related taxa in a range from 1.2 billion to 410 Ma, much earlier than the glaciation of Antarctica, and that are functionally distinct from known related taxa. The hypothesis that Antarctic cryptoendolithic bacterial lineages were generated by the selection of pre-existing cold-tolerant organisms whose origin dates back to the Tonian glaciations gives new insights for the possibility of life on Mars.

## Main

The earliest terrestrial niche for life on Earth were rocks that offered protection from UV-radiation (especially before screening pigments were evolved^1^), desiccation and other stresses^2^. Porous rocks in particular remain the ultimate refuge for life in harsh environments as in the ice-free areas of Antarctica, where complex life-forms became extinct about 60-30 Ma, when the continent reached the South Pole and the Antarctic Circumpolar Current was established. If putative Martian life ever evolved, it may have had the same fate finding a last refuge within the rock airspaces during the cooling of the Red Planet. Therefore, Antarctic cryptoendolithic communities are accounted as proxies for searching life on Earth-like planets as Mars. The McMurdo Dry Valleys, covering a surface of approximately 4,800 square km in continental Antarctica, are among the most extreme regions on Earth with only minimal resources suitable for supporting life^3,4^. In these desert areas, where soils have been eroded by glaciers and strong winds, life is confined to the endolithic niche, that provides microorganisms with protection from abiotic stresses and access to mineral nutrients, rock moisture and growth surfaces^5,6^. The endolithic environment is a ubiquitous habitat for microorganisms on the Earth^6,7^, but in the most extreme terrestrial climates it is often the primary or even exclusive refuge of life^8^.

Endolithic microbial communities are self-sustaining ecosystems relying on the phototrophic activity of microalgae and Cyanobacteria as primary producers which support a diversity of consumers including Fungi, Bacteria and Archaea^5,9–11^. In the Antarctic desert areas, the Lichen-Dominated Communities (LDC) are the most complex and successful^5^. Recently, next-generation sequencing studies have brought new insights into their composition, showing that Lecanoromycete lichens and free living fungi in the Dothideomycetes (Ascomycota) are the dominant heterotrophic eukaryotes, while Actinobacteria and Proteobacteria are the most abundant Bacteria^12,13^. Due to their ubiquity in deserts and low taxonomic complexity and biodiversity^14^, endoliths are important study systems to understand evolutionary processes in the early history of life, to model how life evolves during the progression of desertification, and the extreme aridity approaches the limits of life providing a model for searches for life elsewhere in the Solar System. However, the understanding of the microbial biodiversity in these communities is limited and our comprehension about their physiology and stress responses is still at its infancy^15^.

In this study, we performed metagenomic sequencing of eighteen LDC-colonized rock samples (Fig. 1a) collected in Antarctic ice-free areas distributed over a distance of 350 km (Fig. 1b,c) to provide an initial survey of the genomic repertoire of bacteria from endolithic ecosystems^16^. The metagenomic assemblies generated more than 10 million contigs which were binned into 497 novel bacterial genomes and classified as 269 previously unknown species-level clusters, substantially expanding the sampled genomic diversity within 33 bacterial orders. The new taxa define lineages that are specific to these communities and according to a molecular clock analysis, diverged from their closest known bacterial lineages much earlier than the estimated time of separation of Antarctica from Gondwanaland. The microbial lineages in these communities are likely a product of selection of existing extremo-tolerant organisms, and not by development of novel genetic traits in response to environmental conditions. Whether these organisms will adapt and survive in the face of fast-occurring climate changes remains an important and open question.

**Fig. 1.**
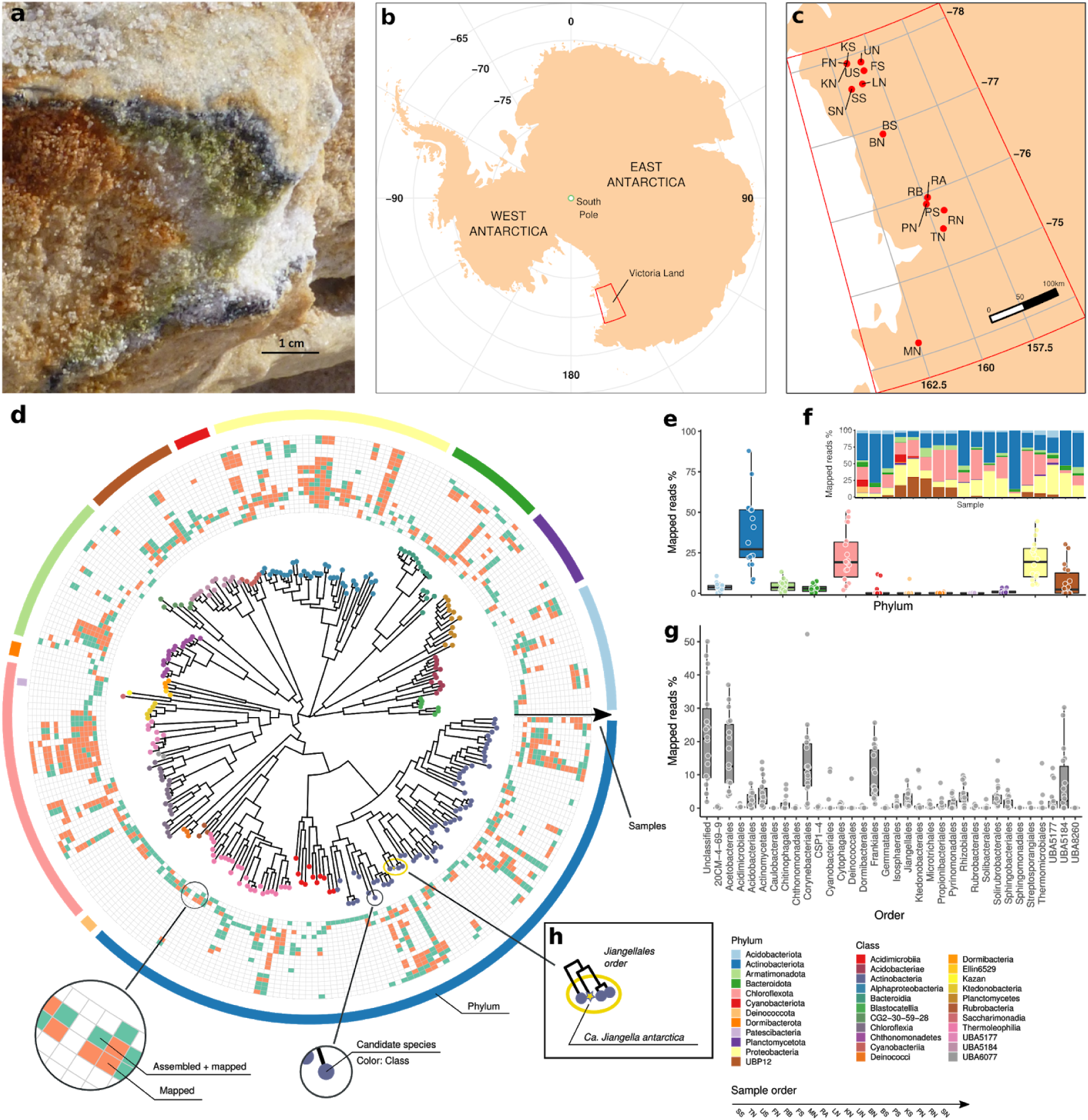
**a)** Cryptoendolithic lichen-dominated community colonizing sandstone at Linnaeus Terrace, McMurdo Dry Valleys, Southern Victoria Land, Continental Antarctica. **b)** Map of Antarctica. The area encircled by the red lines represents Victoria Land. **c)** Map of the study area showing the location of the sampling sites. Sampled sites are listed in Supplementary Table 4. **d)** Phylogenetic tree built from the GTDB-Tk multiple sequence alignment (MSA) of the 269 CBS representatives. Tip points are colored according to the GTDB taxonomic classification at the class level. Phylum-level classification is indicated by the colors in the outer circle. The 18 inner circles represent distribution of each CBS in the samples. Presence inferred by the assembly is indicated in green, while presence inferred from the alignment of the read to the CBS representative genome is indicated in orange. **e)** Percentage of reads that could be mapped to the CBS representatives, grouped by Phylum. **f)** Per sample percentage of the reads that could be mapped to the CBS representatives, grouped by Phylum. **g)** Same as **e**, at the order taxonomic level. **h)** Jiangellales CBS, including the *Candidatus Jiangella antarctica* (yellow star).

### Metagenomic assembly identifies novel bacterial species and broadly expands the tree of life

Using shotgun sequencing and assembly, we produced more than 10 million contigs that were binned into a total of 1660 metagenome-assembled genomes (MAGs), among which 497 were identified as bacterial and none as archaeal. The bacterial MAGs were partitioned into 263 high quality (HQ) and 234 medium quality (MQ) according to their estimated completeness and contamination (see Methods). Assembly, completeness and contamination statistics and the taxonomic classification of the 497 bacterial MAGs are given in Supplementary Table 1. Species-level dereplication of the MAGs produced a set of 269 clusters - or candidate bacterial species (CBS) - each represented by the MAG of highest quality. The CBSs were taxonomically classified using GTDB-Tk^17^ (see Methods). While all CBSs could be assigned to a known phylum or class, none could be classified into existing species (Table 1). The most common phylum, both in terms of number and abundance of CBS (estimated by the fraction of mapped reads, Fig. 1e, f, Supplementary Table 2), was *Actinobacteria* with 101 CBS (median percentage of mapped reads 27.2%, IQR 29.5%), followed by *Chloroflexi* and *Proteobacteria*, that are ‘core’ members of Antarctic endolithic communities^12^. The identified MAGs significantly increase the repertoire of known genomic data for several taxa, and represent the first example of bacterial genomes recovered from endolithic communities. Examining the represented orders, the newly assembled MAGs increase by more than 50% the number of representative species in the Genome Taxonomy Database^18^ (GTDB) for *Jiangellales, Frankiales, Thermomicrobiales, Isosphaerales, Solirubrobacterales* and for the order-level *UBA5184* UBA lineage^19^ (Supplementary Table 3, Fig. 1g).

**Table 1.**
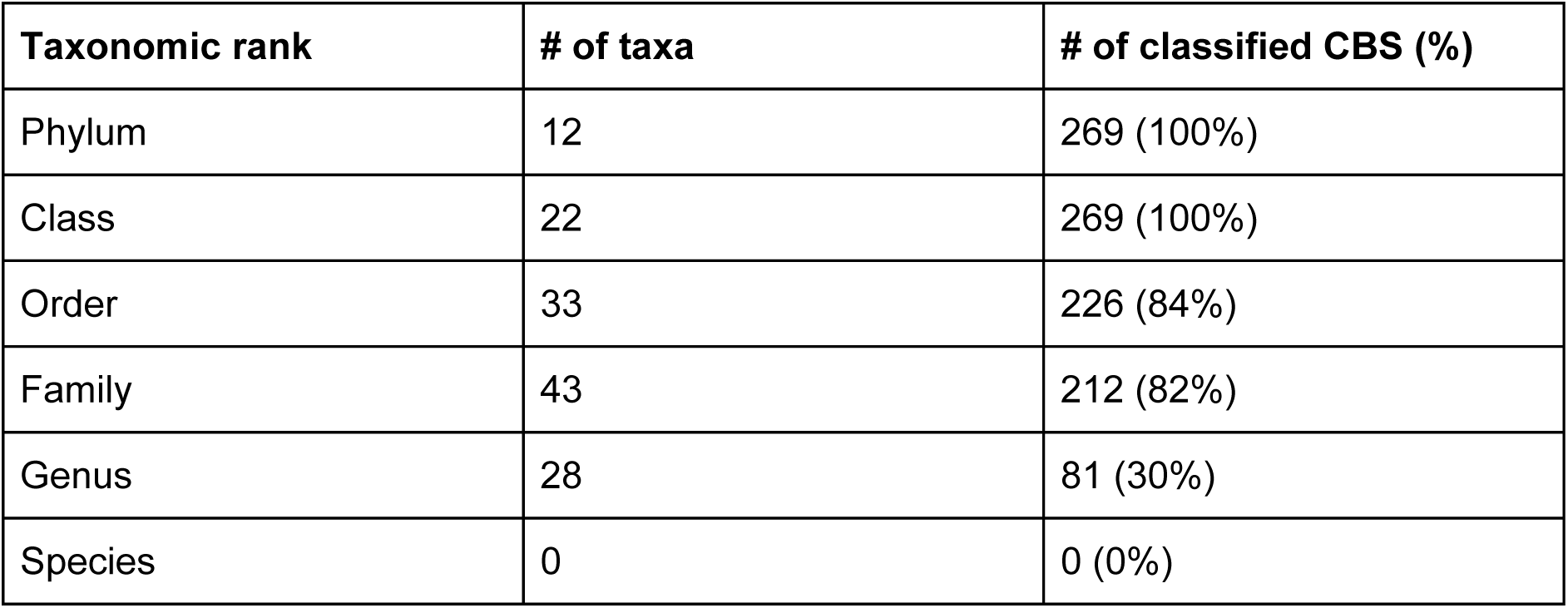
Number of identified taxa and classified CBS for each taxonomic rank. While 100% of the CBS could be assigned to a known phylum, only 81% were classified at the genus level and none at the species level.

| Taxonomic rank | # of taxa | # of classified CBS (%) |
| --- | --- | --- |
| Phylum | 12 | 269 (100%) |
| Class | 22 | 269 (100%) |
| Order | 33 | 226 (84%) |
| Family | 43 | 212 (82%) |
| Genus | 28 | 81 (30%) |
| Species | 0 | 0 (0%) |

### Antarctic cryptoendolithic communities share a core of ubiquitous species

We investigated the distribution of CBSs across the wide range of sampled conditions (see Methods and Supplementary Table 4), in order to identify a core of ubiquitous bacterial species throughout the Antarctic desert, or to assess if local environmental conditions and dispersal limitations selected site-specific species. With this purpose in mind, we considered a species present in a sample either: i) if an assembled genome assigned to the CBS was recovered from that sample; or ii) if the breadth of coverage of the mapped reads on the CBS representative is ≥50% and the ANI between the consensus sequence and the CBS is ≥ 0.95 (see Methods). The latter condition allowed us to reveal the presence of a CBS also in samples where its abundance was too low to allow assembly. We could define a set of 10 CBS that were present in at least 75% (14/18) of the samples (Fig. 1d, Supplementary Table 5). This core of conserved species was complemented by lineages, *i.e.* group of phylogenetically closely related CBS that were widely present in a large fraction of the samples. The core species were distributed in two phyla, (*Actinobacteria* and *Proteobacteria*) and two classes, i.e. *Actinobacteria* and *Alphaproteobacteria* (Supplementary Fig. 1). Amongst others, we found one CBS of the *Jiangellales*^*20*^ (Fig. 1h), an order from the class *Actinobacteria* that encompasses species isolated from different environments including indoor environments, cold springs on the Qinghai-Tibet Plateau^21^, cave^22^ and desert soils^23^. This CBS, that herein we named “*Candidatus Jiangella antarctica*”, was present across all samples (average percentage of mapped reads 1.92%, SD 1.93%), suggesting a high adaptation and specialization of this species to the harshest Antarctic environment. Moreover, extracting and classifying the near full-length 16S from the *Ca. Jiangella antarctica* (1513 bp), we did not found any significant match both in the Ribosomal Database Project^24^ (RDP, classified as “unclassified *Actinomycetales*”) and SILVA^25^ (identity of the best hit 92.09%), confirming that this species has not been previously reported. We also found three less ubiquitous species that were related to the Antarctic *Jiangella*. At the other end of the spectrum, we reported site-specific or near site-specific lineages; in particular, we found that, while all sample host at least one representative of the class *Chloroflexia*, with one CBS present in 67% of the samples, three samples (SS, TN, US) host the majority of CBS from this class, which in many cases are absent from all other samples (Fig. 1d, Supplementary Table 5).

### Antarctic bacteria form ancient monophyletic groups that evolved much earlier than the separation of Antarctica

We investigated if the reconstructed MAGs define new clades specific for Antarctica and if the estimated time of divergence of these clades from known lineages may indicate that Antarctic endoliths diversified and evolved in response to the glaciation of the Continent, started in the early Oligocene^26^ (~34 Ma), or, alternatively, if they were selected by the mutated environmental conditions in a pool of pre-existing species. To answer this question, for each bacterial order with at least 4 CBS (for a total of 19 orders, 377 MAGs and 200 CBS), we built a phylogenetic tree including both the MQ and HQ MAGs and reference genomes belonging to the same order from the GTDB database (see Methods). In order to generate homogeneously sized datasets, we selected sequences from the 19 order-specific datasets including all the Antarctic MAGs plus all their immediate reference sister taxon (as defined from the corresponding RAxML^27^ phylogenetic tree), plus reference representatives of other more distant clades distributed within the tree^28^. The size of the datasets ranged between 46 taxa in the *Solibacteriales* to 189 taxa for the *Corynebacteriales*, with most dataset comprising between 50 and 100 taxa. Using a molecular clock approach and available divergence estimates for calibrating the trees^28^, we inferred estimates for the age of the divergence of the Antarctica clades from the main tree within each bacterial order. Our phylogenetic and clock analyses indicated that the Antarctic MAGs (red branches in Fig. 2b,c, and Supplementary Data 1 are grouped into ancient monophyletic clades. In some cases, all Antarctic samples form a unique clade within a certain bacterial order, as in *Jiangellales, Microtrichales* and UBA5184, while in other cases we observed a large clustering of Antarctic MAGs interleaved by just one or two reference genomes as in *Thermomicrobiales, Solirubrobacteriales, Ktedonobacterales*, and *Isosphaerales*. In almost all other orders (e.g. *Acetobacterales, Acidobacteriales, Actinomycetales, Corynebacterales, Frankiales*), two or more unrelated Antarctic clades are revealed. Only in a few orders such as *Sphingomonadales* and *Actinomycetales*, Antarctic MAGs did not form distinct clades. Interestingly, no evidence of segregation of Antarctic lineages according to sampling site was found, suggesting that dispersal did not play a key role in determining the structure of these communities that is highly variable regardless of geography. Similar conclusions were reached by Archer and colleagues^11^, who recently reported that persistent local airborne inputs were unable to fully explain Antarctic soil community assembly. Our divergence estimates indicate that the vast majority of the Antarctic clades are extremely old (green and orange estimates in Fig. 2a). The diversification of the oldest Antarctic clades occurred on average circa 800 Ma, with estimates ranging from 1.2 billion to 410 Ma (Supplementary Table 6). While the oldest *Cyanobacteriales* and *Ktedonobacterales* Antarctic clades are Silurian to Devonian (before 410 Ma), the oldest Antarctic clades in all other orders are pre-Cambrian, with most of them originated in the Tonian (1000-720 Ma). Most of the oldest Antarctic clades may have therefore originated during the Tonian glaciations and before the many glaciations of the Cryogenian^29,30^, when Antarctica was part of the Supercontinent Rodinia.

**Fig. 2.**
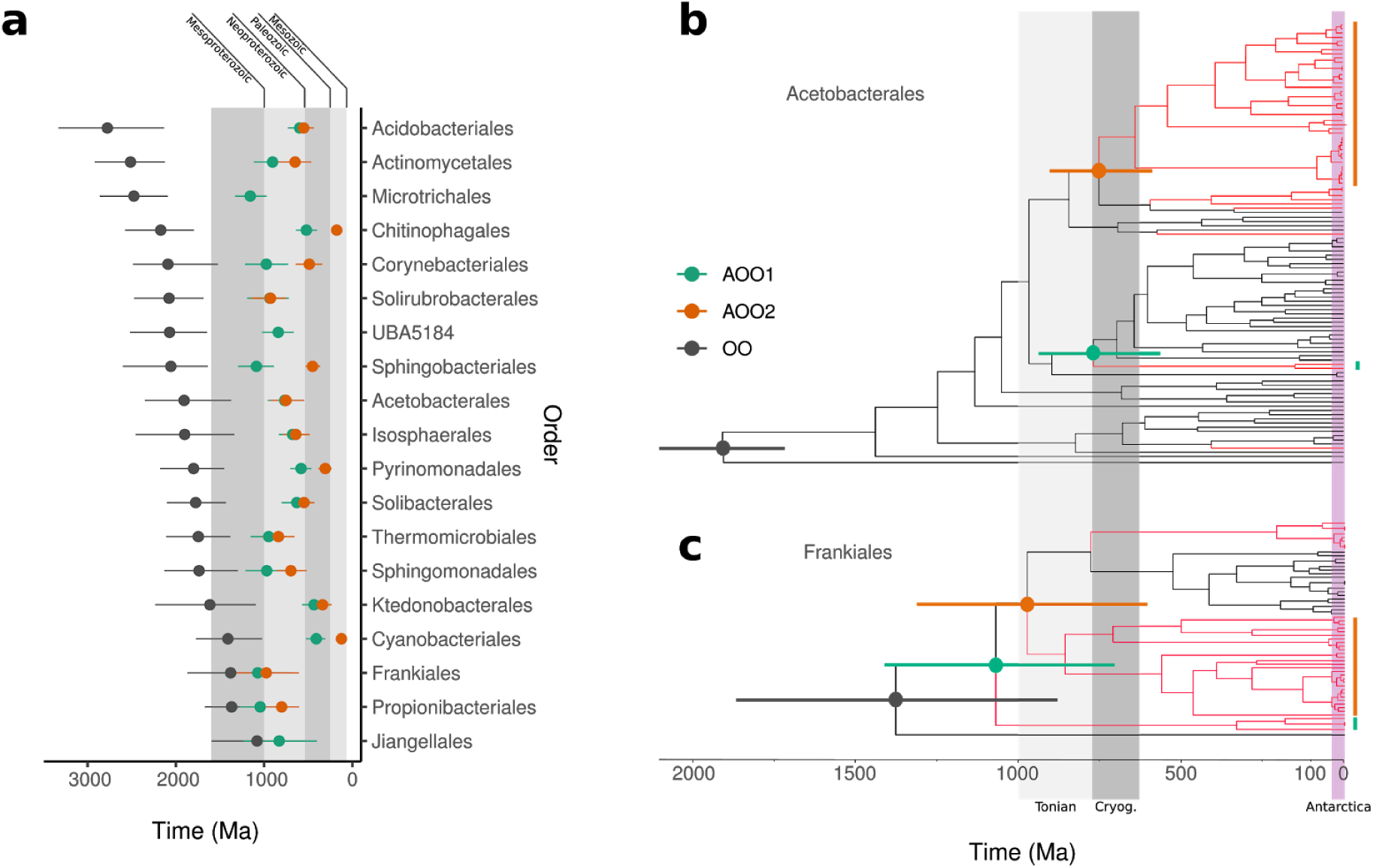
The neoproterozoic age of Antarctic cryptoendolithic clades. **a)** Summary of Bayesian divergence estimates indicate that most of the oldest lineages of Antarctic cryptoendoliths originated during the Neoproterozoic (1000-541 Ma). For each of the orders we show the origin of the order (OO: the split of the order from the closest order), the origin of the oldest uniquely Antarctic clade (AOO1: the split of the Antarctic calde from a non-Antarctic lineage of the same order), and, where present, the origin of the second oldest antarctic clade (AOO2) in each Order. **b)** Bayesian divergence tree of the *Acetobacterales* and **c)** of the *Frankiales* with the Antarctic clades highlighted in red. Time-trees for all other orders are in Supplementary Data 1. The shaded grey areas in a) identify geological eras, while those in b,c) identify the Tonian and Cryogenian periods of the Neoproterozoic era; pink area in 2b,c) identify a period compatible with glacial Antarctica.

### Antarctic species encode functions that distinguish them from known references, but that are not specific and common to all Antarctic MAGs

To understand if the Antarctic MAGs encode a common and specific set of functions that is responsible for their persistence in the Antarctic endolithic ecosystem, we characterized the functional potential of the HQ CBS representatives using EggNOG-mapper, and compared to the GTDB representative genomes from the same order. We found that the number of protein coding genes and the fraction of protein coding genes with homology to known protein families (Fig. 3a, upper panel and Supplementary Table 7) was usually similar to what was found for reference genomes. Comparing the functional profiles of Antarctic MAGs to reference genomes, we could not identify a set of functions that characterize the Antarctic MAGs across the whole dataset. Computing the Jaccard distance between the KEGG^31^ functional profiles we found that, in most cases, the pairwise distance between Antarctic MAGs was lower than between MAGs and GTDB representatives of the same order (Fig. 3b). At high taxonomic level and down to the order level, Antarctic MAGs tend to segregate together with reference genomes of the same taxonomy in a low dimensional projection (Fig. 3c). At the order level the distinction of Antarctic MAGs from reference genomes becomes in most cases apparent. These results confirm, using functional annotation, the results of the phylogenetic analysis that the CBSs form separate lineages within known orders.

**Fig. 3.**
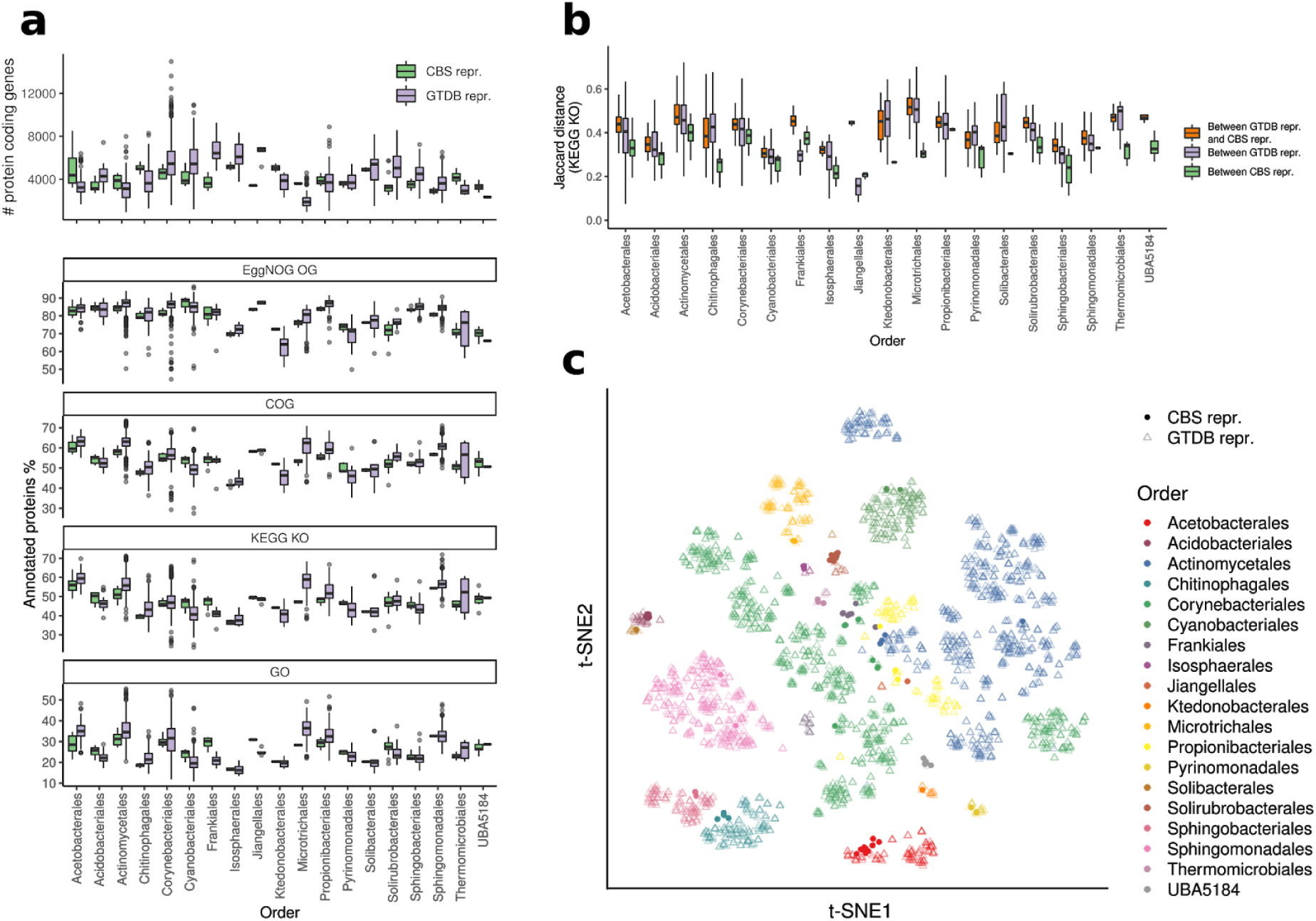
**a)** Upper panel: number of protein coding genes in the CBS representatives compared to the reference genomes from the same order; lower panels: fraction of protein coding genes with homologs in EggNOG OGs, COGs, KEGG KOs, and GO. **b)** Jaccard distance between the KEGG functional profiles for each Order. **c)** Starting from the Jaccard distances between the functional profiles, the t-SNE (t-distributed stochastic neighbor embedding) dimensionality reduction highlights the separation between genomes from different orders. Only HQ CBS representative genomes were considered in this analysis.

For instance, the Antarctic members of the family *Acetobacteraceae* (order *Acetobacterales*) form a distinct phylogenetic clade (midpoint-rooted maximum-likelihood phylogenetic tree in Fig. 4a) and encode a set of metabolic functions that distinguishes them from the other members of the same family (Fig. 4b). Amongst other high-level functional categories, we found that the Antarctic *Acetobacteraceae* were enriched in genes related to transport and metabolism, in particular of carbohydrates (Fig. 4c). Another notable example is the Antarctic *Frankiales*, that form two groups that are clearly distinct from all known species of the same order (Supplementary Fig. 2 and Supplementary Data 1).

**Fig. 4.**
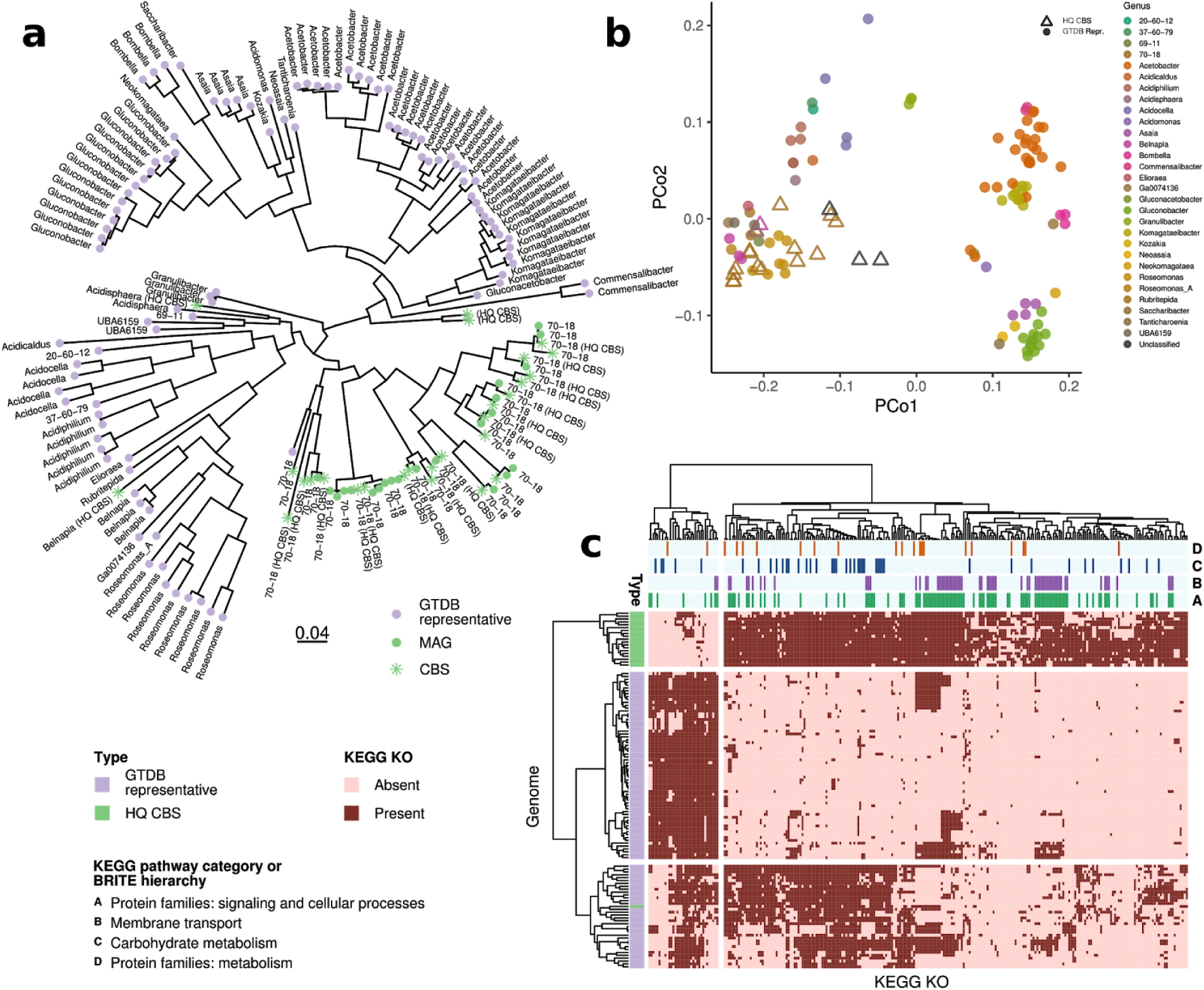
Antarctic CBS form a distinct clade in the order *Acetobacterales*, family *Acetobacteraceae*, with characteristic metabolic potential. **a)** Maximum-likelihood phylogenetic tree based on GTDB 120 core genes including representative genomes (violet) and Antarctic CBS (green). **b)** Principal Coordinate Analysis (Jaccard distance) of the metabolic potential of the GTDB representatives (circles) and Antarctic CBS (triangles). **c)** The Fisher’s exact test (Bonferroni corrected p<0.01) highlights enriched functional categories. Genomes and KEGG orthologs are clustered according to the Hamming distance between the profiles. The top four KEGG categories significantly more present in the Antarctic CBS are highlighted in the upper bars.

## Discussion

In conclusion, our data report for the first time the genomes of the dominant bacterial species in Antarctic cryptoendolithic communities, substantially expanding our knowledge of several important taxa. Most of these new species are organized into ancient monophyletic clades that differentiated from known bacterial orders in a time range that predates the estimated origin of the modern Antarctica and the establishment of glacial climate conditions. Although our inferences do not allow a precise discussion of the exact timing of the origin of Antarctic clades, they clearly show that diversification was not in response to recent glaciation and formation of the Antarctic climate. Instead, all our data point toward a scenario where extant Antarctic bacterial clades are the remnants of ancient cold-tolerant bacterial lineages which seem now confined to Antarctica where they may have found ecological refugia. While some of these new species are specific to a small number of samples, there are several that are present in all samples, despite the fact that these were collected from geographically distant sites at altitudes ranging from 834 to 3100 m a.s.l. This considerable level of homogeneity of the composition of these samples suggests that the endolithic localization provides a microclimate that is remarkably insensitive to different external conditions. On the other hand, the ubiquitous distribution of several species is consistent with a model in which the endolithic niche was colonized by preexisting organisms that sought refuge in this sheltered niche to survive changing environmental conditions. Given the extremely long timescales over which the evolution of these organisms occurred, it remains to be seen how these communities will react to fast occurring climate change. These findings suggest a new scenario for the search of life on Mars where microbial life, if ever evolved in ancient times, may have escaped extinction finding refugia and evolving from ancient lineages under the present conditions. These genomic resources represent only the tip of the iceberg and will underpin future studies on the structure and function of these ecosystems at the edge of life and their evolution in relation to the geological history of the continent.

## Methods

### Sampling area

Victoria Land is a region of Continental Antarctica which fronts the western side of the Ross Sea and the Ross Ice Shelf; this land, positioned between the Polar Plateau and the coast and exposed to a wide spectrum of climatic extremes, including temperature and precipitation regimes, covers a latitudinal gradient of 8° from Darwin Glacier (78°00’) to Cape Adare (70°30’S)^32^. Ice-free areas dominate the landscape of Southern Victoria Land and the high-altitude locations of Northern Victoria Land, while low-elevation coastal soils of Northern Victoria Land see considerable marine and biological influence.

Sandstone rocks were collected by L. Selbmann in Victoria Land along a latitudinal transect ranging from 74°10’44.0”S 162°30’53.0”E (Mt. New Zealand, Northern Victoria Land) to 77°52’28.6”S 160°44’22.6”E (University Valley, Southern Victoria Land) during the XXXI Italian Antarctic Expedition (Dec. 2015-Jan. 2016). Samples were collected at different conditions namely sun exposure and an altitudinal transect, from 834 to 3100 m a.s.l. to provide a comprehensive overview of endolithic diversity (Fig. 1a,b,c). Rocks were excised using a geologic hammer and sterile chisel, and rock samples, preserved in sterile plastic bags, transported and stored at -20°C in the Culture Collection from Extreme Environments (CCFEE), Mycological Section on the Italian Antarctic National Museum, MNA, at the Tuscia University (Viterbo, Italy), until downstream analysis.

### DNA extraction, library preparation and sequencing

DNA was extracted from three samples for each site and then pooled. Metagenomic DNA was extracted from 1 g of crushed rocks using MoBio Powersoil kit (MOBIO Laboratories, Carlsbad, CA, USA). The quality of the DNA extracted was determined by electrophoresis using a 1.5 % agarose gel and with a spectrophotometer (VWR International) and quantified using the Qubit dsDNA HS Assay Kit (Life Technologies, USA).

Shotgun metagenomic libraries were prepared and sequenced at the DOE Joint Genome Institute (JGI) as a part of a Community Science Project (Pi: Laura Selbmann; co-PI: Jason E. Stajich) at JGI^16^. Paired-end sequencing libraries were constructed and sequenced as 2×151 bp using the Illumina NovaSeq platform (Illumina Inc, San Diego, CA).

### Sequencing reads preparation and assembly

BBDuk (http://sourceforge.net/projects/bbmap/) v38.25 was used to remove contaminants, trim adapters and low-quality sequences. The procedure removed reads that contained 4 or more ‘N’ bases, had an average quality score across the read less than 3 or had a minimum length <= 51 bp or 33% of the full read length. Filtered and trimmed paired-end reads were error corrected using BFC^33^ r181 with parameters `-1 -s 10g -k 21 -t 10` and orphan reads were removed. Assembly was performed with SPAdes^34^ 3.12.0 using the parameters `-m 2000 -o spades3 --only-assembler -k 33,55,77,99,127 --meta -t 32`.

### Binning

Metagenomic contigs were binned into candidate genomes using MetaBAT2^35^ (Metagenome Binning based on Abundance and Tetranucleotide frequency) version 2.12.1. Briefly, high quality reads were mapped on assembled contigs using Bowtie2^36^ 2.3.4.3. Samtools^37^ 1.3.1 (htslib 1.3.2) was used to create and sort the BAM files (`.bam`). The depth of coverage was estimated applying the jgi_summarize_bam_contig_depths tool. Contigs sequences and the depth of coverage estimates were used by MetaBAT2 to recover the candidate MAGs.

### Quality assessment and dereplication

Completeness and contamination estimates of bacterial and archaeal MAGs were obtained by CheckM^38^. According to recent guidelines^39^, MAGs were classified into “high-quality draft” (HQ) with >90% completeness and <5% contamination and “medium-quality draft” (MQ) with completeness estimates of ≥50% and less than 10% contamination. Candidate bacterial species (CBS) were identified by clustering HQ and MQ MAGs at species level^40^ (>95% Average Nucleotide Identity - ANI) using dRep^41^ v2.0.0. For each CBS, the MAG with the highest quality score has been chosen as representative.

### Taxonomic classification

MQ and HQ MAGs were taxonomically classified using the genome taxonomy database toolkit^18^ (GTDB-Tk) v0.1.6 and the GTDB release 86, following the recently proposed nomenclature of Prokaryotes^42,43^. GTDB-Tk classifies a query genome combining its placement in the GTDB^18^ reference tree (release 86 includes a total of 21.263 genomes in the tree), its RED^18^, and its ANI to reference genomes. Approximately-maximum-likelihood phylogenetic tree from the GTDB protein alignments of the 269 CBS representatives (Fig. 1), and of the orders *Acetobacterales* (Fig. 4) and *Frankiales* (Supplementary Fig. 2) were inferred using FastTree^44^ 2.1.10 (WAG+CAT model, options `-wag -gamma`) and rooted at midpoint.

### Percentage of mapping reads and CBS presence estimation

For each metagenomic sample, high quality reads were aligned against each CBS representative using Bowtie2 2.3.4.3 using the parameter --no-unal. Samtools 1.3.1 (htslib 1.3.2) was used to create and index the BAM files (.bam). The depth of coverage, the breadth *B*_*n*_ (i.e. the fraction of bases of the CBS representative genome that are covered with depth *n*) and the number of mapped reads were calculated on the BAM file using pysam (https://github.com/pysam-developers/pysam) 0.15.2 and Python 3.5.3. The fraction of reads mapping on a CBS representative was computed as the number of successfully aligned reads normalized by the total number of reads aligning the entire set of the CBS representatives. Regions with no coverage were identified using BEDtools^45^ 2.26.0 with the options `-bga -split`. Variant calling was performed with `samtools mpileup` and bcftools call^46^ (v1.3.1, options `--ploidy 1 -mv`). Tabix^47^ 1.3.2 was used to index the output VCF file. The consensus sequence was generated using the command `bcftools consensus` masking the zero coverage regions previously identified. The ANI between the consensus sequence and the CBS representative (ANI_CBS_) was estimated using fastANI^40,46^ v1.1. Finally, a CBS was tagged as present in a sample if the breadth of coverage (at depth 2) *B*_*2*_ was ≥ 0.5 and ANI_CBS_ ≥ 95%.

### Divergence estimates

Divergence times were independently estimated on orders containing at least 4 CBS, for a total of 19 analyzed orders. For each order, we built a protein MSA using the 120 GTDB bacterial marker genes including: i) 32 reference sequences from outgroups outside the order; ii) the GTDB representatives; iii) the MQ and HQ Antarctic MAGs, iv) a set of outgroup in order to reconstruct the first radiation within Eubacteria as in^28^ and using it as a calibration point. The 19 datasets were calibrated with this same prior. We calibrated the crown (divergence) of Eubacteria using a prior on the root of 3453 million years ago (Ma) and a standard deviation of 60 Ma (values kindly provided by Davide Pisani) and corresponding to the posterior estimate for the crown of the Eubacteria^28^. Since our taxon sampling replicates the taxon sampling^28^, we could safely apply the previous estimate for the crown of the Eubacteria to our root (which coincides with the crown of Eubacteria, we did not use Archaea or Eukaryotes outgroups). Markov chain Monte Carlo (MCMC) analyses were performed using Beast^48^ 1.10 for 100 million generations sampling every 1000 generations. Convergence was assessed by using the Effective Sample Sizes (ESS) estimated by Tracer^49^ v1.7.1 on posteriors and log-likelihood. In order to maximize the ESS statistics, a burn-in ranging from 50% to 80% of the simulation was used. For computational reasons, we performed model selection using only one dataset (Acidobacteriales) as representative. We compared a relaxed clock (log-normal) versus the strict clock, and a coalescence (constant) versus a speciation (birth-death) demographic model. The most fitting combination of priors (relaxed clock plus coalescence) was found using path sampling and AICM. Amino-acid substitutions were modelled using the LG matrix with amino acid frequencies inferred from the data; among-site rate variation was modelled using a gamma distribution with four discrete categories. All Bayesian consensus trees are in Supplementary Data 1. For each order, the mean age (plus the 95% High Posterior Densities heights) for the first split of a uniquely Antarctic group (green node) from the known reference sequence from that particular order was plotted. In the case of more than one monophyletic Antarctic group, the age of the second oldest Antarctic group (orange node) was also shown.

### Functional annotation

Functional annotation was performed only on HQ CBS representatives of orders containing at least 4 CBS (for a total of 19 orders analyzed). In order to avoid systematic effects due to different annotation methods, both HQ MAGs and GTDB representative genomes were processed as follows: i) translated coding DNA sequences (CDSs) were predicted using Prokka^50^ 1.13.4 which wraps the software Prodigal^51^. ii) the CDSs were functionally annotated using EggNOG-mapper^52^ (option `--database bact`) and the eggNOG Orthologous Groups (OGs) database^53^ 4.5.1. The EggNOG database integrates functional annotations collected from several sources, including KEGG functional orthologs^54^, COG categories^55^ and Gene Ontology (GO) terms.

### Statistical analysis

Downstream analysis was performed using the R environment (https://www.R-project.org/) version 3.6.1. T-SNE dimensionality reduction was carried out using the R package “tsne” (https://CRAN.R-project.org/package=tsne) version 0.1-3 and the PCoA (Principal Coordinate Analysis) using the function “pcoa()” (default parameters) available in the R library “ape”^56^ version 5.3. Fisher’s exact tests were conducted using the function “fisher.test()” (default parameters) available in the R package “stats” version 3.6.1.

### Data availability

Raw metagenomes reads and assemblies are deposited under the NCBI accession numbers listed in Supplementary Table 4. Metagenome assemblies, gene predictions, and JGI annotations are available in the IMG/M web site (https://img.jgi.doe.gov/) and in the zenodo repository (https://zenodo.org/record/3610489; DOI: 10.5281/zenodo.3610489). MAGs, translated coding sequences and annotations for high quality MAGs, *Candidatus Jiangella antarctica* ribosomal rRNA genes and metadata are available at the zenodo repository (DOI: 10.5281/zenodo.3671353).

## Author contributions

L.S. and J.E.S. are PI and co-PI of the “Metagenomic reconstruction of endolithic communities from Victoria Land, Antarctica” Joint Genome Institute Community Sequencing Project 503708. S.T. is the Leader of the Metagenomic Group. Samples were collected by L.S. during the XXXI Italian Antarctic Expedition (2015-2016); C.C., L.S., J.E.S, D.A. and C.D. designed the research; C.C. performed DNA extraction and quality check control; D.A., C.C., O.R.S. and C.D. analyzed the data; C.C., L.S.,D.A., J.E.S. and C.D. wrote the paper with input from O.R.S., S.O., and S.T.

## Acknowledgements

L.S. and C.C. wish to thank the Italian National Program for Antarctic Research for funding sampling campaigns and research activities in Italy in the frame of PNRA projects. The Italian Antarctic National Museum (MNA) is kindly acknowledged for financial support to the Mycological Section of the MNA and for providing rock samples used in this study stored in the Culture Collection of Fungi from Extreme Environments (CCFEE), University of Tuscia, Italy. The work conducted by the U.S. Department of Energy (DOE) Joint Genome Institute, a DOE Office of Science User Facility, was supported by the Office of Science of the DOE under Contract No. DE-AC02-05CH11231. We wish to thank Davide Pisani for kindly providing mean posterior estimates for calibration purpose. J. Stajich is a CIFAR fellow in the program Fungal Kingdom: Threats and Opportunities.

