## Supplementary Data 1 for "Antarctic cryptoendolithic bacterial lineages of pre-Cambrian origin as proxy for Mars colonization"

Bayesian posterior consensus trees of each of the 19 bacterial order datasets. For all nodes we provide the mean estimate (in million of years) and the 95% High Posterior densities of estimates (as bars). Uniquely Antarctic clades or lineages are  depicted in red. For each of the orders we show the origin of the Order (OO: the split of the order from the closest order) with a black circle; the  origin of the oldest uniquely Antarctic clade or lineage with a green circle (AOO1: the split of the Antarctic clade from a non-Antarctic lineage of the same order); where present, the origin of the second oldest Antarctic clade (AOO2) with a orange circle, and the origin of other eventual only Antarctic clades or lineages (A00n) with grey circle.

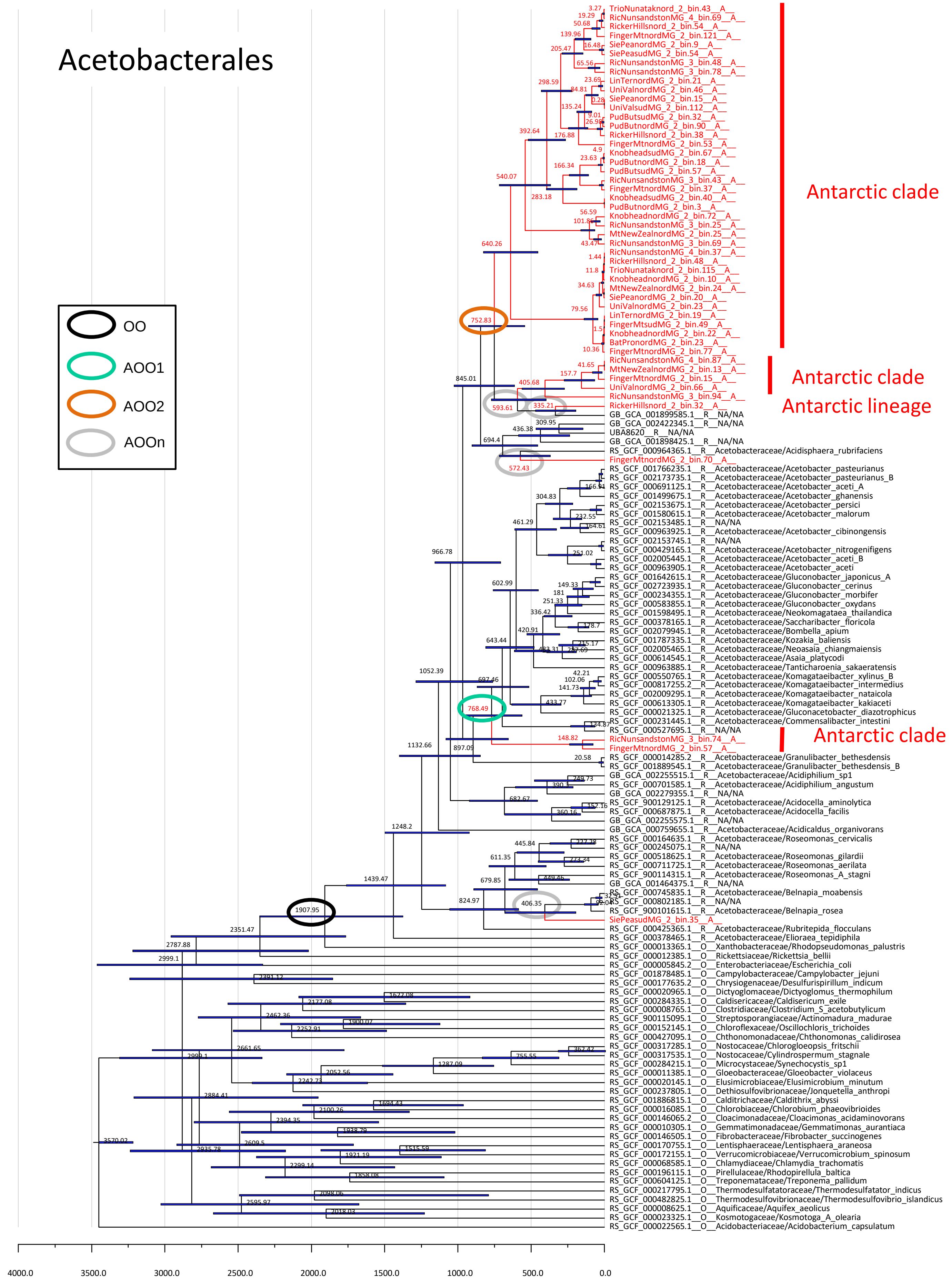
Acetobacterales
OO
AOO1
AOO2
AOOn
Antarctic clade
Antarctic clade
Antarctic lineage
Antarctic clade
TrioNunataknord_2_bin.43__A__
RicNunsandstonMG_4_bin.69__A__
RickerHillsnord_2_bin.54__A__
FingerMtnordMG_2_bin.121__A__
SiePeanordMG_2_bin.9__A__
SiePeasudMG_2_bin.54__A__
RicNunsandstonMG_3_bin.48__A__
RicNunsandstonMG_3_bin.78__A__
LinTernordMG_2_bin.21__A__
UniValnordMG_2_bin.46__A__
SiePeanordMG_2_bin.15__A__
UniValsudMG_2_bin.112__A__
PudButsudMG_2_bin.32__A__
PudButnordMG_2_bin.90__A__
RickerHillsnord_2_bin.38__A__
FingerMtnordMG_2_bin.53__A__
KnobheadsudMG_2_bin.67__A__
PudButnordMG_2_bin.18__A__
PudButsudMG_2_bin.57__A__
RicNunsandstonMG_3_bin.43__A__
FingerMtnordMG_2_bin.37__A__
KnobheadsudMG_2_bin.40__A__
PudButnordMG_2_bin.3__A__
KnobheadnordMG_2_bin.72__A__
RicNunsandstonMG_3_bin.25__A__
MtNewZealnordMG_2_bin.25__A__
RicNunsandstonMG_3_bin.69__A__
RicNunsandstonMG_4_bin.37__A__
RickerHillsnord_2_bin.48__A__
TrioNunataknord_2_bin.115__A__
KnobheadnordMG_2_bin.10__A__
MtNewZealnordMG_2_bin.24__A__
SiePeanordMG_2_bin.20__A__
UniValnordMG_2_bin.23__A__
LinTernordMG_2_bin.19__A__
FingerMtsudMG_2_bin.49__A__
KnobheadnordMG_2_bin.22__A__
BatPronordMG_2_bin.23__A__
FingerMtnordMG_2_bin.77__A__
RicNunsandstonMG_4_bin.87__A__
MtNewZealnordMG_2_bin.13__A__
FingerMtnordMG_2_bin.15__A__
UniValnordMG_2_bin.66__A__
RicNunsandstonMG_3_bin.94__A__
RickerHillsnord_2_bin.32__A__
GB_GCA_001899585.1__R__NA/NA
GB_GCA_002422345.1__R__NA/NA
UBA8620__R__NA/NA
GB_GCA_001898425.1__R__NA/NA
RS_GCF_000964365.1__R__Acetobacteraceae/Acidisphaera_rubrifaciens
FingerMtnordMG_2_bin.70__A__
RS_GCF_001766235.1__R__Acetobacteraceae/Acetobacter_pasteurianus
RS_GCF_002173735.1__R__Acetobacteraceae/Acetobacter_pasteurianus_B
RS_GCF_000691125.1__R__Acetobacteraceae/Acetobacter_aceti_A
RS_GCF_001499675.1__R__Acetobacteraceae/Acetobacter_ghanensis
RS_GCF_002153675.1__R__Acetobacteraceae/Acetobacter_persici
RS_GCF_001580615.1__R__Acetobacteraceae/Acetobacter_malorum
RS_GCF_002153485.1__R__NA/NA
RS_GCF_000963925.1__R__Acetobacteraceae/Acetobacter_cibinongensis
RS_GCF_002153745.1__R__NA/NA
RS_GCF_000429165.1__R__Acetobacteraceae/Acetobacter_nitrogenifigens
RS_GCF_002005445.1__R__Acetobacteraceae/Acetobacter_aceti_B
RS_GCF_000963905.1__R__Acetobacteraceae/Acetobacter_aceti
RS_GCF_001642615.1__R__Acetobacteraceae/Gluconobacter_japonicus_A
RS_GCF_002723935.1__R__Acetobacteraceae/Gluconobacter_cerinus
RS_GCF_000234355.1__R__Acetobacteraceae/Gluconobacter_morbifer
RS_GCF_000583855.1__R__Acetobacteraceae/Gluconobacter_oxydans
RS_GCF_001598495.1__R__Acetobacteraceae/Neokomagataea_thailandica
RS_GCF_000378165.1__R__Acetobacteraceae/Saccharibacter_floricola
RS_GCF_002079945.1__R__Acetobacteraceae/Bombella_apium
RS_GCF_001787335.1__R__Acetobacteraceae/Kozakia_baliensis
RS_GCF_002005465.1__R__Acetobacteraceae/Neoasaia_chiangmaiensis
RS_GCF_000614545.1__R__Acetobacteraceae/Asaia_platycodi
RS_GCF_000963885.1__R__Acetobacteraceae/Tanticharoenia_sakaeratensis
RS_GCF_000550765.1__R__Acetobacteraceae/Komagataeibacter_xylinus_B
RS_GCF_000817255.2__R__Acetobacteraceae/Komagataeibacter_intermedius
RS_GCF_002009295.1__R__Acetobacteraceae/Komagataeibacter_nataicola
RS_GCF_000613305.1__R__Acetobacteraceae/Komagataeibacter_kakiaceti
RS_GCF_000021325.1__R__Acetobacteraceae/Gluconacetobacter_diazotrophicus
RS_GCF_000231445.1__R__Acetobacteraceae/Commensalibacter_intestini
RS_GCF_000527695.1__R__NA/NA
RicNunsandstonMG_3_bin.74__A__
FingerMtnordMG_2_bin.57__A__
RS_GCF_000014285.2__R__Acetobacteraceae/Granulibacter_bethesdensis
RS_GCF_001889545.1__R__Acetobacteraceae/Granulibacter_bethesdensis_B
GB_GCA_002255515.1__R__Acetobacteraceae/Acidiphilium_sp1
RS_GCF_000701585.1__R__Acetobacteraceae/Acidiphilium_angustum
GB_GCA_002279355.1__R__NA/NA
RS_GCF_900129125.1__R__Acetobacteraceae/Acidocella_aminolytica
RS_GCF_000687875.1__R__Acetobacteraceae/Acidocella_facilis
GB_GCA_002255575.1__R__NA/NA
GB_GCA_000759655.1__R__Acetobacteraceae/Acidicaldus_organivorans
RS_GCF_000164635.1__R__Acetobacteraceae/Roseomonas_cervicalis
RS_GCF_000245075.1__R__NA/NA
RS_GCF_000518625.1__R__Acetobacteraceae/Roseomonas_gilardii
RS_GCF_000711725.1__R__Acetobacteraceae/Roseomonas_aerilata
RS_GCF_900114315.1__R__Acetobacteraceae/Roseomonas_A_stagni
GB_GCA_001464375.1__R__NA/NA
RS_GCF_000745835.1__R__Acetobacteraceae/Belnapia_moabensis
RS_GCF_000802185.1__R__NA/NA
RS_GCF_900101615.1__R__Acetobacteraceae/Belnapia_rosea
SiePeasudMG_2_bin.35__A__
RS_GCF_000425365.1__R__Acetobacteraceae/Rubritepida_flocculans
RS_GCF_000378465.1__R__Acetobacteraceae/Elioraea_tepidiphila
RS_GCF_000013365.1__O__Xanthobacteraceae/Rhodopseudomonas_palustris
RS_GCF_000012385.1__O__Rickettsiaceae/Rickettsia_bellii
RS_GCF_000005845.2__O__Enterobacteriaceae/Escherichia_coli
RS_GCF_001878485.1__O__Campylobacteraceae/Campylobacter_jejuni
RS_GCF_000177635.2__O__Chrysiogenaceae/Desulfurispirillum_indicum
RS_GCF_000020965.1__O__Dictyoglomaceae/Dictyoglomus_thermophilum
RS_GCF_000284335.1__O__Caldisericaceae/Caldisericum_exile
RS_GCF_000008765.1__O__Clostridiaceae/Clostridium_S_acetobutylicum
RS_GCF_900115095.1__O__Streptosporangiaceae/Actinomadura_madurae
RS_GCF_000152145.1__O__Chloroflexaceae/Oscillochloris_trichoides
RS_GCF_000427095.1__O__Chthonomonadaceae/Chthonomonas_calidirosea
RS_GCF_000317285.1__O__Nostocaceae/Chlorogloeopsis_fritschii
RS_GCF_000317535.1__O__Nostocaceae/Cylindrospermum_stagnale
RS_GCF_000284215.1__O__Microcystaceae/Synechocystis_sp1
RS_GCF_000011385.1__O__Gloeobacteraceae/Gloeobacter_violaceus
RS_GCF_000020145.1__O__Elusimicrobiaceae/Elusimicrobium_minutum
RS_GCF_000237805.1__O__Dethiosulfovibrionaceae/Jonquetella_anthropi
RS_GCF_001886815.1__O__Calditrichaceae/Caldithrix_abyssi
RS_GCF_000016085.1__O__Chlorobiaceae/Chlorobium_phaeovibrioides
RS_GCF_000146065.2__O__Cloacimonadaceae/Cloacimonas_acidaminovorans
RS_GCF_000010305.1__O__Gemmatimonadaceae/Gemmatimonas_aurantiaca
RS_GCF_000146505.1__O__Fibrobacteraceae/Fibrobacter_succinogenes
RS_GCF_000170755.1__O__Lentisphaeraceae/Lentisphaera_araneosa
RS_GCF_000172155.1__O__Verrucomicrobiaceae/Verrucomicrobium_spinosum
RS_GCF_000068585.1__O__Chlamydiaceae/Chlamydia_trachomatis
RS_GCF_000196115.1__O__Pirellulaceae/Rhodopirellula_baltica
RS_GCF_000604125.1__O__Treponemataceae/Treponema_pallidum
RS_GCF_000217795.1__O__Thermodesulfatatoraceae/Thermodesulfatator_indicus
RS_GCF_000482825.1__O__Thermodesulfovibrionaceae/Thermodesulfovibrio_islandicus
RS_GCF_000008625.1__O__Aquificaceae/Aquifex_aeolicus
RS_GCF_000023325.1__O__Kosmotogaceae/Kosmotoga_A_olearia
RS_GCF_000022565.1__O__Acidobacteriaceae/Acidobacterium_capsulatum
4000.0
3500.0
3000.0
2500.0
2000.0
1500.0
1000.0
500.0
0.0

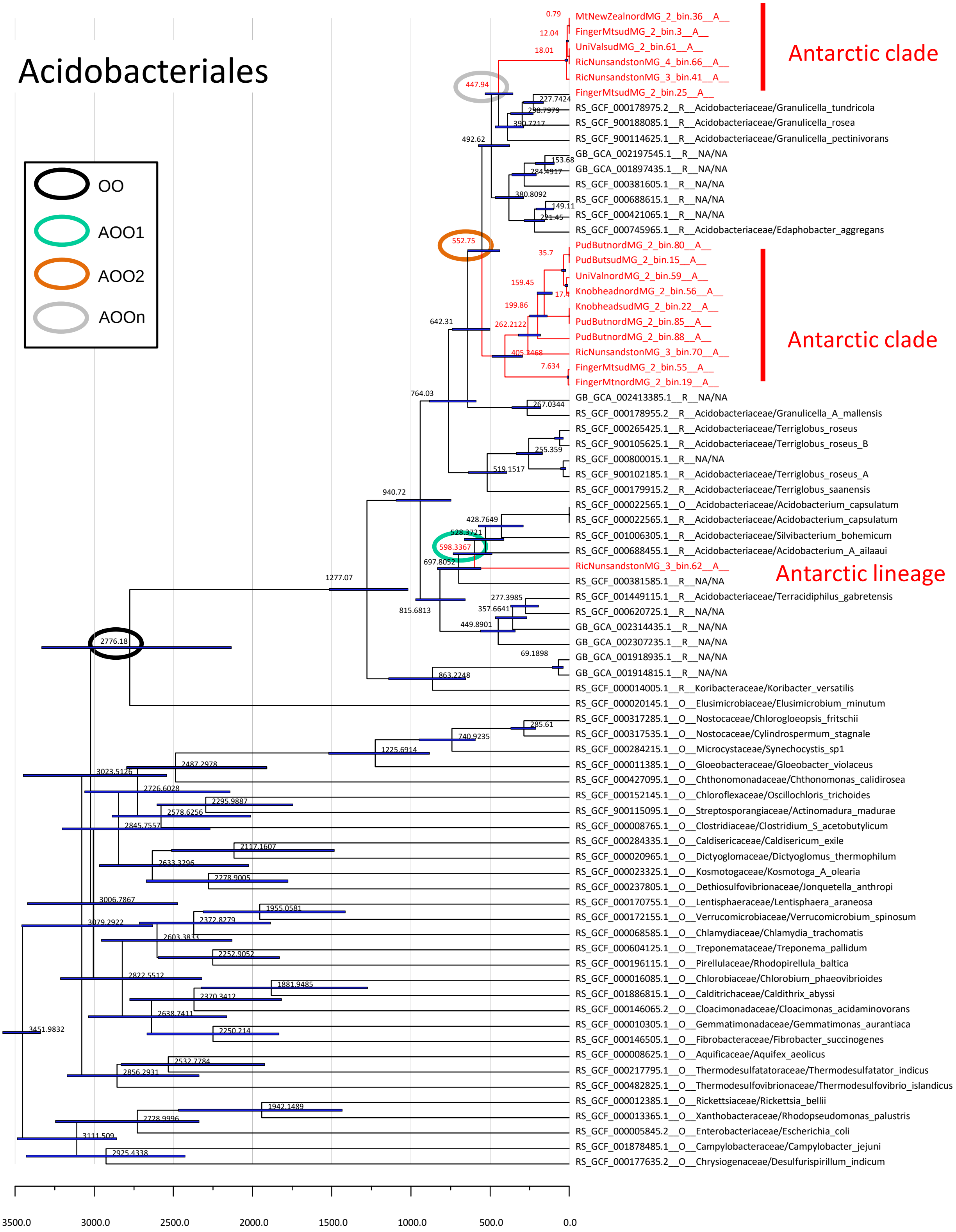

Acidobacteriales
OO
AOO1
AOO2
AOOn
Antarctic clade
Antarctic clade
Antarctic lineage
MtNewZealnordMG_2_bin.36__A__
FingerMtsudMG_2_bin.3__A__
UniValsudMG_2_bin.61__A__
RicNunsandstonMG_4_bin.66__A__
RicNunsandstonMG_3_bin.41__A__
FingerMtsudMG_2_bin.25__A__
RS_GCF_000178975.2__R__Acidobacteriaceae/Granulicella_tundricola
RS_GCF_900188085.1__R__Acidobacteriaceae/Granulicella_rosea
RS_GCF_900114625.1__R__Acidobacteriaceae/Granulicella_pectinivorans
GB_GCA_002197545.1__R__NA/NA
GB_GCA_001897435.1__R__NA/NA
RS_GCF_000381605.1__R__NA/NA
RS_GCF_000688615.1__R__NA/NA
RS_GCF_000421065.1__R__NA/NA
RS_GCF_000745965.1__R__Acidobacteriaceae/Edaphobacter_aggregans
PudButnordMG_2_bin.80__A__
PudButsudMG_2_bin.15__A__
UniValnordMG_2_bin.59__A__
KnobheadnordMG_2_bin.56__A__
KnobheadsudMG_2_bin.22__A__
PudButnordMG_2_bin.85__A__
PudButnordMG_2_bin.88__A__
RicNunsandstonMG_3_bin.70__A__
FingerMtsudMG_2_bin.55__A__
FingerMtnordMG_2_bin.19__A__
GB_GCA_002413385.1__R__NA/NA
RS_GCF_000178955.2__R__Acidobacteriaceae/Granulicella_A_mallensis
RS_GCF_000265425.1__R__Acidobacteriaceae/Terriglobus_roseus
RS_GCF_900105625.1__R__Acidobacteriaceae/Terriglobus_roseus_B
RS_GCF_000800015.1__R__NA/NA
RS_GCF_900102185.1__R__Acidobacteriaceae/Terriglobus_roseus_A
RS_GCF_000179915.2__R__Acidobacteriaceae/Terriglobus_saanensis
RS_GCF_000022565.1__O__Acidobacteriaceae/Acidobacterium_capsulatum
RS_GCF_000022565.1__R__Acidobacteriaceae/Acidobacterium_capsulatum
RS_GCF_001006305.1__R__Acidobacteriaceae/Silvibacterium_bohemicum
RS_GCF_000688455.1__R__Acidobacteriaceae/Acidobacterium_A_ailaaui
RicNunsandstonMG_3_bin.62__A__
RS_GCF_000381585.1__R__NA/NA
RS_GCF_001449115.1__R__Acidobacteriaceae/Terracidiphilus_gabretensis
RS_GCF_000620725.1__R__NA/NA
GB_GCA_002314435.1__R__NA/NA
GB_GCA_002307235.1__R__NA/NA
GB_GCA_001918935.1__R__NA/NA
GB_GCA_001914815.1__R__NA/NA
RS_GCF_000014005.1__R__Koribacteraceae/Koribacter_versatilis
RS_GCF_000020145.1__O__Elusimicrobiaceae/Elusimicrobium_minutum
RS_GCF_000317285.1__O__Nostocaceae/Chlorogloeopsis_fritschii
RS_GCF_000317535.1__O__Nostocaceae/Cylindrospermum_stagnale
RS_GCF_000284215.1__O__Microcystaceae/Synechocystis_sp1
RS_GCF_000011385.1__O__Gloeobacteraceae/Gloeobacter_violaceus
RS_GCF_000427095.1__O__Chthonomonadaceae/Chthonomonas_calidirosea
RS_GCF_000152145.1__O__Chloroflexaceae/Oscillochloris_trichoides
RS_GCF_900115095.1__O__Streptosporangiaceae/Actinomadura_madurae
RS_GCF_000008765.1__O__Clostridiaceae/Clostridium_S_acetobutylicum
RS_GCF_000284335.1__O__Caldisericaceae/Caldisericum_exile
RS_GCF_000020965.1__O__Dictyoglomaceae/Dictyoglomus_thermophilum
RS_GCF_000023325.1__O__Kosmotogaceae/Kosmotoga_A_olearia
RS_GCF_000237805.1__O__Dethiosulfovibrionaceae/Jonquetella_anthropi
RS_GCF_000170755.1__O__Lentisphaeraceae/Lentisphaera_araneosa
RS_GCF_000172155.1__O__Verrucomicrobiaceae/Verrucomicrobium_spinosum
RS_GCF_000068585.1__O__Chlamydiaceae/Chlamydia_trachomatis
RS_GCF_000604125.1__O__Treponemataceae/Treponema_pallidum
RS_GCF_000196115.1__O__Pirellulaceae/Rhodopirellula_baltica
RS_GCF_000016085.1__O__Chlorobiaceae/Chlorobium_phaeovibrioides
RS_GCF_001886815.1__O__Calditrichaceae/Caldithrix_abyssi
RS_GCF_000146065.2__O__Cloacimonadaceae/Cloacimonas_acidaminovorans
RS_GCF_000010305.1__O__Gemmatimonadaceae/Gemmatimonas_aurantiaca
RS_GCF_000146505.1__O__Fibrobacteraceae/Fibrobacter_succinogenes
RS_GCF_000008625.1__O__Aquificaceae/Aquifex_aeolicus
RS_GCF_000217795.1__O__Thermodesulfatatoraceae/Thermodesulfatator_indicus
RS_GCF_000482825.1__O__Thermodesulfovibrionaceae/Thermodesulfovibrio_islandicus
RS_GCF_000012385.1__O__Rickettsiaceae/Rickettsia_bellii
RS_GCF_000013365.1__O__Xanthobacteraceae/Rhodopseudomonas_palustris
RS_GCF_000005845.2__O__Enterobacteriaceae/Escherichia_coli
RS_GCF_001878485.1__O__Campylobacteraceae/Campylobacter_jejuni
RS_GCF_000177635.2__O__Chrysiogenaceae/Desulfurispirillum_indicum
0.79
12.04
18.01
447.94
227.7424
298.7979
390.7217
492.62
153.68
284.4917
380.8092
149.11
221.45
552.75
35.7
159.45
17.4
199.86
262.2122
405.2468
7.634
642.31
764.03
267.0344
255.359
519.1517
940.72
428.7649
528.3721
598.3367
697.8052
1277.07
277.3985
357.6641
815.6813
449.8901
69.1898
2776.18
863.2248
285.61
740.9235
1225.6914
2487.2978
3023.5126
2726.6028
2295.9887
2578.6256
2845.7557
2117.1607
2633.3296
2278.9005
3006.7867
1955.0581
3079.2922
2372.8279
2603.3833
2252.9052
2822.5512
1881.9485
2370.3412
2638.7411
3451.9832
2250.214
2532.7784
2856.2931
1942.1489
2728.9996
3111.509
2925.4338
3500.0
3000.0
2500.0
2000.0
1500.0
1000.0
500.0
0.0

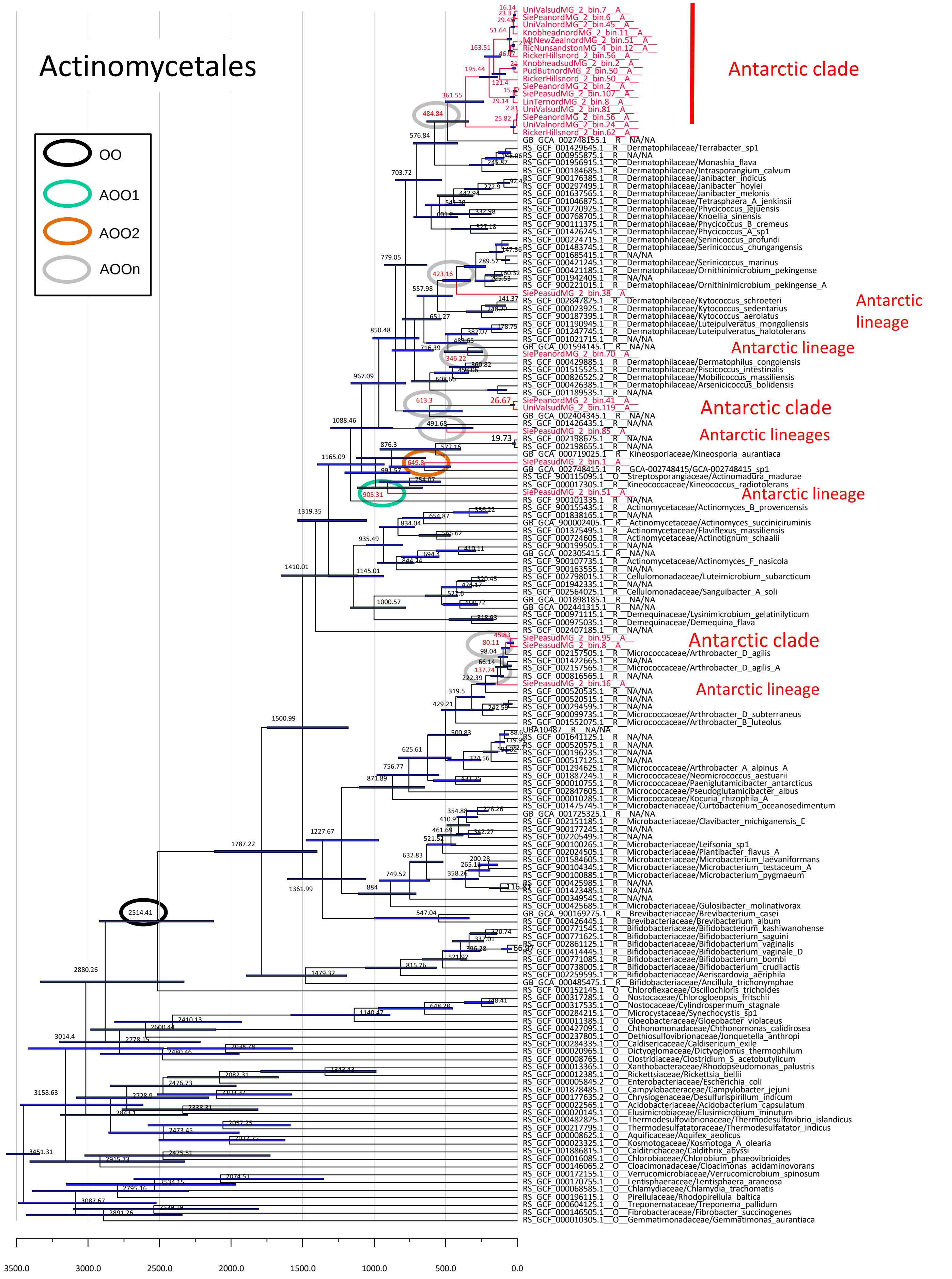

Actinomycetales
OO
AOO1
AOO2
AOOn
Antarctic clade
Antarctic lineage
Antarctic lineage
Antarctic clade
Antarctic lineages
Antarctic lineage
Antarctic clade
Antarctic lineage
UniValsudMG_2_bin.7__A__
SiePeanordMG_2_bin.6__A__
UniValnordMG_2_bin.45__A__
KnobheadnordMG_2_bin.11__A__
MtNewZealnordMG_2_bin.51__A__
RicNunsandstonMG_4_bin.12__A__
RickerHillsnord_2_bin.56__A__
KnobheadsudMG_2_bin.2__A__
PudButnordMG_2_bin.50__A__
RickerHillsnord_2_bin.50__A__
SiePeanordMG_2_bin.2__A__
SiePeasudMG_2_bin.107__A__
LinTernordMG_2_bin.8__A__
UniValsudMG_2_bin.81__A__
SiePeanordMG_2_bin.56__A__
UniValnordMG_2_bin.24__A__
RickerHillsnord_2_bin.62__A__
GB_GCA_002748155.1__R__NA/NA
RS_GCF_001429645.1__R__Dermatophilaceae/Terrabacter_sp1
RS_GCF_000955875.1__R__NA/NA
RS_GCF_001956915.1__R__Dermatophilaceae/Monashia_flava
RS_GCF_000184685.1__R__Dermatophilaceae/Intrasporangium_calvum
RS_GCF_900176385.1__R__Dermatophilaceae/Janibacter_indicus
RS_GCF_000297495.1__R__Dermatophilaceae/Janibacter_hoylei
RS_GCF_001637565.1__R__Dermatophilaceae/Janibacter_melonis
RS_GCF_001046875.1__R__Dermatophilaceae/Tetrasphaera_A_jenkinsii
RS_GCF_000720925.1__R__Dermatophilaceae/Phycicoccus_jejuensis
RS_GCF_000768705.1__R__Dermatophilaceae/Knoellia_sinensis
RS_GCF_900111375.1__R__Dermatophilaceae/Phycicoccus_B_cremeus
RS_GCF_001426245.1__R__Dermatophilaceae/Phycicoccus_A_sp1
RS_GCF_000224715.1__R__Dermatophilaceae/Serinicoccus_profundi
RS_GCF_001483745.1__R__Dermatophilaceae/Serinicoccus_chungangensis
RS_GCF_001685415.1__R__NA/NA
RS_GCF_000421245.1__R__Dermatophilaceae/Serinicoccus_marinus
RS_GCF_000421185.1__R__Dermatophilaceae/Ornithinimicrobium_pekingense
RS_GCF_001942405.1__R__NA/NA
RS_GCF_900221015.1__R__Dermatophilaceae/Ornithinimicrobium_pekingense_A
SiePeasudMG_2_bin.38__A__
RS_GCF_002847825.1__R__Dermatophilaceae/Kytococcus_schroeteri
RS_GCF_000023925.1__R__Dermatophilaceae/Kytococcus_sedentarius
RS_GCF_900187395.1__R__Dermatophilaceae/Kytococcus_aerolatus
RS_GCF_001190945.1__R__Dermatophilaceae/Luteipulveratus_mongoliensis
RS_GCF_001247745.1__R__Dermatophilaceae/Luteipulveratus_halotolerans
RS_GCF_001021715.1__R__NA/NA
GB_GCA_001594145.1__R__NA/NA
SiePeanordMG_2_bin.70__A__
RS_GCF_000429885.1__R__Dermatophilaceae/Dermatophilus_congolensis
RS_GCF_001515525.1__R__Dermatophilaceae/Piscicoccus_intestinalis
RS_GCF_000826525.2__R__Dermatophilaceae/Mobilicoccus_massiliensis
RS_GCF_000426385.1__R__Dermatophilaceae/Arsenicicoccus_bolidensis
RS_GCF_001189535.1__R__NA/NA
SiePeanordMG_2_bin.41__A__
UniValsudMG_2_bin.119__A__
GB_GCA_002404345.1__R__NA/NA
RS_GCF_001426435.1__R__NA/NA
SiePeasudMG_2_bin.85__A__
RS_GCF_002198675.1__R__NA/NA
RS_GCF_002198655.1__R__NA/NA
GB_GCA_000719025.1__R__Kineosporiaceae/Kineosporia_aurantiaca
SiePeasudMG_2_bin.1__A__
GB_GCA_002748415.1__R__GCA-002748415/GCA-002748415_sp1
RS_GCF_900115095.1__O__Streptosporangiaceae/Actinomadura_madurae
RS_GCF_000017305.1__R__Kineococcaceae/Kineococcus_radiotolerans
SiePeasudMG_2_bin.51__A__
RS_GCF_900101335.1__R__NA/NA
RS_GCF_900155435.1__R__Actinomycetaceae/Actinomyces_B_provencensis
RS_GCF_001838165.1__R__NA/NA
GB_GCA_900002405.1__R__Actinomycetaceae/Actinomyces_succiniciruminis
RS_GCF_001375495.1__R__Actinomycetaceae/Flaviflexus_massiliensis
RS_GCF_000724605.1__R__Actinomycetaceae/Actinotignum_schaalii
RS_GCF_900199505.1__R__NA/NA
GB_GCA_002305415.1__R__NA/NA
RS_GCF_900107735.1__R__Actinomycetaceae/Actinomyces_F_nasicola
RS_GCF_900163555.1__R__NA/NA
RS_GCF_002798015.1__R__Cellulomonadaceae/Luteimicrobium_subarcticum
RS_GCF_001942335.1__R__NA/NA
RS_GCF_002564025.1__R__Cellulomonadaceae/Sanguibacter_A_soli
GB_GCA_001898185.1__R__NA/NA
GB_GCA_002441315.1__R__NA/NA
RS_GCF_000971115.1__R__Demequinaceae/Lysinimicrobium_gelatinilyticum
RS_GCF_000975035.1__R__Demequinaceae/Demequina_flava
RS_GCF_002407185.1__R__NA/NA
SiePeasudMG_2_bin.95__A__
SiePeasudMG_2_bin.8__A__
RS_GCF_002157505.1__R__Micrococcaceae/Arthrobacter_D_agilis
RS_GCF_001422665.1__R__NA/NA
RS_GCF_002157565.1__R__Micrococcaceae/Arthrobacter_D_agilis_A
RS_GCF_000816565.1__R__NA/NA
SiePeasudMG_2_bin.16__A__
RS_GCF_000520535.1__R__NA/NA
RS_GCF_000520515.1__R__NA/NA
RS_GCF_000294595.1__R__NA/NA
RS_GCF_900099735.1__R__Micrococcaceae/Arthrobacter_D_subterraneus
RS_GCF_001552075.1__R__Micrococcaceae/Arthrobacter_B_luteolus
UBA10487__R__NA/NA
RS_GCF_001641125.1__R__NA/NA
RS_GCF_000520575.1__R__NA/NA
RS_GCF_000196235.1__R__NA/NA
RS_GCF_000517125.1__R__NA/NA
RS_GCF_001294625.1__R__Micrococcaceae/Arthrobacter_A_alpinus_A
RS_GCF_001887245.1__R__Micrococcaceae/Neomicrococcus_aestuarii
RS_GCF_900010755.1__R__Micrococcaceae/Paeniglutamicibacter_antarcticus
RS_GCF_002847605.1__R__Micrococcaceae/Pseudoglutamicibacter_albus
RS_GCF_000010285.1__R__Micrococcaceae/Kocuria_rhizophila_A
RS_GCF_001475745.1__R__Microbacteriaceae/Curtobacterium_oceanosedimentum
GB_GCA_001725325.1__R__NA/NA
RS_GCF_002151185.1__R__Microbacteriaceae/Clavibacter_michiganensis_E
RS_GCF_900177245.1__R__NA/NA
RS_GCF_002205495.1__R__NA/NA
RS_GCF_900100265.1__R__Microbacteriaceae/Leifsonia_sp1
RS_GCF_002024505.1__R__Microbacteriaceae/Plantibacter_flavus_A
RS_GCF_001584605.1__R__Microbacteriaceae/Microbacterium_laevaniformans
RS_GCF_900104345.1__R__Microbacteriaceae/Microbacterium_testaceum_A
RS_GCF_900100885.1__R__Microbacteriaceae/Microbacterium_pygmaeum
RS_GCF_000425985.1__R__NA/NA
RS_GCF_001423485.1__R__NA/NA
RS_GCF_000349545.1__R__NA/NA
RS_GCF_000425685.1__R__Microbacteriaceae/Gulosibacter_molinativorax
GB_GCA_900169275.1__R__Brevibacteriaceae/Brevibacterium_casei
RS_GCF_000426445.1__R__Brevibacteriaceae/Brevibacterium_album
RS_GCF_000771545.1__R__Bifidobacteriaceae/Bifidobacterium_kashiwanohense
RS_GCF_000771625.1__R__Bifidobacteriaceae/Bifidobacterium_saguini
RS_GCF_002861125.1__R__Bifidobacteriaceae/Bifidobacterium_vaginalis
RS_GCF_000414445.1__R__Bifidobacteriaceae/Bifidobacterium_vaginale_D
RS_GCF_000771085.1__R__Bifidobacteriaceae/Bifidobacterium_bombi
RS_GCF_000738005.1__R__Bifidobacteriaceae/Bifidobacterium_crudilactis
RS_GCF_002259595.1__R__Bifidobacteriaceae/Aeriscardovia_aeriphila
GB_GCA_000485475.1__R__Bifidobacteriaceae/Ancillula_trichonymphae
RS_GCF_000152145.1__O__Chloroflexaceae/Oscillochloris_trichoides
RS_GCF_000317285.1__O__Nostocaceae/Chlorogloeopsis_fritschii
RS_GCF_000317535.1__O__Nostocaceae/Cylindrospermum_stagnale
RS_GCF_000284215.1__O__Microcystaceae/Synechocystis_sp1
RS_GCF_000011385.1__O__Gloeobacteraceae/Gloeobacter_violaceus
RS_GCF_000427095.1__O__Chthonomonadaceae/Chthonomonas_calidirosea
RS_GCF_000237805.1__O__Dethiosulfovibrionaceae/Jonquetella_anthropi
RS_GCF_000284335.1__O__Caldisericaceae/Caldisericum_exile
RS_GCF_000020965.1__O__Dictyoglomaceae/Dictyoglomus_thermophilum
RS_GCF_000008765.1__O__Clostridiaceae/Clostridium_S_acetobutylicum
RS_GCF_000013365.1__O__Xanthobacteraceae/Rhodopseudomonas_palustris
RS_GCF_000012385.1__O__Rickettsiaceae/Rickettsia_bellii
RS_GCF_000005845.2__O__Enterobacteriaceae/Escherichia_coli
RS_GCF_001878485.1__O__Campylobacteraceae/Campylobacter_jejuni
RS_GCF_000177635.2__O__Chrysiogenaceae/Desulfurispirillum_indicum
RS_GCF_000022565.1__O__Acidobacteriaceae/Acidobacterium_capsulatum
RS_GCF_000020145.1__O__Elusimicrobiaceae/Elusimicrobium_minutum
RS_GCF_000482825.1__O__Thermodesulfovibrionaceae/Thermodesulfovibrio_islandicus
RS_GCF_000217795.1__O__Thermodesulfatatoraceae/Thermodesulfatator_indicus
RS_GCF_000008625.1__O__Aquificaceae/Aquifex_aeolicus
RS_GCF_000023325.1__O__Kosmotogaceae/Kosmotoga_A_olearia
RS_GCF_001886815.1__O__Caldithrichaceae/Caldithrix_abyssi
RS_GCF_000016085.1__O__Chlorobiaceae/Chlorobium_phaeovibrioides
RS_GCF_000146065.2__O__Cloacimonadaceae/Cloacimonas_acidaminovorans
RS_GCF_000172155.1__O__Verrucomicrobiaceae/Verrucomicrobium_spinosum
RS_GCF_000170755.1__O__Lentisphaeraceae/Lentisphaera_araneosa
RS_GCF_000068585.1__O__Chlamydiaceae/Chlamydia_trachomatis
RS_GCF_000196115.1__O__Pirellulaceae/Rhodopirellula_baltica
RS_GCF_000604125.1__O__Treponemataceae/Treponema_pallidum
RS_GCF_000146505.1__O__Fibrobacteraceae/Fibrobacter_succinogenes
RS_GCF_000010305.1__O__Gemmatimonadaceae/Gemmatimonas_aurantiaca
3500.0
3000.0
2500.0
2000.0
1500.0
1000.0
500.0
0.0

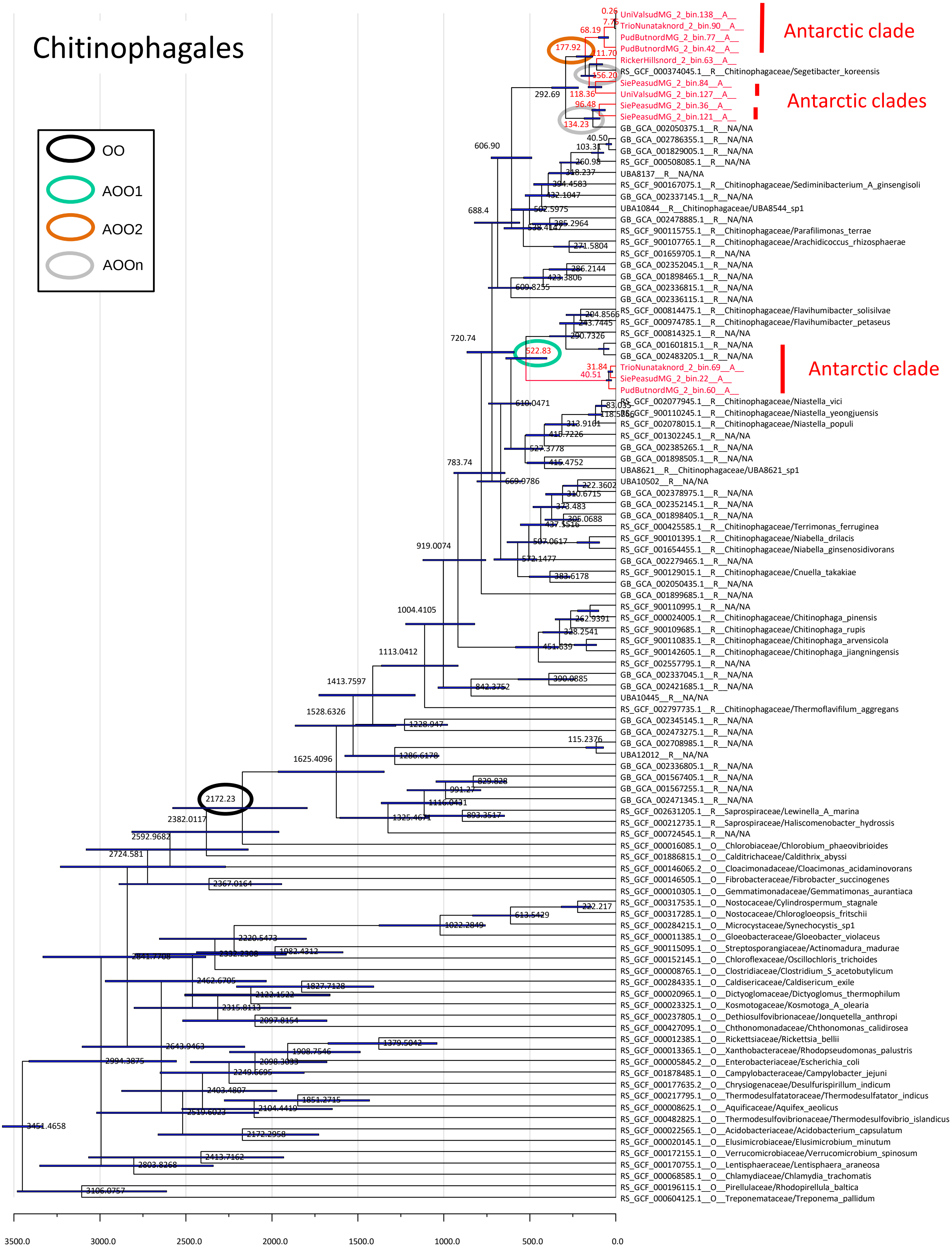

Chitinophagales
OO
AOO1
AOO2
AOOn
Antarctic clade
Antarctic clades
Antarctic clade
UniValsudMG_2_bin.138__A__
TrioNunataknord_2_bin.90__A__
PudButnordMG_2_bin.77__A__
PudButnordMG_2_bin.42__A__
RickerHillsnord_2_bin.63__A__
RS_GCF_000374045.1__R__Chitinophagaceae/Segetibacter_koreensis
SiePeasudMG_2_bin.84__A__
UniValsudMG_2_bin.127__A__
SiePeasudMG_2_bin.36__A__
SiePeasudMG_2_bin.121__A__
GB_GCA_002050375.1__R__NA/NA
GB_GCA_002786355.1__R__NA/NA
GB_GCA_001829005.1__R__NA/NA
RS_GCF_000508085.1__R__NA/NA
UBA8137__R__NA/NA
RS_GCF_900167075.1__R__Chitinophagaceae/Sediminibacterium_A_ginsengisoli
GB_GCA_002337145.1__R__NA/NA
UBA10844__R__Chitinophagaceae/UBA8544_sp1
GB_GCA_002478885.1__R__NA/NA
RS_GCF_900115755.1__R__Chitinophagaceae/Parafilimonas_terrae
RS_GCF_900107765.1__R__Chitinophagaceae/Arachidicoccus_rhizosphaerae
RS_GCF_001659705.1__R__NA/NA
GB_GCA_002352045.1__R__NA/NA
GB_GCA_001898465.1__R__NA/NA
GB_GCA_002336815.1__R__NA/NA
GB_GCA_002336115.1__R__NA/NA
RS_GCF_000814475.1__R__Chitinophagaceae/Flavihumibacter_solisilvae
RS_GCF_000974785.1__R__Chitinophagaceae/Flavihumibacter_petaseus
RS_GCF_000814325.1__R__NA/NA
GB_GCA_001601815.1__R__NA/NA
GB_GCA_002483205.1__R__NA/NA
TrioNunataknord_2_bin.69__A__
SiePeasudMG_2_bin.22__A__
PudButnordMG_2_bin.60__A__
RS_GCF_002077945.1__R__Chitinophagaceae/Niastella_vici
RS_GCF_900110245.1__R__Chitinophagaceae/Niastella_yeongjuensis
RS_GCF_002078015.1__R__Chitinophagaceae/Niastella_populi
RS_GCF_001302245.1__R__NA/NA
RS_GCF_002385265.1__R__NA/NA
GB_GCA_001898505.1__R__NA/NA
UBA8621__R__Chitinophagaceae/UBA8621_sp1
UBA10502__R__NA/NA
GB_GCA_002378975.1__R__NA/NA
GB_GCA_002352145.1__R__NA/NA
GB_GCA_001898405.1__R__NA/NA
RS_GCF_000425585.1__R__Chitinophagaceae/Terrimonas_ferruginea
RS_GCF_900101395.1__R__Chitinophagaceae/Niabella_drilacis
RS_GCF_001654455.1__R__Chitinophagaceae/Niabella_ginsenosidivorans
GB_GCA_002279465.1__R__NA/NA
RS_GCF_900129015.1__R__Chitinophagaceae/Cnuella_takakiae
GB_GCA_002050435.1__R__NA/NA
GB_GCA_001899685.1__R__NA/NA
RS_GCF_900110995.1__R__NA/NA
RS_GCF_000024005.1__R__Chitinophagaceae/Chitinophaga_pinensis
RS_GCF_900109685.1__R__Chitinophagaceae/Chitinophaga_rupis
RS_GCF_900110835.1__R__Chitinophagaceae/Chitinophaga_arvensicola
RS_GCF_900142605.1__R__Chitinophagaceae/Chitinophaga_jiangningensis
RS_GCF_002557795.1__R__NA/NA
GB_GCA_002337045.1__R__NA/NA
GB_GCA_002421685.1__R__NA/NA
UBA10445__R__NA/NA
RS_GCF_002797735.1__R__Chitinophagaceae/Thermoflavifilum_aggregans
GB_GCA_002345145.1__R__NA/NA
GB_GCA_002473275.1__R__NA/NA
GB_GCA_002708985.1__R__NA/NA
UBA12012__R__NA/NA
GB_GCA_002336805.1__R__NA/NA
GB_GCA_001567405.1__R__NA/NA
GB_GCA_001567255.1__R__NA/NA
GB_GCA_002471345.1__R__NA/NA
RS_GCF_002631205.1__R__Saprospiraceae/Lewinella_A_marina
RS_GCF_000212735.1__R__Saprospiraceae/Haliscomenobacter_hydrossis
RS_GCF_000724545.1__R__NA/NA
RS_GCF_000016085.1__O__Chlorobiaceae/Chlorobium_phaeovibrioides
RS_GCF_001886815.1__O__Calditrichaceae/Caldithrix_abyssi
RS_GCF_000146065.2__O__Cloacimonadaceae/Cloacimonas_acidaminovorans
RS_GCF_000146505.1__O__Fibrobacteraceae/Fibrobacter_succinogenes
RS_GCF_000010305.1__O__Gemmatimonadaceae/Gemmatimonas_aurantiaca
RS_GCF_000317535.1__O__Nostocaceae/Cylindrospermum_stagnale
RS_GCF_000317285.1__O__Nostocaceae/Chlorogloeopsis_fritschii
RS_GCF_000284215.1__O__Microcystaceae/Synechocystis_sp1
RS_GCF_000011385.1__O__Gloeobacteraceae/Gloeobacter_violaceus
RS_GCF_900115095.1__O__Streptosporangiaceae/Actinomadura_madurae
RS_GCF_000152145.1__O__Chloroflexaceae/Oscillochloris_trichoides
RS_GCF_000008765.1__O__Clostridiaceae/Clostridium_S_acetobutylicum
RS_GCF_000284335.1__O__Caldisericaceae/Caldisericum_exile
RS_GCF_000020965.1__O__Dictyoglomaceae/Dictyoglomus_thermophilum
RS_GCF_000023325.1__O__Kosmotogaceae/Kosmotoga_A_olearia
RS_GCF_000237805.1__O__Dethiosulfovibrionaceae/Jonquetella_anthropi
RS_GCF_000427095.1__O__Chthonomonadaceae/Chthonomonas_calidirosea
RS_GCF_000012385.1__O__Rickettsiaceae/Rickettsia_bellii
RS_GCF_000013365.1__O__Xanthobacteraceae/Rhodopseudomonas_palustris
RS_GCF_000005845.2__O__Enterobacteriaceae/Escherichia_coli
RS_GCF_001878485.1__O__Campylobacteraceae/Campylobacter_jejuni
RS_GCF_000177635.2__O__Chrysiogenaceae/Desulfurispirillum_indicum
RS_GCF_000217795.1__O__Thermodesulfatatoraceae/Thermodesulfatator_indicus
RS_GCF_000008625.1__O__Aquificaceae/Aquifex_aeolicus
RS_GCF_000482825.1__O__Thermodesulfovibrionaceae/Thermodesulfovibrio_islandicus
RS_GCF_000022565.1__O__Acidobacteriaceae/Acidobacterium_capsulatum
RS_GCF_000020145.1__O__Elusimicrobiaceae/Elusimicrobium_minutum
RS_GCF_000172155.1__O__Verrucomicrobiaceae/Verrucomicrobium_spinosum
RS_GCF_000170755.1__O__Lentisphaeraceae/Lentisphaera_araneosa
RS_GCF_000068585.1__O__Chlamydiaceae/Chlamydia_trachomatis
RS_GCF_000196115.1__O__Pirellulaceae/Rhodopirellula_baltica
RS_GCF_000604125.1__O__Treponemataceae/Treponema_pallidum
177.92
522.83
2172.23
292.69
606.90
688.4
720.74
783.74
919.0074
1004.4105
1113.0412
1413.7597
1528.6326
1625.4096
2382.0117
2592.9682
2724.581
2994.3875
3451.4658
3500.0
3000.0
2500.0
2000.0
1500.0
1000.0
500.0
0.0

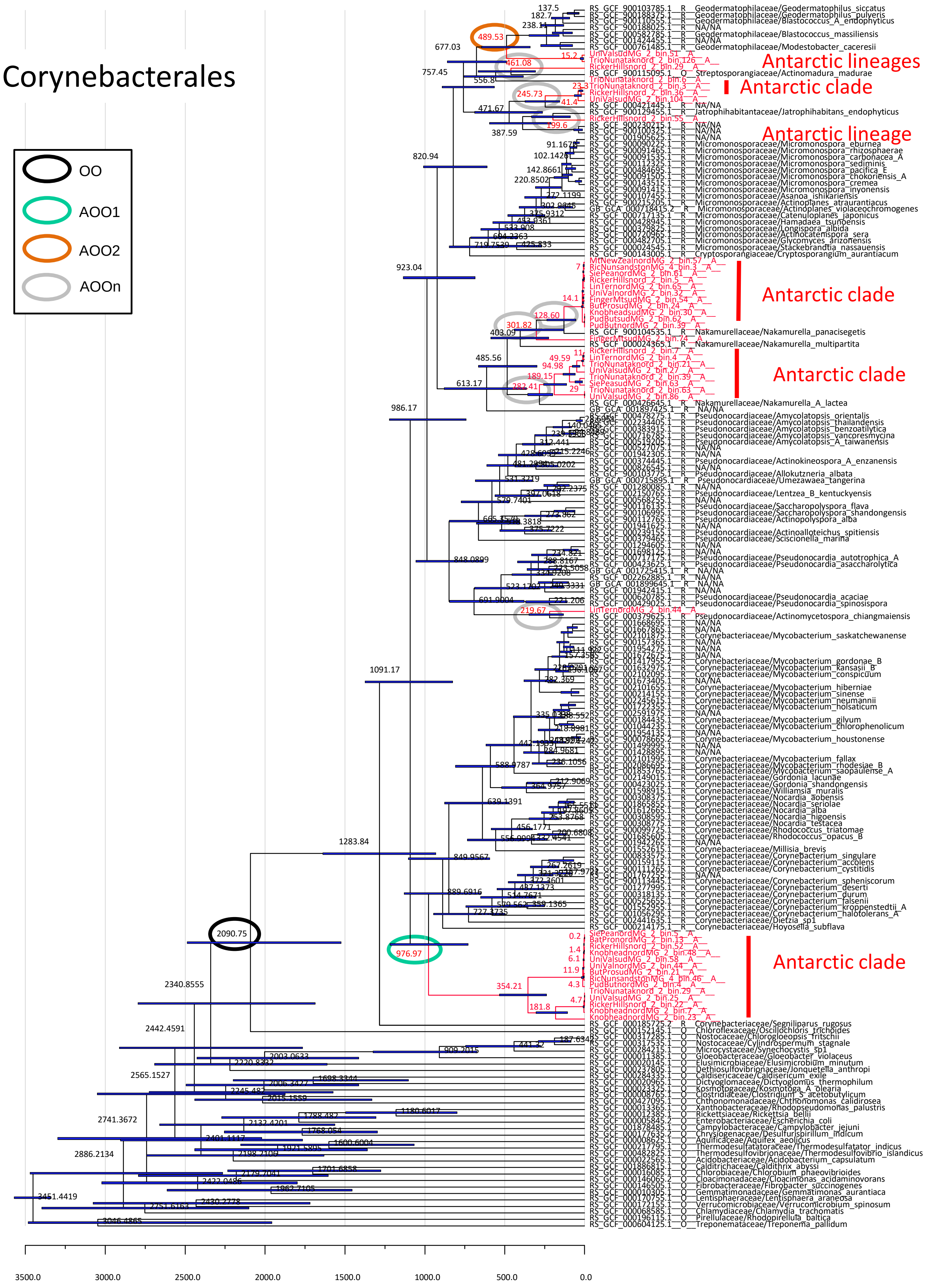
Corynebacterales
OO
AOO1
AOO2
AOOn
Antarctic lineages
Antarctic clade
Antarctic lineage
Antarctic clade
Antarctic clade
Antarctic clade
3500.0
3000.0
2500.0
2000.0
1500.0
1000.0
500.0
0.0

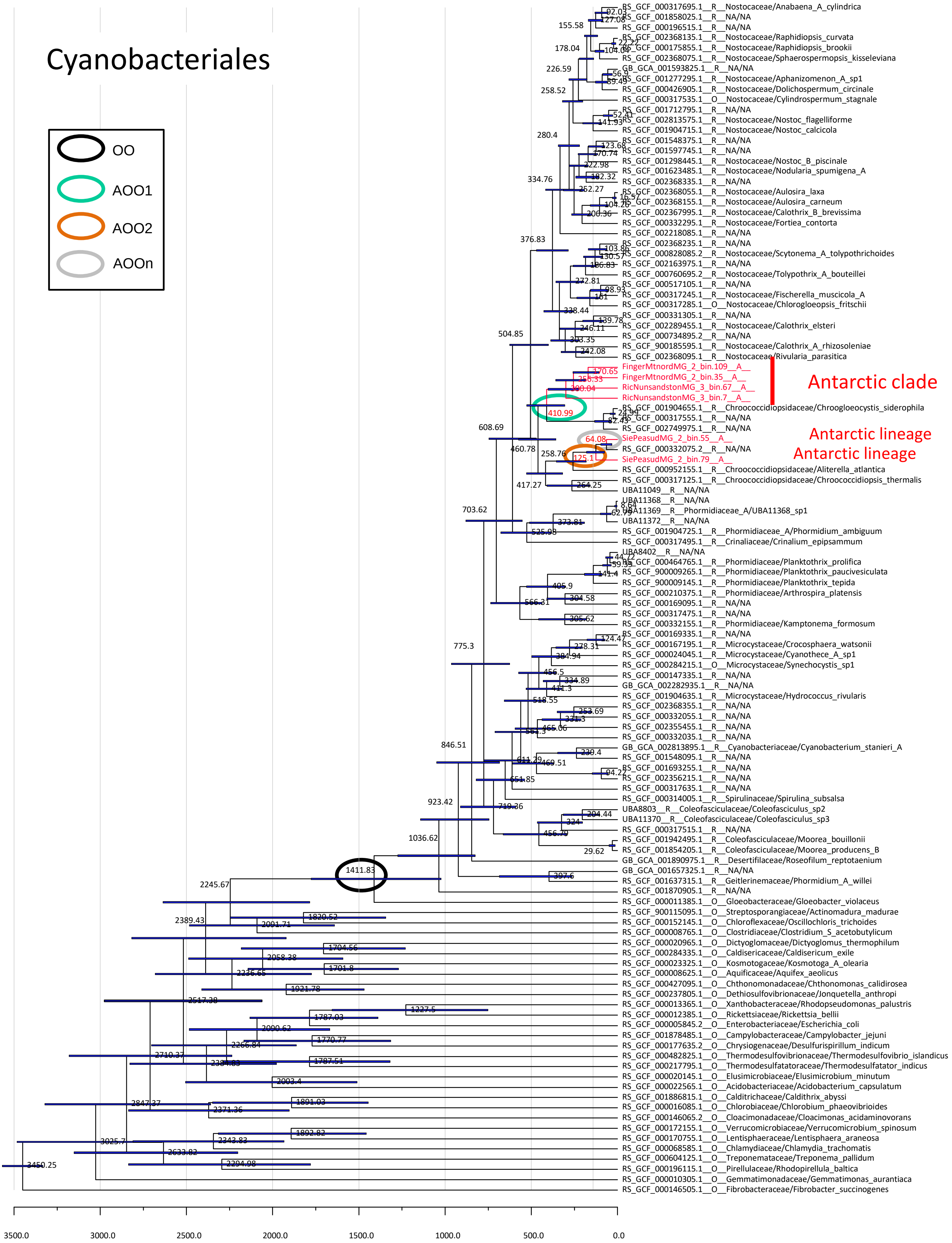

Cyanobacteriales
OO
AOO1
AOO2
AOOn
Antarctic clade
Antarctic lineage
Antarctic lineage
RS_GCF_000317695.1__R__Nostocaceae/Anabaena_A_cylindrica
RS_GCF_001858025.1__R__NA/NA
RS_GCF_000196515.1__R__NA/NA
RS_GCF_002368135.1__R__Nostocaceae/Raphidiopsis_curvata
RS_GCF_000175855.1__R__Nostocaceae/Raphidiopsis_brookii
RS_GCF_002368075.1__R__Nostocaceae/Sphaerospermopsis_kisseleviana
GB_GCA_001593825.1__R__NA/NA
RS_GCF_001277295.1__R__Nostocaceae/Aphanizomenon_A_sp1
RS_GCF_000426905.1__R__Nostocaceae/Dolichospermum_circinale
RS_GCF_000317535.1__O__Nostocaceae/Cylindrospermum_stagnale
RS_GCF_001712795.1__R__NA/NA
RS_GCF_002813575.1__R__Nostocaceae/Nostoc_flagelliforme
RS_GCF_001904715.1__R__Nostocaceae/Nostoc_calcicola
RS_GCF_001548375.1__R__NA/NA
RS_GCF_001597745.1__R__NA/NA
RS_GCF_001298445.1__R__Nostocaceae/Nostoc_B_piscinale
RS_GCF_001623485.1__R__Nostocaceae/Nodularia_spumigena_A
RS_GCF_002368335.1__R__NA/NA
RS_GCF_002368055.1__R__Nostocaceae/Aulosira_laxa
RS_GCF_002368155.1__R__Nostocaceae/Aulosira_carneum
RS_GCF_002367995.1__R__Nostocaceae/Calothrix_B_brevissima
RS_GCF_000332295.1__R__Nostocaceae/Fortiea_contorta
RS_GCF_002218085.1__R__NA/NA
RS_GCF_002368235.1__R__NA/NA
RS_GCF_000828085.2__R__Nostocaceae/Scytonema_A_tolypothrichoides
RS_GCF_002163975.1__R__NA/NA
RS_GCF_000760695.2__R__Nostocaceae/Tolypothrix_A_bouteillei
RS_GCF_000517105.1__R__NA/NA
RS_GCF_000317245.1__R__Nostocaceae/Fischerella_muscicola_A
RS_GCF_000317285.1__O__Nostocaceae/Chlorogloeopsis_fritschii
RS_GCF_000331305.1__R__NA/NA
RS_GCF_002289455.1__R__Nostocaceae/Calothrix_elsteri
RS_GCF_000734895.2__R__NA/NA
RS_GCF_900185595.1__R__Nostocaceae/Calothrix_A_rhizosoleniae
RS_GCF_002368095.1__R__Nostocaceae/Rivularia_parasitica
FingerMtnordMG_2_bin.109__A__
FingerMtnordMG_2_bin.35__A__
RicNunsandstonMG_3_bin.67__A__
RicNunsandstonMG_3_bin.7__A__
RS_GCF_001904655.1__R__Chroococcidiopsidaceae/Chroogloeocystis_siderophila
RS_GCF_000317555.1__R__NA/NA
RS_GCF_002749975.1__R__NA/NA
SiePeasudMG_2_bin.55__A__
RS_GCF_000332075.2__R__NA/NA
SiePeasudMG_2_bin.79__A__
RS_GCF_000952155.1__R__Chroococcidiopsidaceae/Aliterella_atlantica
RS_GCF_000317125.1__R__Chroococcidiopsidaceae/Chroococcidiopsis_thermalis
UBA11049__R__NA/NA
UBA11368__R__NA/NA
UBA11369__R__Phormidiaceae_A/UBA11368_sp1
UBA11372__R__NA/NA
RS_GCF_001904725.1__R__Phormidiaceae_A/Phormidium_ambiguum
RS_GCF_000317495.1__R__Crinaliaceae/Crinalium_epipsammum
UBA8402__R__NA/NA
RS_GCF_000464765.1__R__Phormidiaceae/Planktothrix_prolifica
RS_GCF_900009265.1__R__Phormidiaceae/Planktothrix_paucivesiculata
RS_GCF_900009145.1__R__Phormidiaceae/Planktothrix_tepida
RS_GCF_000210375.1__R__Phormidiaceae/Arthrospira_platensis
RS_GCF_000169095.1__R__NA/NA
RS_GCF_000317475.1__R__NA/NA
RS_GCF_000332155.1__R__Phormidiaceae/Kamptonema_formosum
RS_GCF_000169335.1__R__NA/NA
RS_GCF_000167195.1__R__Microcystaceae/Crocosphaera_watsonii
RS_GCF_000024045.1__R__Microcystaceae/Cyanothece_A_sp1
RS_GCF_000284215.1__O__Microcystaceae/Synechocystis_sp1
RS_GCF_000147335.1__R__NA/NA
GB_GCA_002282935.1__R__NA/NA
RS_GCF_001904635.1__R__Microcystaceae/Hydrococcus_rivularis
RS_GCF_002368355.1__R__NA/NA
RS_GCF_000332055.1__R__NA/NA
RS_GCF_002354455.1__R__NA/NA
RS_GCF_000332035.1__R__NA/NA
GB_GCA_002813895.1__R__Cyanobacteriaceae/Cyanobacterium_stanieri_A
RS_GCF_001548095.1__R__NA/NA
RS_GCF_001693255.1__R__NA/NA
RS_GCF_002356215.1__R__NA/NA
RS_GCF_000317635.1__R__NA/NA
RS_GCF_000314005.1__R__Spirulinaceae/Spirulina_subsalsa
UBA8803__R__Coleofasciculaceae/Coleofasciculus_sp2
UBA11370__R__Coleofasciculaceae/Coleofasciculus_sp3
RS_GCF_000317515.1__R__NA/NA
RS_GCF_001942495.1__R__Coleofasciculaceae/Moorea_bouillonii
RS_GCF_001854205.1__R__Coleofasciculaceae/Moorea_producens_B
GB_GCA_001890975.1__R__Desertifilaceae/Roseofilum_reptotaenium
GB_GCA_001657325.1__R__NA/NA
RS_GCF_001637315.1__R__Geitlerinemaceae/Phormidium_A_willei
RS_GCF_001870905.1__R__NA/NA
RS_GCF_000011385.1__O__Gloeobacteraceae/Gloeobacter_violaceus
RS_GCF_900115095.1__O__Streptosporangiaceae/Actinomadura_madurae
RS_GCF_000152145.1__O__Chloroflexaceae/Oscillochloris_trichoides
RS_GCF_000008765.1__O__Clostridiaceae/Clostridium_S_acetobutylicum
RS_GCF_000020965.1__O__Dictyoglomaceae/Dictyoglomus_thermophilum
RS_GCF_000284335.1__O__Caldisericaceae/Caldisericum_exile
RS_GCF_000023325.1__O__Kosmotogaceae/Kosmotoga_A_olearia
RS_GCF_000008625.1__O__Aquificaceae/Aquifex_aeolicus
RS_GCF_000427095.1__O__Chthonomonadaceae/Chthonomonas_calidirosea
RS_GCF_000237805.1__O__Dethiosulfovibrionaceae/Jonquetella_anthropi
RS_GCF_000013365.1__O__Xanthobacteraceae/Rhodopseudomonas_palustris
RS_GCF_000012385.1__O__Rickettsiaceae/Rickettsia_bellii
RS_GCF_000005845.2__O__Enterobacteriaceae/Escherichia_coli
RS_GCF_001878485.1__O__Campylobacteraceae/Campylobacter_jejuni
RS_GCF_000177635.2__O__Chrysiogenaceae/Desulfurispirillum_indicum
RS_GCF_000482825.1__O__Thermodesulfovibrionaceae/Thermodesulfovibrio_islandicus
RS_GCF_000217795.1__O__Thermodesulfatatoraceae/Thermodesulfatator_indicus
RS_GCF_000020145.1__O__Elusimicrobiaceae/Elusimicrobium_minutum
RS_GCF_000022565.1__O__Acidobacteriaceae/Acidobacterium_capsulatum
RS_GCF_001886815.1__O__Calditrichaceae/Caldithrix_abyssi
RS_GCF_000016085.1__O__Chlorobiaceae/Chlorobium_phaeovibrioides
RS_GCF_000146065.2__O__Cloacimonadaceae/Cloacimonas_acidaminovorans
RS_GCF_000172155.1__O__Verrucomicrobiaceae/Verrucomicrobium_spinosum
RS_GCF_000170755.1__O__Lentisphaeraceae/Lentisphaera_araneosa
RS_GCF_000068585.1__O__Chlamydiaceae/Chlamydia_trachomatis
RS_GCF_000604125.1__O__Treponemataceae/Treponema_pallidum
RS_GCF_000196115.1__O__Pirellulaceae/Rhodopirellula_baltica
RS_GCF_000010305.1__O__Gemmatimonadaceae/Gemmatimonas_aurantiaca
RS_GCF_000146505.1__O__Fibrobacteraceae/Fibrobacter_succinogenes
3500.0
3000.0
2500.0
2000.0
1500.0
1000.0
500.0
0.0

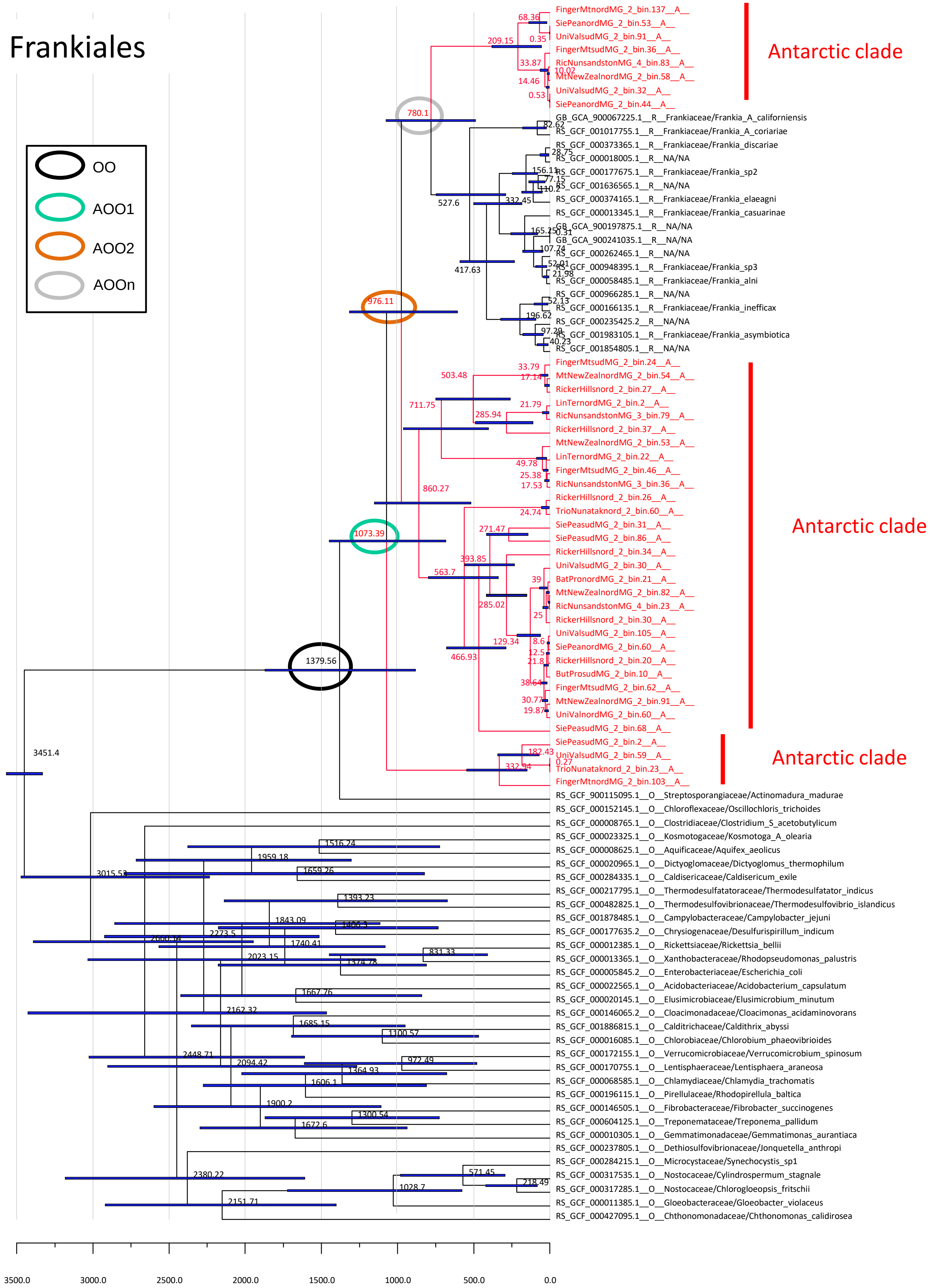

Frankiales
OO
AOO1
AOO2
AOOn
Antarctic clade
Antarctic clade
Antarctic clade
FingerMtnordMG_2_bin.137__A__
SiePeanordMG_2_bin.53__A__
UniValsudMG_2_bin.91__A__
FingerMtsudMG_2_bin.36__A__
RicNunsandstonMG_4_bin.83__A__
MtNewZealnordMG_2_bin.58__A__
UniValsudMG_2_bin.32__A__
SiePeanordMG_2_bin.44__A__
GB_GCA_900067225.1__R__Frankiaceae/Frankia_A_californiensis
RS_GCF_001017755.1__R__Frankiaceae/Frankia_A_coriariae
RS_GCF_000373365.1__R__Frankiaceae/Frankia_discariae
RS_GCF_000018005.1__R__NA/NA
RS_GCF_000177675.1__R__Frankiaceae/Frankia_sp2
RS_GCF_001636565.1__R__NA/NA
RS_GCF_000374165.1__R__Frankiaceae/Frankia_elaeagni
RS_GCF_000013345.1__R__Frankiaceae/Frankia_casuarinae
GB_GCA_900197875.1__R__NA/NA
GB_GCA_900241035.1__R__NA/NA
RS_GCF_000262465.1__R__NA/NA
RS_GCF_000948395.1__R__Frankiaceae/Frankia_sp3
RS_GCF_000058485.1__R__Frankiaceae/Frankia_alni
RS_GCF_000966285.1__R__NA/NA
RS_GCF_000166135.1__R__Frankiaceae/Frankia_inefficax
RS_GCF_000235425.2__R__NA/NA
RS_GCF_001983105.1__R__Frankiaceae/Frankia_asymbiotica
RS_GCF_001854805.1__R__NA/NA
FingerMtsudMG_2_bin.24__A__
MtNewZealnordMG_2_bin.54__A__
RickerHillsnord_2_bin.27__A__
LinTernordMG_2_bin.2__A__
RicNunsandstonMG_3_bin.79__A__
RickerHillsnord_2_bin.37__A__
MtNewZealnordMG_2_bin.53__A__
LinTernordMG_2_bin.22__A__
FingerMtsudMG_2_bin.46__A__
RicNunsandstonMG_3_bin.36__A__
RickerHillsnord_2_bin.26__A__
TrioNunataknord_2_bin.60__A__
SiePeasudMG_2_bin.31__A__
SiePeasudMG_2_bin.86__A__
RickerHillsnord_2_bin.34__A__
UniValsudMG_2_bin.30__A__
BatPronordMG_2_bin.21__A__
MtNewZealnordMG_2_bin.82__A__
RicNunsandstonMG_4_bin.23__A__
RickerHillsnord_2_bin.30__A__
UniValsudMG_2_bin.105__A__
SiePeanordMG_2_bin.60__A__
RickerHillsnord_2_bin.20__A__
ButProsudMG_2_bin.10__A__
FingerMtsudMG_2_bin.62__A__
MtNewZealnordMG_2_bin.91__A__
UniValnordMG_2_bin.60__A__
SiePeasudMG_2_bin.68__A__
SiePeasudMG_2_bin.2__A__
UniValsudMG_2_bin.59__A__
TrioNunataknord_2_bin.23__A__
FingerMtnordMG_2_bin.103__A__
RS_GCF_900115095.1__O__Streptosporangiaceae/Actinomadura_madurae
RS_GCF_000152145.1__O__Chloroflexaceae/Oscillochloris_trichoides
RS_GCF_000008765.1__O__Clostridiaceae/Clostridium_S_acetobutylicum
RS_GCF_000023325.1__O__Kosmotogaceae/Kosmotoga_A_olearia
RS_GCF_000008625.1__O__Aquificaceae/Aquifex_aeolicus
RS_GCF_000020965.1__O__Dictyoglomaceae/Dictyoglomus_thermophilum
RS_GCF_000284335.1__O__Caldisericaceae/Caldisericum_exile
RS_GCF_000217795.1__O__Thermodesulfatatoraceae/Thermodesulfatator_indicus
RS_GCF_000482825.1__O__Thermodesulfovibrionaceae/Thermodesulfovibrio_islandicus
RS_GCF_001878485.1__O__Campylobacteraceae/Campylobacter_jejuni
RS_GCF_000177635.2__O__Chrysiogenaceae/Desulfurispirillum_indicum
RS_GCF_000012385.1__O__Rickettsiaceae/Rickettsia_bellii
RS_GCF_000013365.1__O__Xanthobacteraceae/Rhodopseudomonas_palustris
RS_GCF_000005845.2__O__Enterobacteriaceae/Escherichia_coli
RS_GCF_000022565.1__O__Acidobacteriaceae/Acidobacterium_capsulatum
RS_GCF_000020145.1__O__Elusimicrobiaceae/Elusimicrobium_minutum
RS_GCF_000146065.2__O__Cloacimonadaceae/Cloacimonas_acidaminovorans
RS_GCF_001886815.1__O__Calditrichaceae/Caldithrix_abyssi
RS_GCF_000016085.1__O__Chlorobiaceae/Chlorobium_phaeovibrioides
RS_GCF_000172155.1__O__Verrucomicrobiaceae/Verrucomicrobium_spinosum
RS_GCF_000170755.1__O__Lentisphaeraceae/Lentisphaera_araneosa
RS_GCF_000068585.1__O__Chlamydiaceae/Chlamydia_trachomatis
RS_GCF_000196115.1__O__Pirellulaceae/Rhodopirellula_baltica
RS_GCF_000146505.1__O__Fibrobacteraceae/Fibrobacter_succinogenes
RS_GCF_000604125.1__O__Treponemataceae/Treponema_pallidum
RS_GCF_000010305.1__O__Gemmatimonadaceae/Gemmatimonas_aurantiaca
RS_GCF_000237805.1__O__Dethiosulfovibrionaceae/Jonquetella_anthropi
RS_GCF_000284215.1__O__Microcystaceae/Synechocystis_sp1
RS_GCF_000317535.1__O__Nostocaceae/Cylindrospermum_stagnale
RS_GCF_000317285.1__O__Nostocaceae/Chlorogloeopsis_fritschii
RS_GCF_000011385.1__O__Gloeobacteraceae/Gloeobacter_violaceus
RS_GCF_000427095.1__O__Chthonomonadaceae/Chthonomonas_calidirosea
3451.4
1379.56
1073.39
976.11
780.1
860.27
3500.0
3000.0
2500.0
2000.0
1500.0
1000.0
500.0
0.0

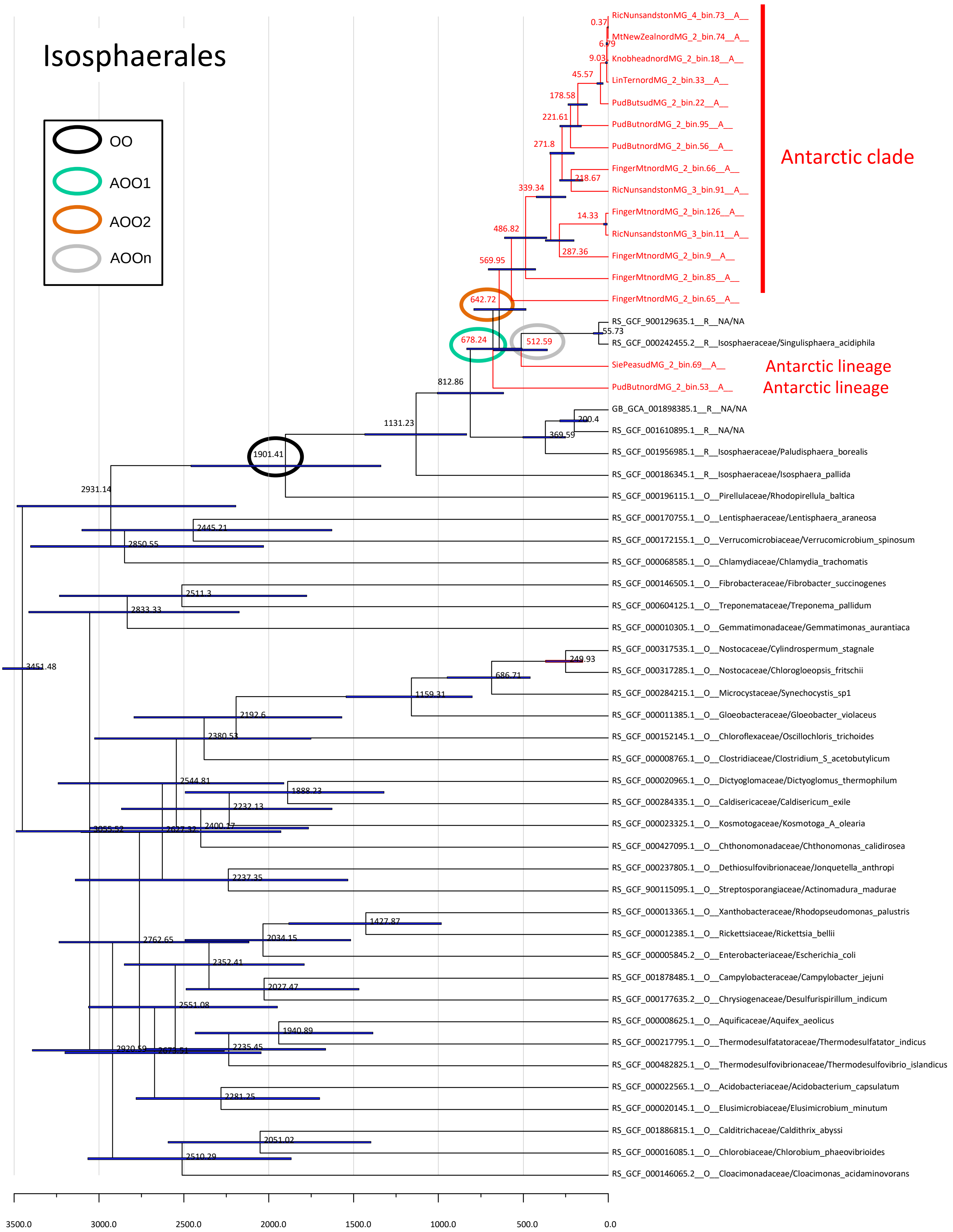

Isosphaerales
OO
AOO1
AOO2
AOOn
Antarctic clade
Antarctic lineage
Antarctic lineage
RicNunsandstonMG_4_bin.73__A__
MtNewZealnordMG_2_bin.74__A__
KnobheadnordMG_2_bin.18__A__
LinTernordMG_2_bin.33__A__
PudButsudMG_2_bin.22__A__
PudButnordMG_2_bin.95__A__
PudButnordMG_2_bin.56__A__
FingerMtnordMG_2_bin.66__A__
RicNunsandstonMG_3_bin.91__A__
FingerMtnordMG_2_bin.126__A__
RicNunsandstonMG_3_bin.11__A__
FingerMtnordMG_2_bin.9__A__
FingerMtnordMG_2_bin.85__A__
FingerMtnordMG_2_bin.65__A__
RS_GCF_900129635.1__R__NA/NA
RS_GCF_000242455.2__R__Isosphaeraceae/Singulisphaera_acidiphila
SiePeasudMG_2_bin.69__A__
PudButnordMG_2_bin.53__A__
GB_GCA_001898385.1__R__NA/NA
RS_GCF_001610895.1__R__NA/NA
RS_GCF_001956985.1__R__Isosphaeraceae/Paludisphaera_borealis
RS_GCF_000186345.1__R__Isosphaeraceae/Isosphaera_pallida
RS_GCF_000196115.1__O__Pirellulaceae/Rhodopirellula_baltica
RS_GCF_000170755.1__O__Lentisphaeraceae/Lentisphaera_araneosa
RS_GCF_000172155.1__O__Verrucomicrobiaceae/Verrucomicrobium_spinosum
RS_GCF_000068585.1__O__Chlamydiaceae/Chlamydia_trachomatis
RS_GCF_000146505.1__O__Fibrobacteraceae/Fibrobacter_succinogenes
RS_GCF_000604125.1__O__Treponemataceae/Treponema_pallidum
RS_GCF_000010305.1__O__Gemmatimonadaceae/Gemmatimonas_aurantiaca
RS_GCF_000317535.1__O__Nostocaceae/Cylindrospermum_stagnale
RS_GCF_000317285.1__O__Nostocaceae/Chlorogloeopsis_fritschii
RS_GCF_000284215.1__O__Microcystaceae/Synechocystis_sp1
RS_GCF_000011385.1__O__Gloeobacteraceae/Gloeobacter_violaceus
RS_GCF_000152145.1__O__Chloroflexaceae/Oscillochloris_trichoides
RS_GCF_000008765.1__O__Clostridiaceae/Clostridium_S_acetobutylicum
RS_GCF_000020965.1__O__Dictyoglomaceae/Dictyoglomus_thermophilum
RS_GCF_000284335.1__O__Caldisericaceae/Caldisericum_exile
RS_GCF_000023325.1__O__Kosmotogaceae/Kosmotoga_A_olearia
RS_GCF_000427095.1__O__Chthonomonadaceae/Chthonomonas_calidirosea
RS_GCF_000237805.1__O__Dethiosulfovibrionaceae/Jonquetella_anthropi
RS_GCF_900115095.1__O__Streptosporangiaceae/Actinomadura_madurae
RS_GCF_000013365.1__O__Xanthobacteraceae/Rhodopseudomonas_palustris
RS_GCF_000012385.1__O__Rickettsiaceae/Rickettsia_bellii
RS_GCF_000005845.2__O__Enterobacteriaceae/Escherichia_coli
RS_GCF_001878485.1__O__Campylobacteraceae/Campylobacter_jejuni
RS_GCF_000177635.2__O__Chrysiogenaceae/Desulfurispirillum_indicum
RS_GCF_000008625.1__O__Aquificaceae/Aquifex_aeolicus
RS_GCF_000217795.1__O__Thermodesulfatatoraceae/Thermodesulfatator_indicus
RS_GCF_000482825.1__O__Thermodesulfovibrionaceae/Thermodesulfovibrio_islandicus
RS_GCF_000022565.1__O__Acidobacteriaceae/Acidobacterium_capsulatum
RS_GCF_000020145.1__O__Elusimicrobiaceae/Elusimicrobium_minutum
RS_GCF_001886815.1__O__Calditrichaceae/Caldithrix_abyssi
RS_GCF_000016085.1__O__Chlorobiaceae/Chlorobium_phaeovibrioides
RS_GCF_000146065.2__O__Cloacimonadaceae/Cloacimonas_acidaminovorans
0.37
6.79
9.03
45.57
178.58
221.61
271.8
218.67
339.34
14.33
486.82
287.36
569.95
642.72
55.73
678.24
512.59
812.86
200.4
1131.23
369.59
1901.41
2931.14
2445.21
2850.55
2511.3
2833.33
249.93
3451.48
686.71
1159.31
2192.6
2380.53
2544.81
1888.23
2232.13
3055.52
2627.32
2400.17
2237.35
1427.87
2762.65
2034.15
2352.41
2027.47
2551.08
1940.89
2920.59
2673.51
2235.45
2281.25
2051.02
2510.29
3500.0
3000.0
2500.0
2000.0
1500.0
1000.0
500.0
0.0

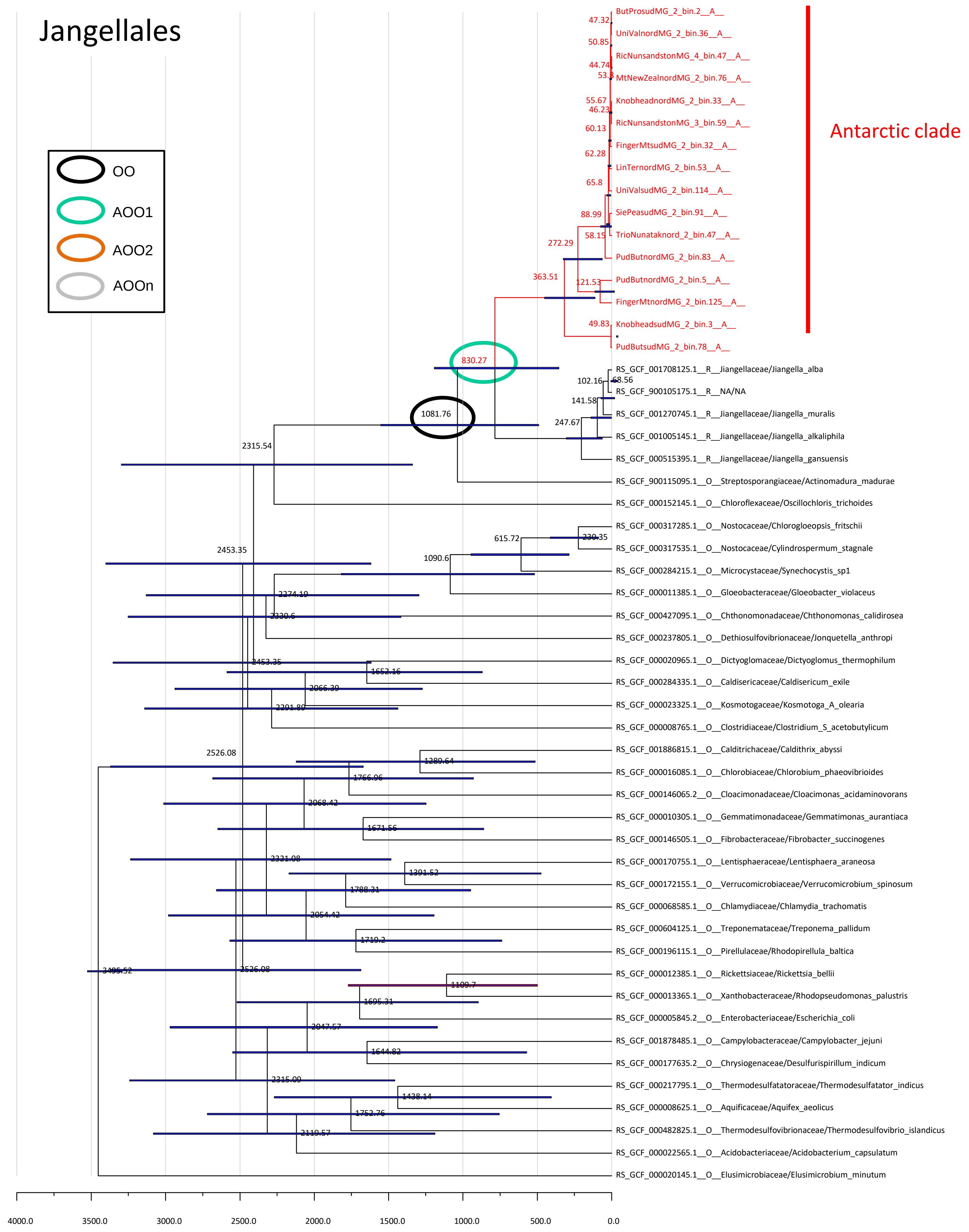
Jangellales
OO
AOO1
AOO2
AOOn
Antarctic clade
ButProsudMG_2_bin.2__A__
UniValnordMG_2_bin.36__A__
RicNunsandstonMG_4_bin.47__A__
MtNewZealnordMG_2_bin.76__A__
KnobheadnordMG_2_bin.33__A__
RicNunsandstonMG_3_bin.59__A__
FingerMtsudMG_2_bin.32__A__
LinTernordMG_2_bin.53__A__
UniValsudMG_2_bin.114__A__
SiePeasudMG_2_bin.91__A__
TrioNunataknord_2_bin.47__A__
PudButnordMG_2_bin.83__A__
PudButnordMG_2_bin.5__A__
FingerMtnordMG_2_bin.125__A__
KnobheadsudMG_2_bin.3__A__
PudButsudMG_2_bin.78__A__
RS_GCF_001708125.1__R__Jiangellaceae/Jiangella_alba
RS_GCF_900105175.1__R__NA/NA
RS_GCF_001270745.1__R__Jiangellaceae/Jiangella_muralis
RS_GCF_001005145.1__R__Jiangellaceae/Jiangella_alkaliphila
RS_GCF_000515395.1__R__Jiangellaceae/Jiangella_gansuensis
RS_GCF_900115095.1__O__Streptosporangiaceae/Actinomadura_madurae
RS_GCF_000152145.1__O__Chloroflexaceae/Oscillochloris_trichoides
RS_GCF_000317285.1__O__Nostocaceae/Chlorogloeopsis_fritschii
RS_GCF_000317535.1__O__Nostocaceae/Cylindrospermum_stagnale
RS_GCF_000284215.1__O__Microcystaceae/Synechocystis_sp1
RS_GCF_000011385.1__O__Gloeobacteraceae/Gloeobacter_violaceus
RS_GCF_000427095.1__O__Chthonomonadaceae/Chthonomonas_calidirosea
RS_GCF_000237805.1__O__Dethiosulfovibrionaceae/Jonquetella_anthropi
RS_GCF_000020965.1__O__Dictyoglomaceae/Dictyoglomus_thermophilum
RS_GCF_000284335.1__O__Caldisericaceae/Caldisericum_exile
RS_GCF_000023325.1__O__Kosmotogaceae/Kosmotoga_A_olearia
RS_GCF_000008765.1__O__Clostridiaceae/Clostridium_S_acetobutylicum
RS_GCF_001886815.1__O__Calditrichaceae/Caldithrix_abyssi
RS_GCF_000016085.1__O__Chlorobiaceae/Chlorobium_phaeovibrioides
RS_GCF_000146065.2__O__Cloacimonadaceae/Cloacimonas_acidaminovorans
RS_GCF_000010305.1__O__Gemmatimonadaceae/Gemmatimonas_aurantiaca
RS_GCF_000146505.1__O__Fibrobacteraceae/Fibrobacter_succinogenes
RS_GCF_000170755.1__O__Lentisphaeraceae/Lentisphaera_araneosa
RS_GCF_000172155.1__O__Verrucomicrobiaceae/Verrucomicrobium_spinosum
RS_GCF_000068585.1__O__Chlamydiaceae/Chlamydia_trachomatis
RS_GCF_000604125.1__O__Treponemataceae/Treponema_pallidum
RS_GCF_000196115.1__O__Pirellulaceae/Rhodopirellula_baltica
RS_GCF_000012385.1__O__Rickettsiaceae/Rickettsia_bellii
RS_GCF_000013365.1__O__Xanthobacteraceae/Rhodopseudomonas_palustris
RS_GCF_000005845.2__O__Enterobacteriaceae/Escherichia_coli
RS_GCF_001878485.1__O__Campylobacteraceae/Campylobacter_jejuni
RS_GCF_000177635.2__O__Chrysiogenaceae/Desulfurispirillum_indicum
RS_GCF_000217795.1__O__Thermodesulfatatoraceae/Thermodesulfatator_indicus
RS_GCF_000008625.1__O__Aquificaceae/Aquifex_aeolicus
RS_GCF_000482825.1__O__Thermodesulfovibrionaceae/Thermodesulfovibrio_islandicus
RS_GCF_000022565.1__O__Acidobacteriaceae/Acidobacterium_capsulatum
RS_GCF_000020145.1__O__Elusimicrobiaceae/Elusimicrobium_minutum
47.32
50.85
44.74
53.3
55.67
46.23
60.13
62.28
65.8
88.99
58.15
272.29
363.51
121.53
49.83
830.27
102.16
68.56
141.58
1081.76
247.67
2315.54
615.72
230.35
2453.35
1090.6
2274.19
2330.6
2453.35
1652.16
2066.39
2291.89
2526.08
1289.64
1766.96
2068.42
1671.56
2321.08
1391.52
1788.31
2054.42
1719.2
3495.52
2526.08
1109.7
1695.31
2047.57
1644.82
2315.09
1438.14
1752.76
2119.57
4000.0
3500.0
3000.0
2500.0
2000.0
1500.0
1000.0
500.0
0.0

### Ktedonobacterales

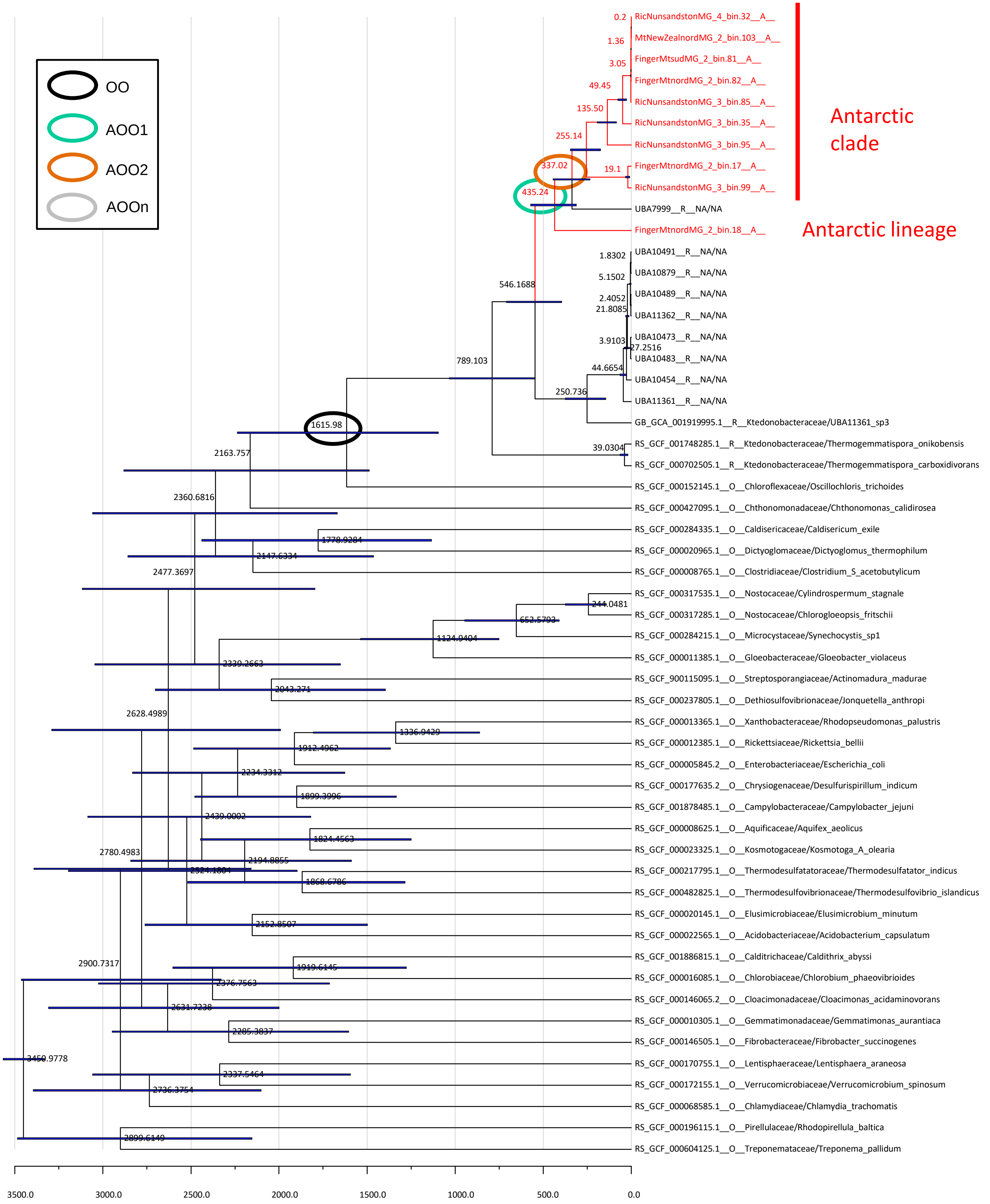

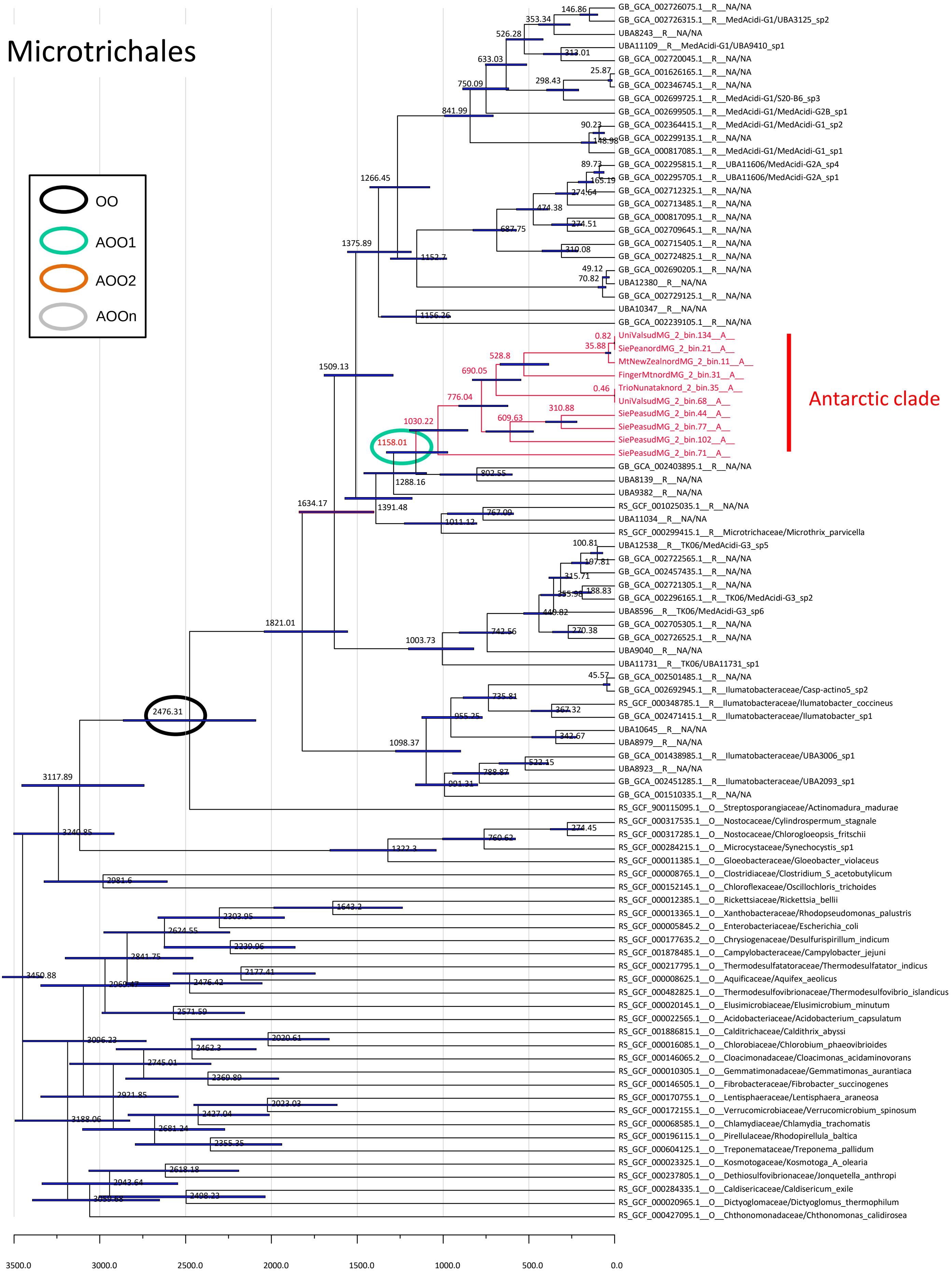
Microtrichales
OO
AOO1
AOO2
AOOn
Antarctic clade
GB_GCA_002726075.1__R__NA/NA
GB_GCA_002726315.1__R__MedAcidi-G1/UBA3125_sp2
UBA8243__R__NA/NA
UBA11109__R__MedAcidi-G1/UBA9410_sp1
GB_GCA_002720045.1__R__NA/NA
GB_GCA_001626165.1__R__NA/NA
GB_GCA_002346745.1__R__NA/NA
GB_GCA_002699725.1__R__MedAcidi-G1/S20-B6_sp3
GB_GCA_002699505.1__R__MedAcidi-G1/MedAcidi-G2B_sp1
GB_GCA_002364415.1__R__MedAcidi-G1/MedAcidi-G1_sp2
GB_GCA_002299135.1__R__NA/NA
GB_GCA_000817085.1__R__MedAcidi-G1/MedAcidi-G1_sp1
GB_GCA_002295815.1__R__UBA11606/MedAcidi-G2A_sp4
GB_GCA_002295705.1__R__UBA11606/MedAcidi-G2A_sp1
GB_GCA_002712325.1__R__NA/NA
GB_GCA_002713485.1__R__NA/NA
GB_GCA_000817095.1__R__NA/NA
GB_GCA_002709645.1__R__NA/NA
GB_GCA_002715405.1__R__NA/NA
GB_GCA_002724825.1__R__NA/NA
GB_GCA_002690205.1__R__NA/NA
UBA12380__R__NA/NA
GB_GCA_002729125.1__R__NA/NA
UBA10347__R__NA/NA
GB_GCA_002239105.1__R__NA/NA
UniValsudMG_2_bin.134__A__
SiePeanordMG_2_bin.21__A__
MtNewZealnordMG_2_bin.11__A__
FingerMtnordMG_2_bin.31__A__
TrioNunataknord_2_bin.35__A__
UniValsudMG_2_bin.68__A__
SiePeasudMG_2_bin.44__A__
SiePeasudMG_2_bin.77__A__
SiePeasudMG_2_bin.102__A__
SiePeasudMG_2_bin.71__A__
GB_GCA_002403895.1__R__NA/NA
UBA8139__R__NA/NA
UBA9382__R__NA/NA
RS_GCF_001025035.1__R__NA/NA
UBA11034__R__NA/NA
RS_GCF_000299415.1__R__Microtrichaceae/Microthrix_parvicella
UBA12538__R__TK06/MedAcidi-G3_sp5
GB_GCA_002722565.1__R__NA/NA
GB_GCA_002457435.1__R__NA/NA
GB_GCA_002721305.1__R__NA/NA
GB_GCA_002296165.1__R__TK06/MedAcidi-G3_sp2
UBA8596__R__TK06/MedAcidi-G3_sp6
GB_GCA_002705305.1__R__NA/NA
GB_GCA_002726525.1__R__NA/NA
UBA9040__R__NA/NA
UBA11731__R__TK06/UBA11731_sp1
GB_GCA_002501485.1__R__NA/NA
GB_GCA_002692945.1__R__Ilumatobacteraceae/Casp-actino5_sp2
RS_GCF_000348785.1__R__Ilumatobacteraceae/Ilumatobacter_coccineus
GB_GCA_002471415.1__R__Ilumatobacteraceae/Ilumatobacter_sp1
UBA10645__R__NA/NA
UBA8979__R__NA/NA
GB_GCA_001438985.1__R__Ilumatobacteraceae/UBA3006_sp1
UBA8923__R__NA/NA
GB_GCA_002451285.1__R__Ilumatobacteraceae/UBA2093_sp1
GB_GCA_001510335.1__R__NA/NA
RS_GCF_900115095.1__O__Streptosporangiaceae/Actinomadura_madurae
RS_GCF_000317535.1__O__Nostocaceae/Cylindrospermum_stagnale
RS_GCF_000317285.1__O__Nostocaceae/Chlorogloeopsis_fritschii
RS_GCF_000284215.1__O__Microcystaceae/Synechocystis_sp1
RS_GCF_000011385.1__O__Gloeobacteraceae/Gloeobacter_violaceus
RS_GCF_000008765.1__O__Clostridiaceae/Clostridium_S_acetobutylicum
RS_GCF_000152145.1__O__Chloroflexaceae/Oscillochloris_trichoides
RS_GCF_000012385.1__O__Rickettsiaceae/Rickettsia_bellii
RS_GCF_000013365.1__O__Xanthobacteraceae/Rhodopseudomonas_palustris
RS_GCF_000005845.2__O__Enterobacteriaceae/Escherichia_coli
RS_GCF_000177635.2__O__Chrysiogenaceae/Desulfurispirillum_indicum
RS_GCF_001878485.1__O__Campylobacteraceae/Campylobacter_jejuni
RS_GCF_000217795.1__O__Thermodesulfatatoraceae/Thermodesulfatator_indicus
RS_GCF_000008625.1__O__Aquificaceae/Aquifex_aeolicus
RS_GCF_000482825.1__O__Thermodesulfovibrionaceae/Thermodesulfovibrio_islandicus
RS_GCF_000020145.1__O__Elusimicrobiaceae/Elusimicrobium_minutum
RS_GCF_000022565.1__O__Acidobacteriaceae/Acidobacterium_capsulatum
RS_GCF_001886815.1__O__Calditrichaceae/Caldithrix_abyssi
RS_GCF_000016085.1__O__Chlorobiaceae/Chlorobium_phaeovibrioides
RS_GCF_000146065.2__O__Cloacimonadaceae/Cloacimonas_acidaminovorans
RS_GCF_000010305.1__O__Gemmatimonadaceae/Gemmatimonas_aurantiaca
RS_GCF_000146505.1__O__Fibrobacteraceae/Fibrobacter_succinogenes
RS_GCF_000170755.1__O__Lentisphaeraceae/Lentisphaera_araneosa
RS_GCF_000172155.1__O__Verrucomicrobiaceae/Verrucomicrobium_spinosum
RS_GCF_000068585.1__O__Chlamydiaceae/Chlamydia_trachomatis
RS_GCF_000196115.1__O__Pirellulaceae/Rhodopirellula_baltica
RS_GCF_000604125.1__O__Treponemataceae/Treponema_pallidum
RS_GCF_000023325.1__O__Kosmotogaceae/Kosmotoga_A_olearia
RS_GCF_000237805.1__O__Dethiosulfovibrionaceae/Jonquetella_anthropi
RS_GCF_000284335.1__O__Caldisericaceae/Caldisericum_exile
RS_GCF_000020965.1__O__Dictyoglomaceae/Dictyoglomus_thermophilum
RS_GCF_000427095.1__O__Chthonomonadaceae/Chthonomonas_calidirosea
146.86
353.34
526.28
313.01
633.03
25.87
298.43
750.09
841.99
90.23
148.98
89.73
165.19
274.64
474.38
1266.45
274.51
687.75
1375.89
310.08
1152.7
49.12
70.82
1156.26
0.82
35.88
528.8
1509.13
690.05
0.46
776.04
310.88
609.63
1030.22
1158.01
802.55
1288.16
1634.17
1391.48
767.09
1011.12
100.81
107.81
315.71
188.83
355.98
440.82
1821.01
742.56
270.38
1003.73
45.57
735.81
367.32
2476.31
955.25
342.67
1098.37
522.15
3117.89
788.87
991.31
274.45
760.62
3240.85
1322.3
2981.6
1643.2
2303.95
2624.55
2230.06
2841.75
2177.41
3450.88
2969.47
2476.42
2571.59
2020.61
3006.23
2462.3
2745.01
2369.89
2921.85
2023.03
2427.04
3188.06
2681.24
2355.35
2618.18
2943.64
2408.23
3500.0
3000.0
2500.0
2000.0
1500.0
1000.0
500.0
0.0

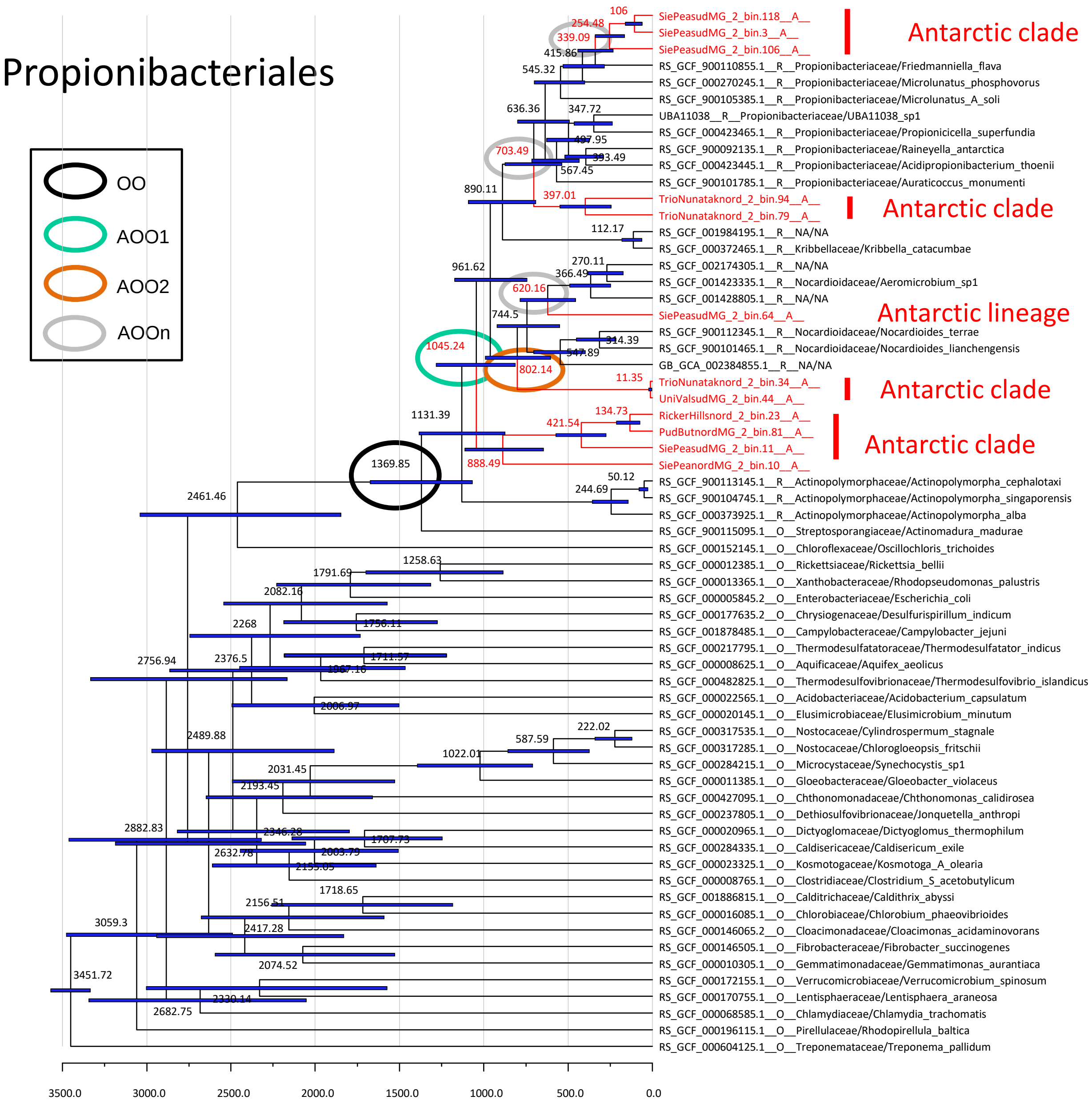
Propionibacteriales
OO
AOO1
AOO2
AOOn
Antarctic clade
Antarctic clade
Antarctic lineage
Antarctic clade
Antarctic clade
SiePeasudMG_2_bin.118__A__
SiePeasudMG_2_bin.3__A__
SiePeasudMG_2_bin.106__A__
RS_GCF_900110855.1__R__Propionibacteriaceae/Friedmanniella_flava
RS_GCF_000270245.1__R__Propionibacteriaceae/Microlunatus_phosphovorus
RS_GCF_900105385.1__R__Propionibacteriaceae/Microlunatus_A_soli
UBA11038__R__Propionibacteriaceae/UBA11038_sp1
RS_GCF_000423465.1__R__Propionibacteriaceae/Propionicicella_superfundia
RS_GCF_900092135.1__R__Propionibacteriaceae/Raineyella_antarctica
RS_GCF_000423445.1__R__Propionibacteriaceae/Acidipropionibacterium_thoenii
RS_GCF_900101785.1__R__Propionibacteriaceae/Auraticoccus_monumenti
TrioNunataknord_2_bin.94__A__
TrioNunataknord_2_bin.79__A__
RS_GCF_001984195.1__R__NA/NA
RS_GCF_000372465.1__R__Kribbellaceae/Kribbella_catacumbae
RS_GCF_002174305.1__R__NA/NA
RS_GCF_001423335.1__R__Nocardioidaceae/Aeromicrobium_sp1
RS_GCF_001428805.1__R__NA/NA
SiePeasudMG_2_bin.64__A__
RS_GCF_900112345.1__R__Nocardioidaceae/Nocardioides_terrae
RS_GCF_900101465.1__R__Nocardioidaceae/Nocardioides_lianchengensis
GB_GCA_002384855.1__R__NA/NA
TrioNunataknord_2_bin.34__A__
UniValsudMG_2_bin.44__A__
RickerHillsnord_2_bin.23__A__
PudButnordMG_2_bin.81__A__
SiePeasudMG_2_bin.11__A__
SiePeanordMG_2_bin.10__A__
RS_GCF_900113145.1__R__Actinopolymorphaceae/Actinopolymorpha_cephalotaxi
RS_GCF_900104745.1__R__Actinopolymorphaceae/Actinopolymorpha_singaporensis
RS_GCF_000373925.1__R__Actinopolymorphaceae/Actinopolymorpha_alba
RS_GCF_900115095.1__O__Streptosporangiaceae/Actinomadura_madurae
RS_GCF_000152145.1__O__Chloroflexaceae/Oscillochloris_trichoides
RS_GCF_000012385.1__O__Rickettsiaceae/Rickettsia_bellii
RS_GCF_000013365.1__O__Xanthobacteraceae/Rhodopseudomonas_palustris
RS_GCF_000005845.2__O__Enterobacteriaceae/Escherichia_coli
RS_GCF_000177635.2__O__Chrysiogenaceae/Desulfurispirillum_indicum
RS_GCF_001878485.1__O__Campylobacteraceae/Campylobacter_jejuni
RS_GCF_000217795.1__O__Thermodesulfatatoraceae/Thermodesulfatator_indicus
RS_GCF_000008625.1__O__Aquificaceae/Aquifex_aeolicus
RS_GCF_000482825.1__O__Thermodesulfovibrionaceae/Thermodesulfovibrio_islandicus
RS_GCF_000022565.1__O__Acidobacteriaceae/Acidobacterium_capsulatum
RS_GCF_000020145.1__O__Elusimicrobiaceae/Elusimicrobium_minutum
RS_GCF_000317535.1__O__Nostocaceae/Cylindrospermum_stagnale
RS_GCF_000317285.1__O__Nostocaceae/Chlorogloeopsis_fritschii
RS_GCF_000284215.1__O__Microcystaceae/Synechocystis_sp1
RS_GCF_000011385.1__O__Gloeobacteraceae/Gloeobacter_violaceus
RS_GCF_000427095.1__O__Chthonomonadaceae/Chthonomonas_calidirosea
RS_GCF_000237805.1__O__Dethiosulfovibrionaceae/Jonquetella_anthropi
RS_GCF_000020965.1__O__Dictyoglomaceae/Dictyoglomus_thermophilum
RS_GCF_000284335.1__O__Caldisericaceae/Caldisericum_exile
RS_GCF_000023325.1__O__Kosmotogaceae/Kosmotoga_A_olearia
RS_GCF_000008765.1__O__Clostridiaceae/Clostridium_S_acetobutylicum
RS_GCF_001886815.1__O__Calditrichaceae/Caldithrix_abyssi
RS_GCF_000016085.1__O__Chlorobiaceae/Chlorobium_phaeovibrioides
RS_GCF_000146065.2__O__Cloacimonadaceae/Cloacimonas_acidaminovorans
RS_GCF_000146505.1__O__Fibrobacteraceae/Fibrobacter_succinogenes
RS_GCF_000010305.1__O__Gemmatimonadaceae/Gemmatimonas_aurantiaca
RS_GCF_000172155.1__O__Verrucomicrobiaceae/Verrucomicrobium_spinosum
RS_GCF_000170755.1__O__Lentisphaeraceae/Lentisphaera_araneosa
RS_GCF_000068585.1__O__Chlamydiaceae/Chlamydia_trachomatis
RS_GCF_000196115.1__O__Pirellulaceae/Rhodopirellula_baltica
RS_GCF_000604125.1__O__Treponemataceae/Treponema_pallidum
3500.0
3000.0
2500.0
2000.0
1500.0
1000.0
500.0
0.0

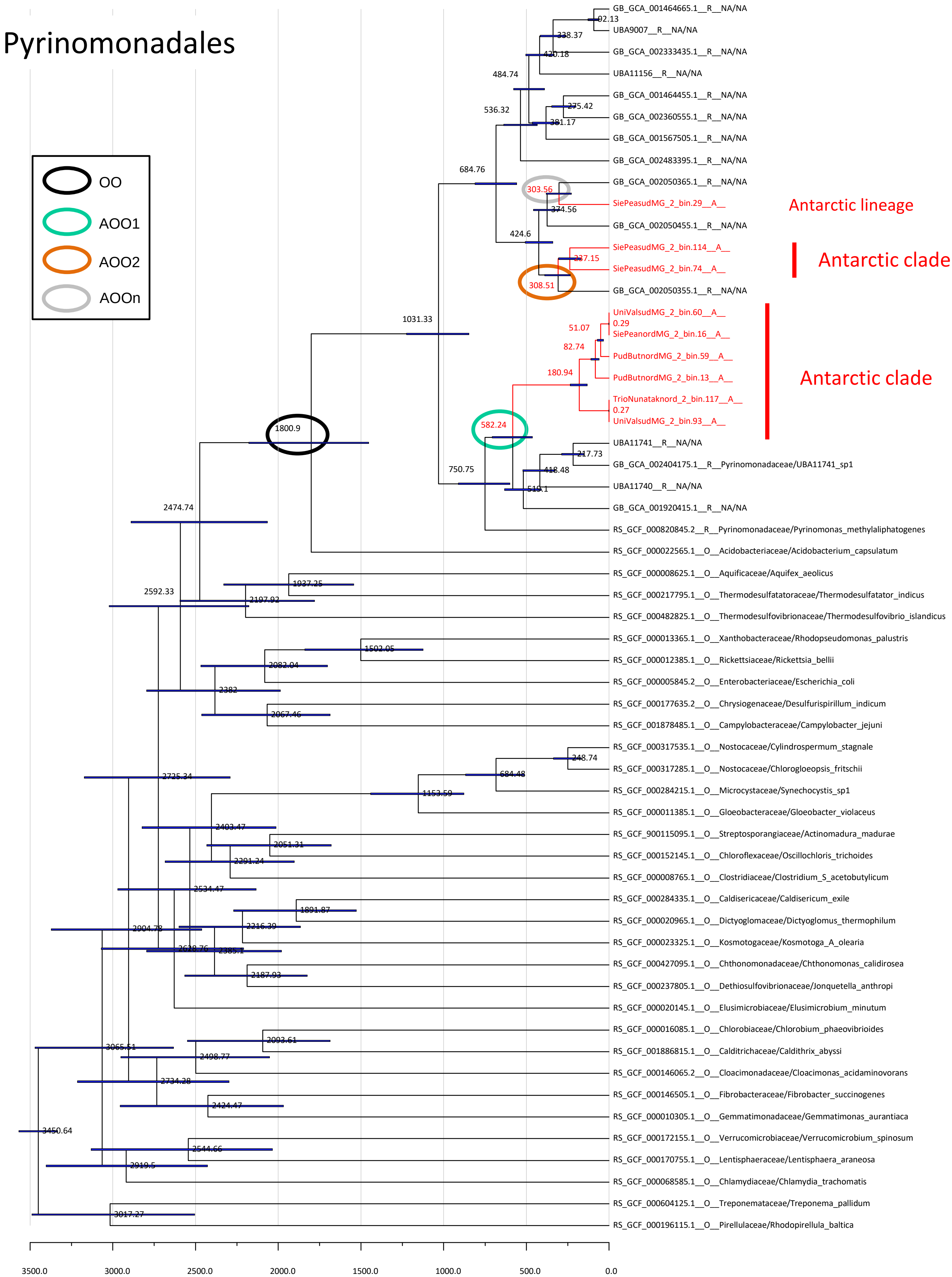

Pyrinomonadales
OO
AOO1
AOO2
AOOn
Antarctic lineage
Antarctic clade
Antarctic clade
GB_GCA_001464665.1__R__NA/NA
92.13
UBA9007__R__NA/NA
238.37
GB_GCA_002333435.1__R__NA/NA
420.18
UBA11156__R__NA/NA
484.74
GB_GCA_001464455.1__R__NA/NA
275.42
536.32
GB_GCA_002360555.1__R__NA/NA
381.17
GB_GCA_001567505.1__R__NA/NA
GB_GCA_002483395.1__R__NA/NA
684.76
GB_GCA_002050365.1__R__NA/NA
303.56
SiePeasudMG_2_bin.29__A__
374.56
GB_GCA_002050455.1__R__NA/NA
424.6
SiePeasudMG_2_bin.114__A__
237.15
SiePeasudMG_2_bin.74__A__
308.51
GB_GCA_002050355.1__R__NA/NA
1031.33
UniValsudMG_2_bin.60__A__
0.29
51.07
SiePeanordMG_2_bin.16__A__
82.74
PudButnordMG_2_bin.59__A__
180.94
PudButnordMG_2_bin.13__A__
TrioNunataknord_2_bin.117__A__
0.27
UniValsudMG_2_bin.93__A__
1800.9
582.24
UBA11741__R__NA/NA
217.73
GB_GCA_002404175.1__R__Pyrinomonadaceae/UBA11741_sp1
750.75
418.48
UBA11740__R__NA/NA
516.1
GB_GCA_001920415.1__R__NA/NA
2474.74
RS_GCF_000820845.2__R__Pyrinomonadaceae/Pyrinomonas_methylaliphatogenes
RS_GCF_000022565.1__O__Acidobacteriaceae/Acidobacterium_capsulatum
RS_GCF_000008625.1__O__Aquificaceae/Aquifex_aeolicus
1937.25
2592.33
2197.92
RS_GCF_000217795.1__O__Thermodesulfatatoraceae/Thermodesulfatator_indicus
RS_GCF_000482825.1__O__Thermodesulfovibrionaceae/Thermodesulfovibrio_islandicus
RS_GCF_000013365.1__O__Xanthobacteraceae/Rhodopseudomonas_palustris
1502.05
2082.04
RS_GCF_000012385.1__O__Rickettsiaceae/Rickettsia_bellii
RS_GCF_000005845.2__O__Enterobacteriaceae/Escherichia_coli
2382
RS_GCF_000177635.2__O__Chrysiogenaceae/Desulfurispirillum_indicum
2067.46
RS_GCF_001878485.1__O__Campylobacteraceae/Campylobacter_jejuni
RS_GCF_000317535.1__O__Nostocaceae/Cylindrospermum_stagnale
248.74
RS_GCF_000317285.1__O__Nostocaceae/Chlorogloeopsis_fritschii
2725.24
684.48
RS_GCF_000284215.1__O__Microcystaceae/Synechocystis_sp1
1153.59
RS_GCF_000011385.1__O__Gloeobacteraceae/Gloeobacter_violaceus
2403.47
RS_GCF_900115095.1__O__Streptosporangiaceae/Actinomadura_madurae
2051.31
RS_GCF_000152145.1__O__Chloroflexaceae/Oscillochloris_trichoides
2291.24
RS_GCF_000008765.1__O__Clostridiaceae/Clostridium_S_acetobutylicum
2534.47
RS_GCF_000284335.1__O__Caldisericaceae/Caldisericum_exile
1891.87
RS_GCF_000020965.1__O__Dictyoglomaceae/Dictyoglomus_thermophilum
2904.78
2216.39
RS_GCF_000023325.1__O__Kosmotogaceae/Kosmotoga_A_olearia
2628.76
2385.1
RS_GCF_000427095.1__O__Chthonomonadaceae/Chthonomonas_calidirosea
2187.93
RS_GCF_000237805.1__O__Dethiosulfovibrionaceae/Jonquetella_anthropi
RS_GCF_000020145.1__O__Elusimicrobiaceae/Elusimicrobium_minutum
RS_GCF_000016085.1__O__Chlorobiaceae/Chlorobium_phaeovibrioides
3065.51
2093.61
2498.77
RS_GCF_001886815.1__O__Calditrichaceae/Caldithrix_abyssi
RS_GCF_000146065.2__O__Cloacimonadaceae/Cloacimonas_acidaminovorans
2734.28
RS_GCF_000146505.1__O__Fibrobacteraceae/Fibrobacter_succinogenes
2424.47
RS_GCF_000010305.1__O__Gemmatimonadaceae/Gemmatimonas_aurantiaca
3450.64
RS_GCF_000172155.1__O__Verrucomicrobiaceae/Verrucomicrobium_spinosum
2544.66
2919.5
RS_GCF_000170755.1__O__Lentisphaeraceae/Lentisphaera_araneosa
RS_GCF_000068585.1__O__Chlamydiaceae/Chlamydia_trachomatis
RS_GCF_000604125.1__O__Treponemataceae/Treponema_pallidum
3017.27
RS_GCF_000196115.1__O__Pirellulaceae/Rhodopirellula_baltica
3500.0
3000.0
2500.0
2000.0
1500.0
1000.0
500.0
0.0

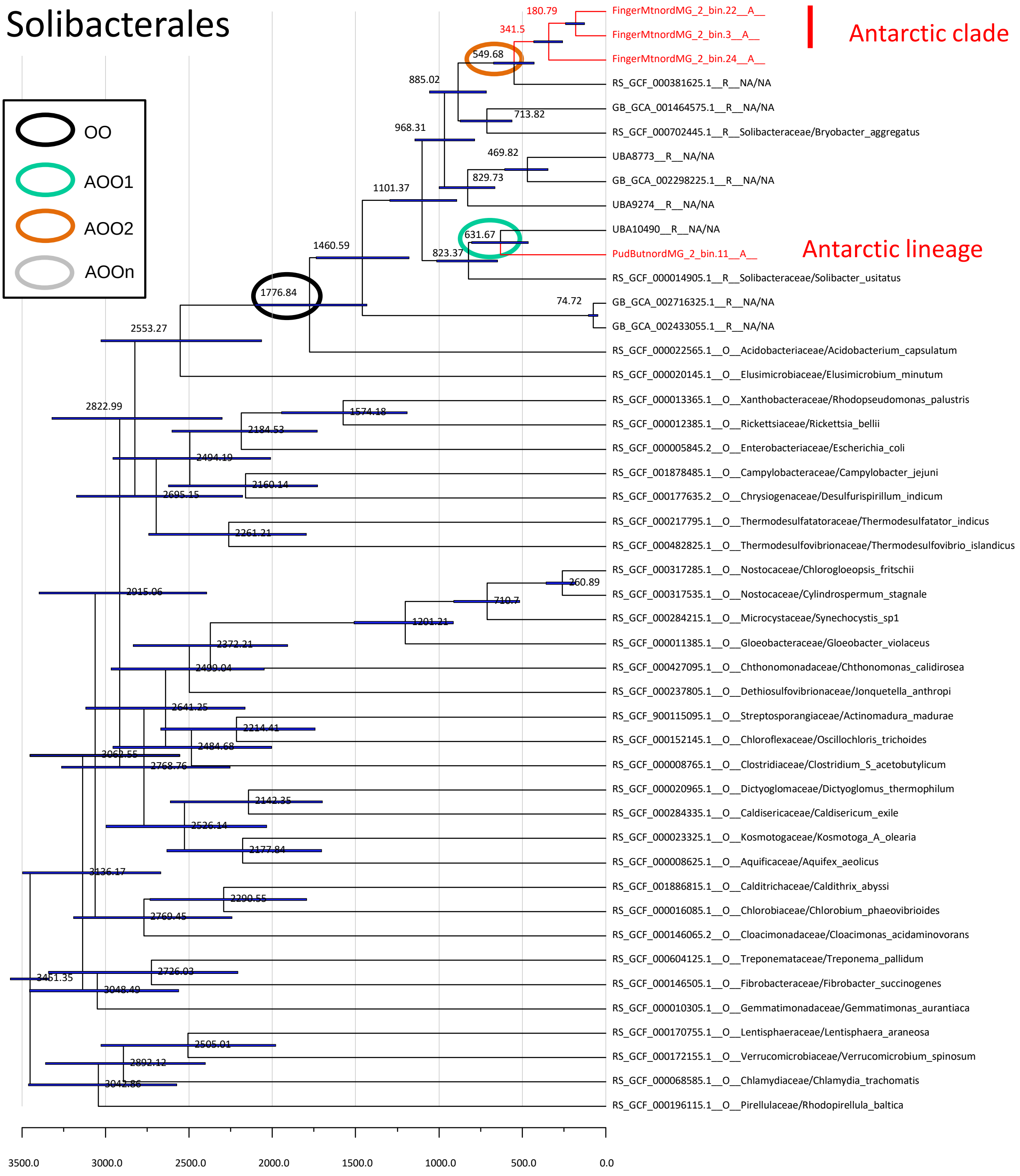

Solibacterales
OO
AOO1
AOO2
AOOn
Antarctic clade
Antarctic lineage
FingerMtnordMG_2_bin.22__A__
FingerMtnordMG_2_bin.3__A__
FingerMtnordMG_2_bin.24__A__
RS_GCF_000381625.1__R__NA/NA
GB_GCA_001464575.1__R__NA/NA
RS_GCF_000702445.1__R__Solibacteraceae/Bryobacter_aggregatus
UBA8773__R__NA/NA
GB_GCA_002298225.1__R__NA/NA
UBA9274__R__NA/NA
UBA10490__R__NA/NA
PudButnordMG_2_bin.11__A__
RS_GCF_000014905.1__R__Solibacteraceae/Solibacter_usitatus
GB_GCA_002716325.1__R__NA/NA
GB_GCA_002433055.1__R__NA/NA
RS_GCF_000022565.1__O__Acidobacteriaceae/Acidobacterium_capsulatum
RS_GCF_000020145.1__O__Elusimicrobiaceae/Elusimicrobium_minutum
RS_GCF_000013365.1__O__Xanthobacteraceae/Rhodopseudomonas_palustris
RS_GCF_000012385.1__O__Rickettsiaceae/Rickettsia_bellii
RS_GCF_000005845.2__O__Enterobacteriaceae/Escherichia_coli
RS_GCF_001878485.1__O__Campylobacteraceae/Campylobacter_jejuni
RS_GCF_000177635.2__O__Chrysiogenaceae/Desulfurispirillum_indicum
RS_GCF_000217795.1__O__Thermodesulfatatoraceae/Thermodesulfatator_indicus
RS_GCF_000482825.1__O__Thermodesulfovibrionaceae/Thermodesulfovibrio_islandicus
RS_GCF_000317285.1__O__Nostocaceae/Chlorogloeopsis_fritschii
RS_GCF_000317535.1__O__Nostocaceae/Cylindrospermum_stagnale
RS_GCF_000284215.1__O__Microcystaceae/Synechocystis_sp1
RS_GCF_000011385.1__O__Gloeobacteraceae/Gloeobacter_violaceus
RS_GCF_000427095.1__O__Chthonomonadaceae/Chthonomonas_calidirosea
RS_GCF_000237805.1__O__Dethiosulfovibrionaceae/Jonquetella_anthropi
RS_GCF_900115095.1__O__Streptosporangiaceae/Actinomadura_madurae
RS_GCF_000152145.1__O__Chloroflexaceae/Oscillochloris_trichoides
RS_GCF_000008765.1__O__Clostridiaceae/Clostridium_S_acetobutylicum
RS_GCF_000020965.1__O__Dictyoglomaceae/Dictyoglomus_thermophilum
RS_GCF_000284335.1__O__Caldisericaceae/Caldisericum_exile
RS_GCF_000023325.1__O__Kosmotogaceae/Kosmotoga_A_olearia
RS_GCF_000008625.1__O__Aquificaceae/Aquifex_aeolicus
RS_GCF_001886815.1__O__Calditrichaceae/Caldithrix_abyssi
RS_GCF_000016085.1__O__Chlorobiaceae/Chlorobium_phaeovibrioides
RS_GCF_000146065.2__O__Cloacimonadaceae/Cloacimonas_acidaminovorans
RS_GCF_000604125.1__O__Treponemataceae/Treponema_pallidum
RS_GCF_000146505.1__O__Fibrobacteraceae/Fibrobacter_succinogenes
RS_GCF_000010305.1__O__Gemmatimonadaceae/Gemmatimonas_aurantiaca
RS_GCF_000170755.1__O__Lentisphaeraceae/Lentisphaera_araneosa
RS_GCF_000172155.1__O__Verrucomicrobiaceae/Verrucomicrobium_spinosum
RS_GCF_000068585.1__O__Chlamydiaceae/Chlamydia_trachomatis
RS_GCF_000196115.1__O__Pirellulaceae/Rhodopirellula_baltica
180.79
341.5
549.68
885.02
713.82
968.31
469.82
829.73
1101.37
631.67
823.37
1460.59
1776.84
74.72
2553.27
2822.99
1574.18
2184.53
2494.19
2160.14
2695.15
2261.21
260.89
2915.06
710.7
1201.21
2372.21
2499.04
2641.25
2214.41
2484.68
3062.55
2768.76
2142.35
2526.14
2177.84
3136.17
2290.55
2769.45
2726.03
3451.35
3048.49
2505.01
2892.12
3042.86
3500.0
3000.0
2500.0
2000.0
1500.0
1000.0
500.0
0.0

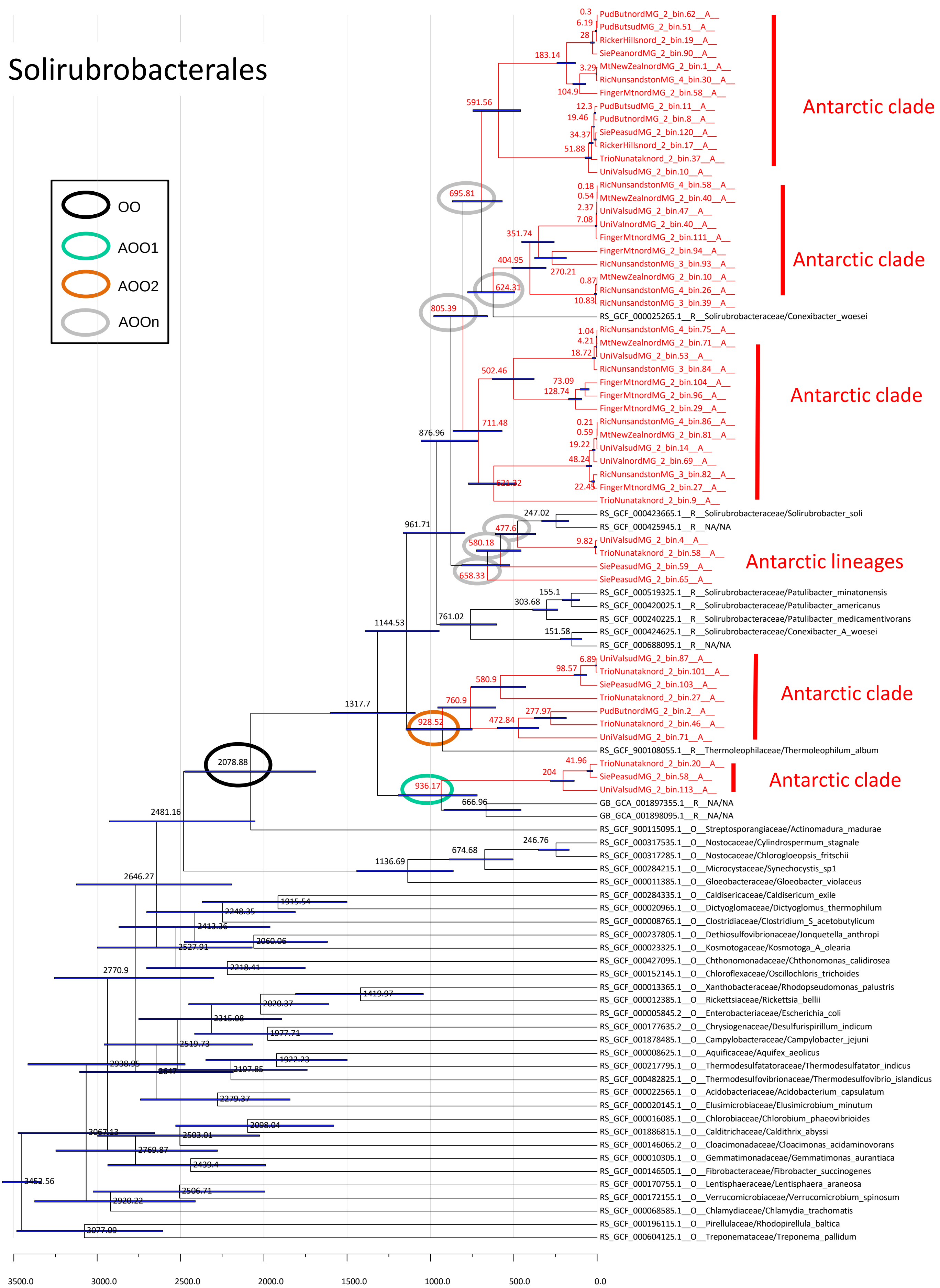

Solirubrobacterales
OO
AOO1
AOO2
AOOn
Antarctic clade
Antarctic clade
Antarctic clade
Antarctic lineages
Antarctic clade
Antarctic clade
PudButnordMG_2_bin.62__A__
PudButsudMG_2_bin.51__A__
RickerHillsnord_2_bin.19__A__
SiePeanordMG_2_bin.90__A__
MtNewZealnordMG_2_bin.1__A__
RicNunsandstonMG_4_bin.30__A__
FingerMtnordMG_2_bin.58__A__
PudButsudMG_2_bin.11__A__
PudButnordMG_2_bin.8__A__
SiePeasudMG_2_bin.120__A__
RickerHillsnord_2_bin.17__A__
TrioNunataknord_2_bin.37__A__
UniValsudMG_2_bin.10__A__
RicNunsandstonMG_4_bin.58__A__
MtNewZealnordMG_2_bin.40__A__
UniValsudMG_2_bin.47__A__
UniValnordMG_2_bin.40__A__
FingerMtnordMG_2_bin.111__A__
FingerMtnordMG_2_bin.94__A__
RicNunsandstonMG_3_bin.93__A__
MtNewZealnordMG_2_bin.10__A__
RicNunsandstonMG_4_bin.26__A__
RicNunsandstonMG_3_bin.39__A__
RS_GCF_000025265.1__R__Solirubrobacteraceae/Conexibacter_woesei
RicNunsandstonMG_4_bin.75__A__
MtNewZealnordMG_2_bin.71__A__
UniValsudMG_2_bin.53__A__
RicNunsandstonMG_3_bin.84__A__
FingerMtnordMG_2_bin.104__A__
FingerMtnordMG_2_bin.96__A__
FingerMtnordMG_2_bin.29__A__
RicNunsandstonMG_4_bin.86__A__
MtNewZealnordMG_2_bin.81__A__
UniValsudMG_2_bin.14__A__
UniValnordMG_2_bin.69__A__
RicNunsandstonMG_3_bin.82__A__
FingerMtnordMG_2_bin.27__A__
TrioNunataknord_2_bin.9__A__
RS_GCF_000423665.1__R__Solirubrobacteraceae/Solirubrobacter_soli
RS_GCF_000425945.1__R__NA/NA
UniValsudMG_2_bin.4__A__
TrioNunataknord_2_bin.58__A__
SiePeasudMG_2_bin.59__A__
SiePeasudMG_2_bin.65__A__
RS_GCF_000519325.1__R__Solirubrobacteraceae/Patulibacter_minatonensis
RS_GCF_000420025.1__R__Solirubrobacteraceae/Patulibacter_americanus
RS_GCF_000240225.1__R__Solirubrobacteraceae/Patulibacter_medicamentivorans
RS_GCF_000424625.1__R__Solirubrobacteraceae/Conexibacter_A_woesei
RS_GCF_000688095.1__R__NA/NA
UniValsudMG_2_bin.87__A__
TrioNunataknord_2_bin.101__A__
SiePeasudMG_2_bin.103__A__
TrioNunataknord_2_bin.27__A__
PudButnordMG_2_bin.2__A__
TrioNunataknord_2_bin.46__A__
UniValsudMG_2_bin.71__A__
RS_GCF_900108055.1__R__Thermoleophilaceae/Thermoleophilum_album
TrioNunataknord_2_bin.20__A__
SiePeasudMG_2_bin.58__A__
UniValsudMG_2_bin.113__A__
GB_GCA_001897355.1__R__NA/NA
GB_GCA_001898095.1__R__NA/NA
RS_GCF_900115095.1__O__Streptosporangiaceae/Actinomadura_madurae
RS_GCF_000317535.1__O__Nostocaceae/Cylindrospermum_stagnale
RS_GCF_000317285.1__O__Nostocaceae/Chlorogloeopsis_fritschii
RS_GCF_000284215.1__O__Microcystaceae/Synechocystis_sp1
RS_GCF_000011385.1__O__Gloeobacteraceae/Gloeobacter_violaceus
RS_GCF_000284335.1__O__Caldisericaceae/Caldisericum_exile
RS_GCF_000020965.1__O__Dictyoglomaceae/Dictyoglomus_thermophilum
RS_GCF_000008765.1__O__Clostridiaceae/Clostridium_S_acetobutylicum
RS_GCF_000237805.1__O__Dethiosulfovibrionaceae/Jonquetella_anthropi
RS_GCF_000023325.1__O__Kosmotogaceae/Kosmotoga_A_olearia
RS_GCF_000427095.1__O__Chthonomonadaceae/Chthonomonas_calidirosea
RS_GCF_000152145.1__O__Chloroflexaceae/Oscillochloris_trichoides
RS_GCF_000013365.1__O__Xanthobacteraceae/Rhodopseudomonas_palustris
RS_GCF_000012385.1__O__Rickettsiaceae/Rickettsia_bellii
RS_GCF_000005845.2__O__Enterobacteriaceae/Escherichia_coli
RS_GCF_000177635.2__O__Chrysiogenaceae/Desulfurispirillum_indicum
RS_GCF_001878485.1__O__Campylobacteraceae/Campylobacter_jejuni
RS_GCF_000008625.1__O__Aquificaceae/Aquifex_aeolicus
RS_GCF_000217795.1__O__Thermodesulfatatoraceae/Thermodesulfatator_indicus
RS_GCF_000482825.1__O__Thermodesulfovibrionaceae/Thermodesulfovibrio_islandicus
RS_GCF_000022565.1__O__Acidobacteriaceae/Acidobacterium_capsulatum
RS_GCF_000020145.1__O__Elusimicrobiaceae/Elusimicrobium_minutum
RS_GCF_000016085.1__O__Chlorobiaceae/Chlorobium_phaeovibrioides
RS_GCF_001886815.1__O__Calditrichaceae/Caldithrix_abyssi
RS_GCF_000146065.2__O__Cloacimonadaceae/Cloacimonas_acidaminovorans
RS_GCF_000010305.1__O__Gemmatimonadaceae/Gemmatimonas_aurantiaca
RS_GCF_000146505.1__O__Fibrobacteraceae/Fibrobacter_succinogenes
RS_GCF_000170755.1__O__Lentisphaeraceae/Lentisphaera_araneosa
RS_GCF_000172155.1__O__Verrucomicrobiaceae/Verrucomicrobium_spinosum
RS_GCF_000068585.1__O__Chlamydiaceae/Chlamydia_trachomatis
RS_GCF_000196115.1__O__Pirellulaceae/Rhodopirellula_baltica
RS_GCF_000604125.1__O__Treponemataceae/Treponema_pallidum
3500.0
3000.0
2500.0
2000.0
1500.0
1000.0
500.0
0.0

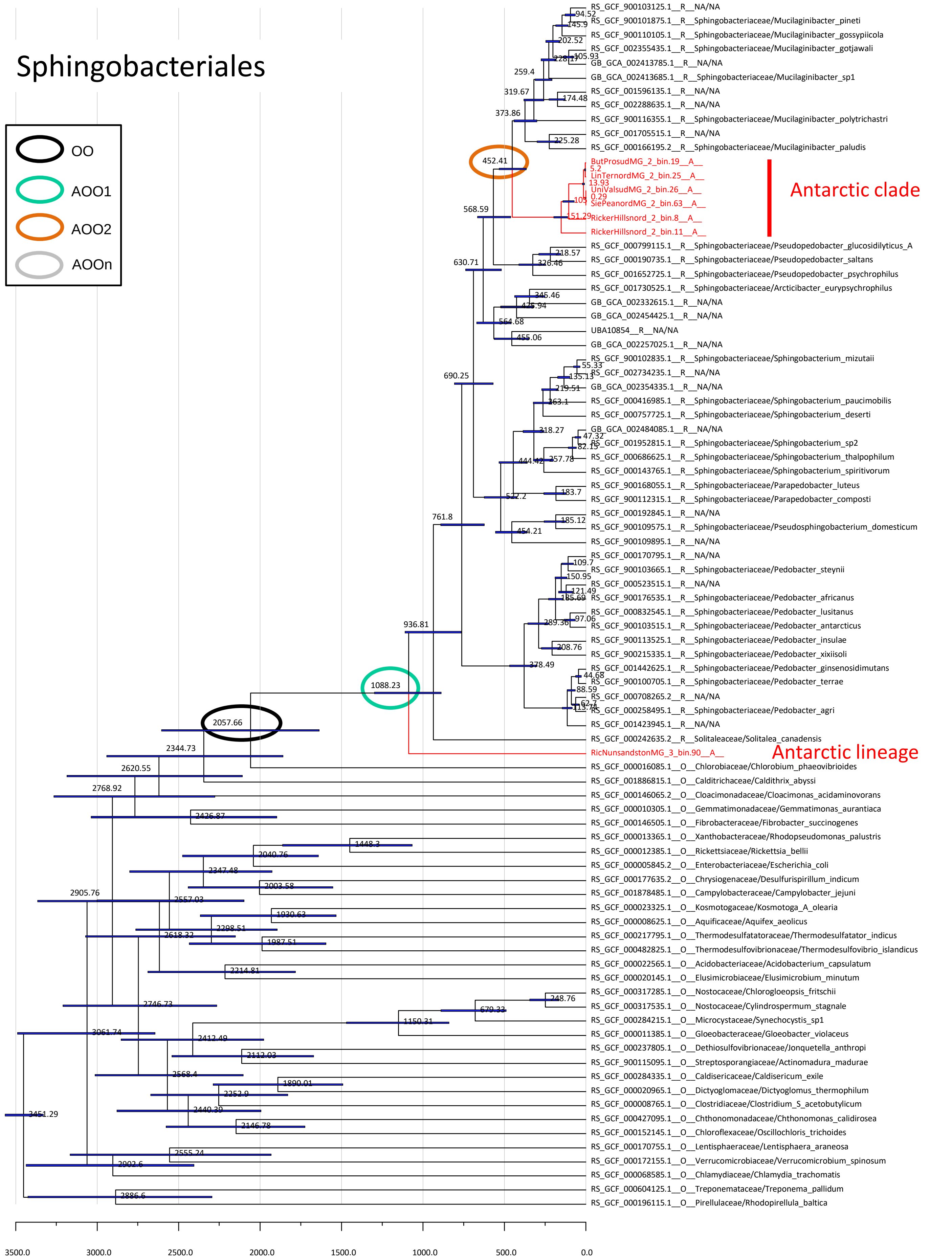

Sphingobacteriales
OO
AOO1
AOO2
AOOn
Antarctic clade
Antarctic lineage
452.41
1088.23
2057.66
RS_GCF_900103125.1__R__NA/NA
RS_GCF_900101875.1__R__Sphingobacteriaceae/Mucilaginibacter_pineti
RS_GCF_900110105.1__R__Sphingobacteriaceae/Mucilaginibacter_gossypiicola
RS_GCF_002355435.1__R__Sphingobacteriaceae/Mucilaginibacter_gotjawali
GB_GCA_002413785.1__R__NA/NA
GB_GCA_002413685.1__R__Sphingobacteriaceae/Mucilaginibacter_sp1
RS_GCF_001596135.1__R__NA/NA
RS_GCF_002288635.1__R__NA/NA
RS_GCF_900116355.1__R__Sphingobacteriaceae/Mucilaginibacter_polytrichastri
RS_GCF_001705515.1__R__NA/NA
RS_GCF_000166195.2__R__Sphingobacteriaceae/Mucilaginibacter_paludis
ButProsudMG_2_bin.19__A__
LinTernordMG_2_bin.25__A__
UniValsudMG_2_bin.26__A__
SiePeanordMG_2_bin.63__A__
RickerHillsnord_2_bin.8__A__
RickerHillsnord_2_bin.11__A__
RS_GCF_000799115.1__R__Sphingobacteriaceae/Pseudopedobacter_glucosidilyticus_A
RS_GCF_000190735.1__R__Sphingobacteriaceae/Pseudopedobacter_saltans
RS_GCF_001652725.1__R__Sphingobacteriaceae/Pseudopedobacter_psychrophilus
RS_GCF_001730525.1__R__Sphingobacteriaceae/Arcticibacter_eurypsychrophilus
GB_GCA_002332615.1__R__NA/NA
GB_GCA_002454425.1__R__NA/NA
UBA10854__R__NA/NA
GB_GCA_002257025.1__R__NA/NA
RS_GCF_900102835.1__R__Sphingobacteriaceae/Sphingobacterium_mizutaii
RS_GCF_002734235.1__R__NA/NA
GB_GCA_002354335.1__R__NA/NA
RS_GCF_000416985.1__R__Sphingobacteriaceae/Sphingobacterium_paucimobilis
RS_GCF_000757725.1__R__Sphingobacteriaceae/Sphingobacterium_deserti
GB_GCA_002484085.1__R__NA/NA
RS_GCF_001952815.1__R__Sphingobacteriaceae/Sphingobacterium_sp2
RS_GCF_000686625.1__R__Sphingobacteriaceae/Sphingobacterium_thalpophilum
RS_GCF_000143765.1__R__Sphingobacteriaceae/Sphingobacterium_spiritivorum
RS_GCF_900168055.1__R__Sphingobacteriaceae/Parapedobacter_luteus
RS_GCF_900112315.1__R__Sphingobacteriaceae/Parapedobacter_composti
RS_GCF_000192845.1__R__NA/NA
RS_GCF_900109575.1__R__Sphingobacteriaceae/Pseudosphingobacterium_domesticum
RS_GCF_900109895.1__R__NA/NA
RS_GCF_000170795.1__R__NA/NA
RS_GCF_900103665.1__R__Sphingobacteriaceae/Pedobacter_steynii
RS_GCF_000523515.1__R__NA/NA
RS_GCF_900176535.1__R__Sphingobacteriaceae/Pedobacter_africanus
RS_GCF_000832545.1__R__Sphingobacteriaceae/Pedobacter_lusitanus
RS_GCF_900103515.1__R__Sphingobacteriaceae/Pedobacter_antarcticus
RS_GCF_900113525.1__R__Sphingobacteriaceae/Pedobacter_insulae
RS_GCF_900215335.1__R__Sphingobacteriaceae/Pedobacter_xixiisoli
RS_GCF_001442625.1__R__Sphingobacteriaceae/Pedobacter_ginsenosidimutans
RS_GCF_900100705.1__R__Sphingobacteriaceae/Pedobacter_terrae
RS_GCF_000708265.2__R__NA/NA
RS_GCF_000258495.1__R__Sphingobacteriaceae/Pedobacter_agri
RS_GCF_001423945.1__R__NA/NA
RS_GCF_000242635.2__R__Solitaleaceae/Solitalea_canadensis
RicNunsandstonMG_3_bin.90__A__
RS_GCF_000016085.1__O__Chlorobiaceae/Chlorobium_phaeovibrioides
RS_GCF_001886815.1__O__Calditrichaceae/Caldithrix_abyssi
RS_GCF_000146065.2__O__Cloacimonadaceae/Cloacimonas_acidaminovorans
RS_GCF_000010305.1__O__Gemmatimonadaceae/Gemmatimonas_aurantiaca
RS_GCF_000146505.1__O__Fibrobacteraceae/Fibrobacter_succinogenes
RS_GCF_000013365.1__O__Xanthobacteraceae/Rhodopseudomonas_palustris
RS_GCF_000012385.1__O__Rickettsiaceae/Rickettsia_bellii
RS_GCF_000005845.2__O__Enterobacteriaceae/Escherichia_coli
RS_GCF_000177635.2__O__Chrysiogenaceae/Desulfurispirillum_indicum
RS_GCF_001878485.1__O__Campylobacteraceae/Campylobacter_jejuni
RS_GCF_000023325.1__O__Kosmotogaceae/Kosmotoga_A_olearia
RS_GCF_000008625.1__O__Aquificaceae/Aquifex_aeolicus
RS_GCF_000217795.1__O__Thermodesulfatatoraceae/Thermodesulfatator_indicus
RS_GCF_000482825.1__O__Thermodesulfovibrionaceae/Thermodesulfovibrio_islandicus
RS_GCF_000022565.1__O__Acidobacteriaceae/Acidobacterium_capsulatum
RS_GCF_000020145.1__O__Elusimicrobiaceae/Elusimicrobium_minutum
RS_GCF_000317285.1__O__Nostocaceae/Chlorogloeopsis_fritschii
RS_GCF_000317535.1__O__Nostocaceae/Cylindrospermum_stagnale
RS_GCF_000284215.1__O__Microcystaceae/Synechocystis_sp1
RS_GCF_000011385.1__O__Gloeobacteraceae/Gloeobacter_violaceus
RS_GCF_000237805.1__O__Dethiosulfovibrionaceae/Jonquetella_anthropi
RS_GCF_900115095.1__O__Streptosporangiaceae/Actinomadura_madurae
RS_GCF_000284335.1__O__Caldisericaceae/Caldisericum_exile
RS_GCF_000020965.1__O__Dictyoglomaceae/Dictyoglomus_thermophilum
RS_GCF_000008765.1__O__Clostridiaceae/Clostridium_S_acetobutylicum
RS_GCF_000427095.1__O__Chthonomonadaceae/Chthonomonas_calidirosea
RS_GCF_000152145.1__O__Chloroflexaceae/Oscillochloris_trichoides
RS_GCF_000170755.1__O__Lentisphaeraceae/Lentisphaera_araneosa
RS_GCF_000172155.1__O__Verrucomicrobiaceae/Verrucomicrobium_spinosum
RS_GCF_000068585.1__O__Chlamydiaceae/Chlamydia_trachomatis
RS_GCF_000604125.1__O__Treponemataceae/Treponema_pallidum
RS_GCF_000196115.1__O__Pirellulaceae/Rhodopirellula_baltica
3500.0
3000.0
2500.0
2000.0
1500.0
1000.0
500.0
0.0

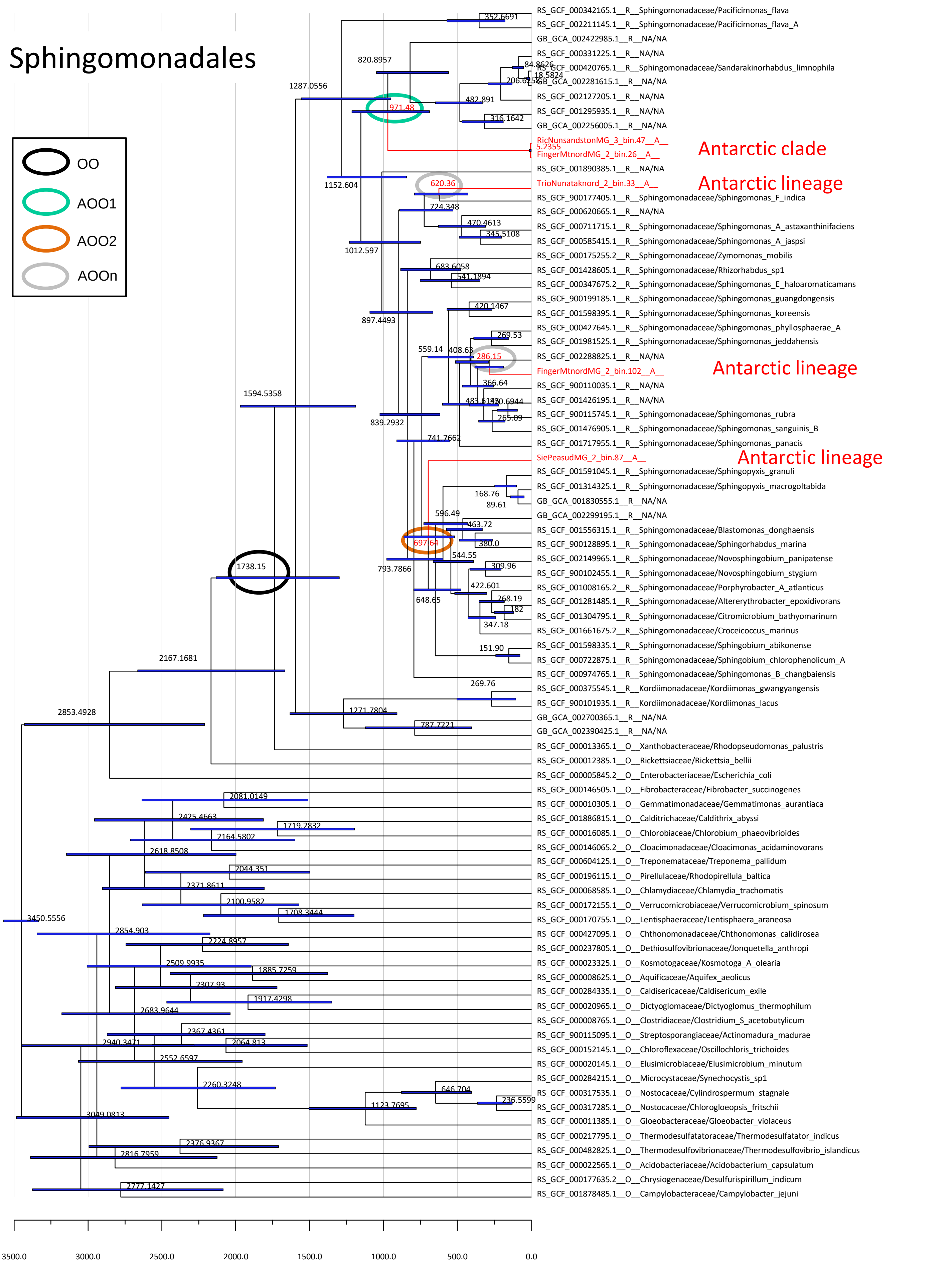
Sphingomonadales
OO
AOO1
AOO2
AOOn
Antarctic clade
Antarctic lineage
Antarctic lineage
Antarctic lineage
RS_GCF_000342165.1__R__Sphingomonadaceae/Pacificimonas_flava
RS_GCF_002211145.1__R__Sphingomonadaceae/Pacificimonas_flava_A
GB_GCA_002422985.1__R__NA/NA
RS_GCF_000331225.1__R__NA/NA
RS_GCF_000420765.1__R__Sphingomonadaceae/Sandarakinorhabdus_limnophila
GB_GCA_002281615.1__R__NA/NA
RS_GCF_002127205.1__R__NA/NA
RS_GCF_001295935.1__R__NA/NA
GB_GCA_002256005.1__R__NA/NA
RicNunsandstonMG_3_bin.47__A__
FingerMtnordMG_2_bin.26__A__
RS_GCF_001890385.1__R__NA/NA
TrioNunataknord_2_bin.33__A__
RS_GCF_900177405.1__R__Sphingomonadaceae/Sphingomonas_F_indica
RS_GCF_000620665.1__R__NA/NA
RS_GCF_000711715.1__R__Sphingomonadaceae/Sphingomonas_A_astaxanthinifaciens
RS_GCF_000585415.1__R__Sphingomonadaceae/Sphingomonas_A_jaspsi
RS_GCF_000175255.2__R__Sphingomonadaceae/Zymomonas_mobilis
RS_GCF_001428605.1__R__Sphingomonadaceae/Rhizorhabdus_sp1
RS_GCF_000347675.2__R__Sphingomonadaceae/Sphingomonas_E_haloaromaticamans
RS_GCF_900199185.1__R__Sphingomonadaceae/Sphingomonas_guangdongensis
RS_GCF_001598395.1__R__Sphingomonadaceae/Sphingomonas_koreensis
RS_GCF_000427645.1__R__Sphingomonadaceae/Sphingomonas_phyllosphaerae_A
RS_GCF_001981525.1__R__Sphingomonadaceae/Sphingomonas_jeddahensis
RS_GCF_002288825.1__R__NA/NA
FingerMtnordMG_2_bin.102__A__
RS_GCF_900110035.1__R__NA/NA
RS_GCF_001426195.1__R__NA/NA
RS_GCF_900115745.1__R__Sphingomonadaceae/Sphingomonas_rubra
RS_GCF_001476905.1__R__Sphingomonadaceae/Sphingomonas_sanguinis_B
RS_GCF_001717955.1__R__Sphingomonadaceae/Sphingomonas_panacis
SiePeasudMG_2_bin.87__A__
RS_GCF_001591045.1__R__Sphingomonadaceae/Sphingopyxis_granuli
RS_GCF_001314325.1__R__Sphingomonadaceae/Sphingopyxis_macrogoltabida
GB_GCA_001830555.1__R__NA/NA
GB_GCA_002299195.1__R__NA/NA
RS_GCF_001556315.1__R__Sphingomonadaceae/Blastomonas_donghaensis
RS_GCF_900128895.1__R__Sphingomonadaceae/Sphingorhabdus_marina
RS_GCF_002149965.1__R__Sphingomonadaceae/Novosphingobium_panipatense
RS_GCF_900102455.1__R__Sphingomonadaceae/Novosphingobium_stygium
RS_GCF_001008165.2__R__Sphingomonadaceae/Porphyrobacter_A_atlanticus
RS_GCF_001281485.1__R__Sphingomonadaceae/Altererythrobacter_epoxidivorans
RS_GCF_001304795.1__R__Sphingomonadaceae/Citromicrobium_bathyomarinum
RS_GCF_001661675.2__R__Sphingomonadaceae/Croceicoccus_marinus
RS_GCF_001598335.1__R__Sphingomonadaceae/Sphingobium_abikonense
RS_GCF_000722875.1__R__Sphingomonadaceae/Sphingobium_chlorophenolicum_A
RS_GCF_000974765.1__R__Sphingomonadaceae/Sphingomonas_B_changbaiensis
RS_GCF_000375545.1__R__Kordiimonadaceae/Kordiimonas_gwangyangensis
RS_GCF_900101935.1__R__Kordiimonadaceae/Kordiimonas_lacus
GB_GCA_002700365.1__R__NA/NA
GB_GCA_002390425.1__R__NA/NA
RS_GCF_000013365.1__O__Xanthobacteraceae/Rhodopseudomonas_palustris
RS_GCF_000012385.1__O__Rickettsiaceae/Rickettsia_bellii
RS_GCF_000005845.2__O__Enterobacteriaceae/Escherichia_coli
RS_GCF_000146505.1__O__Fibrobacteraceae/Fibrobacter_succinogenes
RS_GCF_000010305.1__O__Gemmatimonadaceae/Gemmatimonas_aurantiaca
RS_GCF_001886815.1__O__Caldithrichaceae/Caldithrix_abyssi
RS_GCF_000016085.1__O__Chlorobiaceae/Chlorobium_phaeovibrioides
RS_GCF_000146065.2__O__Cloacimonadaceae/Cloacimonas_acidaminovorans
RS_GCF_000604125.1__O__Treponemataceae/Treponema_pallidum
RS_GCF_000196115.1__O__Pirellulaceae/Rhodopirellula_baltica
RS_GCF_000068585.1__O__Chlamydiaceae/Chlamydia_trachomatis
RS_GCF_000172155.1__O__Verrucomicrobiaceae/Verrucomicrobium_spinosum
RS_GCF_000170755.1__O__Lentisphaeraceae/Lentisphaera_araneosa
RS_GCF_000427095.1__O__Chthonomonadaceae/Chthonomonas_calidirosea
RS_GCF_000237805.1__O__Dethiosulfovibrionaceae/Jonquetella_anthropi
RS_GCF_000023325.1__O__Kosmotogaceae/Kosmotoga_A_olearia
RS_GCF_000008625.1__O__Aquificaceae/Aquifex_aeolicus
RS_GCF_000284335.1__O__Caldisericaceae/Caldisericum_exile
RS_GCF_000020965.1__O__Dictyoglomaceae/Dictyoglomus_thermophilum
RS_GCF_000008765.1__O__Clostridiaceae/Clostridium_S_acetobutylicum
RS_GCF_900115095.1__O__Streptosporangiaceae/Actinomadura_madurae
RS_GCF_000152145.1__O__Chloroflexaceae/Oscillochloris_trichoides
RS_GCF_000020145.1__O__Elusimicrobiaceae/Elusimicrobium_minutum
RS_GCF_000284215.1__O__Microcystaceae/Synechocystis_sp1
RS_GCF_000317535.1__O__Nostocaceae/Cylindrospermum_stagnale
RS_GCF_000317285.1__O__Nostocaceae/Chlorogloeopsis_fritschii
RS_GCF_000011385.1__O__Gloeobacteraceae/Gloeobacter_violaceus
RS_GCF_000217795.1__O__Thermodesulfatatoraceae/Thermodesulfatator_indicus
RS_GCF_000482825.1__O__Thermodesulfovibrionaceae/Thermodesulfovibrio_islandicus
RS_GCF_000022565.1__O__Acidobacteriaceae/Acidobacterium_capsulatum
RS_GCF_000177635.2__O__Chrysiogenaceae/Desulfurispirillum_indicum
RS_GCF_001878485.1__O__Campylobacteraceae/Campylobacter_jejuni
971.48
620.36
286.15
697.64
1738.15
3500.0
3000.0
2500.0
2000.0
1500.0
1000.0
500.0
0.0

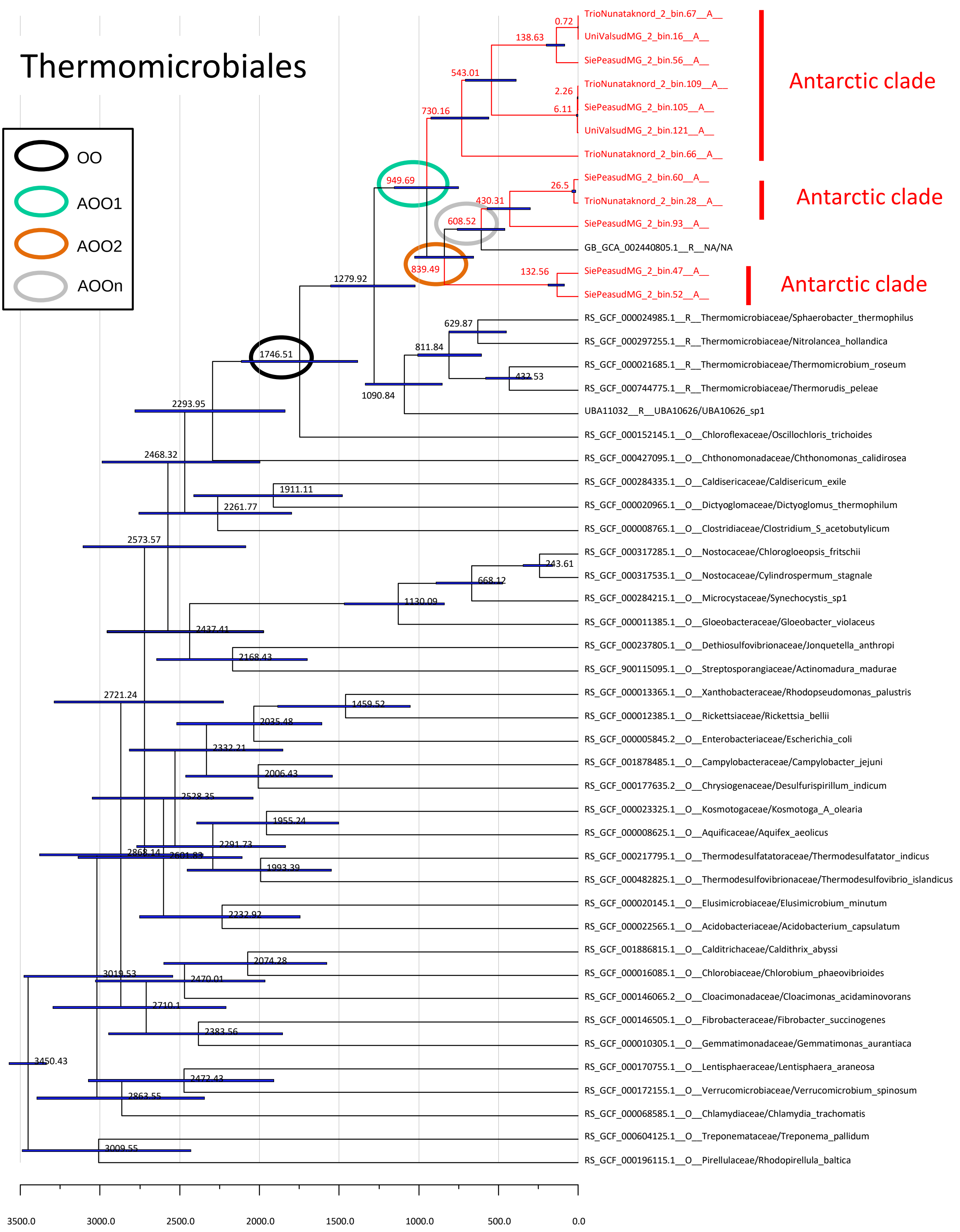

Thermomicrobiales
OO
AOO1
AOO2
AOOn
Antarctic clade
Antarctic clade
Antarctic clade
TrioNunataknord_2_bin.67__A__
UniValsudMG_2_bin.16__A__
SiePeasudMG_2_bin.56__A__
TrioNunataknord_2_bin.109__A__
SiePeasudMG_2_bin.105__A__
UniValsudMG_2_bin.121__A__
TrioNunataknord_2_bin.66__A__
SiePeasudMG_2_bin.60__A__
TrioNunataknord_2_bin.28__A__
SiePeasudMG_2_bin.93__A__
GB_GCA_002440805.1__R__NA/NA
SiePeasudMG_2_bin.47__A__
SiePeasudMG_2_bin.52__A__
RS_GCF_000024985.1__R__Thermomicrobiaceae/Sphaerobacter_thermophilus
RS_GCF_000297255.1__R__Thermomicrobiaceae/Nitrolancea_hollandica
RS_GCF_000021685.1__R__Thermomicrobiaceae/Thermomicrobium_roseum
RS_GCF_000744775.1__R__Thermomicrobiaceae/Thermorudis_peleae
UBA11032__R__UBA10626/UBA10626_sp1
RS_GCF_000152145.1__O__Chloroflexaceae/Oscillochloris_trichoides
RS_GCF_000427095.1__O__Chthonomonadaceae/Chthonomonas_calidirosea
RS_GCF_000284335.1__O__Caldisericaceae/Caldisericum_exile
RS_GCF_000020965.1__O__Dictyoglomaceae/Dictyoglomus_thermophilum
RS_GCF_000008765.1__O__Clostridiaceae/Clostridium_S_acetobutylicum
RS_GCF_000317285.1__O__Nostocaceae/Chlorogloeopsis_fritschii
RS_GCF_000317535.1__O__Nostocaceae/Cylindrospermum_stagnale
RS_GCF_000284215.1__O__Microcystaceae/Synechocystis_sp1
RS_GCF_000011385.1__O__Gloeobacteraceae/Gloeobacter_violaceus
RS_GCF_000237805.1__O__Dethiosulfovibrionaceae/Jonquetella_anthropi
RS_GCF_900115095.1__O__Streptosporangiaceae/Actinomadura_madurae
RS_GCF_000013365.1__O__Xanthobacteraceae/Rhodopseudomonas_palustris
RS_GCF_000012385.1__O__Rickettsiaceae/Rickettsia_bellii
RS_GCF_000005845.2__O__Enterobacteriaceae/Escherichia_coli
RS_GCF_001878485.1__O__Campylobacteraceae/Campylobacter_jejuni
RS_GCF_000177635.2__O__Chrysiogenaceae/Desulfurispirillum_indicum
RS_GCF_000023325.1__O__Kosmotogaceae/Kosmotoga_A_olearia
RS_GCF_000008625.1__O__Aquificaceae/Aquifex_aeolicus
RS_GCF_000217795.1__O__Thermodesulfatatoraceae/Thermodesulfatator_indicus
RS_GCF_000482825.1__O__Thermodesulfovibrionaceae/Thermodesulfovibrio_islandicus
RS_GCF_000020145.1__O__Elusimicrobiaceae/Elusimicrobium_minutum
RS_GCF_000022565.1__O__Acidobacteriaceae/Acidobacterium_capsulatum
RS_GCF_001886815.1__O__Calditrichaceae/Caldithrix_abyssi
RS_GCF_000016085.1__O__Chlorobiaceae/Chlorobium_phaeovibrioides
RS_GCF_000146065.2__O__Cloacimonadaceae/Cloacimonas_acidaminovorans
RS_GCF_000146505.1__O__Fibrobacteraceae/Fibrobacter_succinogenes
RS_GCF_000010305.1__O__Gemmatimonadaceae/Gemmatimonas_aurantiaca
RS_GCF_000170755.1__O__Lentisphaeraceae/Lentisphaera_araneosa
RS_GCF_000172155.1__O__Verrucomicrobiaceae/Verrucomicrobium_spinosum
RS_GCF_000068585.1__O__Chlamydiaceae/Chlamydia_trachomatis
RS_GCF_000604125.1__O__Treponemataceae/Treponema_pallidum
RS_GCF_000196115.1__O__Pirellulaceae/Rhodopirellula_baltica
3500.0
3000.0
2500.0
2000.0
1500.0
1000.0
500.0
0.0

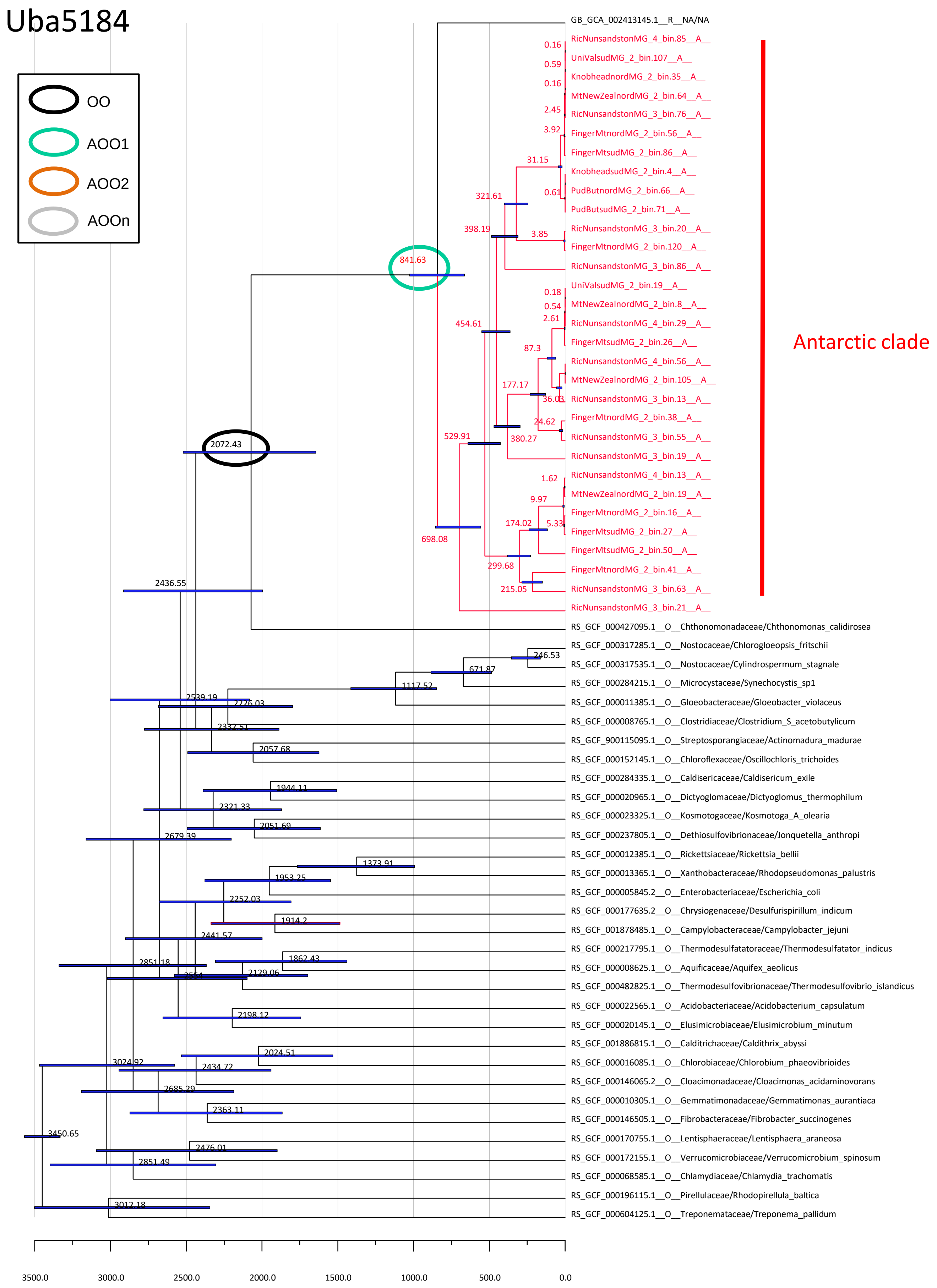
Uba5184
OO
AOO1
AOO2
AOOn
Antarctic clade
GB_GCA_002413145.1__R__NA/NA
RicNunsandstonMG_4_bin.85__A__
UniValsudMG_2_bin.107__A__
KnobheadnordMG_2_bin.35__A__
MtNewZealnordMG_2_bin.64__A__
RicNunsandstonMG_3_bin.76__A__
FingerMtnordMG_2_bin.56__A__
FingerMtsudMG_2_bin.86__A__
KnobheadsudMG_2_bin.4__A__
PudButnordMG_2_bin.66__A__
PudButsudMG_2_bin.71__A__
RicNunsandstonMG_3_bin.20__A__
FingerMtnordMG_2_bin.120__A__
RicNunsandstonMG_3_bin.86__A__
UniValsudMG_2_bin.19__A__
MtNewZealnordMG_2_bin.8__A__
RicNunsandstonMG_4_bin.29__A__
FingerMtsudMG_2_bin.26__A__
RicNunsandstonMG_4_bin.56__A__
MtNewZealnordMG_2_bin.105__A__
RicNunsandstonMG_3_bin.13__A__
FingerMtnordMG_2_bin.38__A__
RicNunsandstonMG_3_bin.55__A__
RicNunsandstonMG_3_bin.19__A__
RicNunsandstonMG_4_bin.13__A__
MtNewZealnordMG_2_bin.19__A__
FingerMtnordMG_2_bin.16__A__
FingerMtsudMG_2_bin.27__A__
FingerMtsudMG_2_bin.50__A__
FingerMtnordMG_2_bin.41__A__
RicNunsandstonMG_3_bin.63__A__
RicNunsandstonMG_3_bin.21__A__
RS_GCF_000427095.1__O__Chthonomonadaceae/Chthonomonas_calidirosea
RS_GCF_000317285.1__O__Nostocaceae/Chlorogloeopsis_fritschii
RS_GCF_000317535.1__O__Nostocaceae/Cylindrospermum_stagnale
RS_GCF_000284215.1__O__Microcystaceae/Synechocystis_sp1
RS_GCF_000011385.1__O__Gloeobacteraceae/Gloeobacter_violaceus
RS_GCF_000008765.1__O__Clostridiaceae/Clostridium_S_acetobutylicum
RS_GCF_900115095.1__O__Streptosporangiaceae/Actinomadura_madurae
RS_GCF_000152145.1__O__Chloroflexaceae/Oscillochloris_trichoides
RS_GCF_000284335.1__O__Caldisericaceae/Caldisericum_exile
RS_GCF_000020965.1__O__Dictyoglomaceae/Dictyoglomus_thermophilum
RS_GCF_000023325.1__O__Kosmotogaceae/Kosmotoga_A_olearia
RS_GCF_000237805.1__O__Dethiosulfovibrionaceae/Jonquetella_anthropi
RS_GCF_000012385.1__O__Rickettsiaceae/Rickettsia_bellii
RS_GCF_000013365.1__O__Xanthobacteraceae/Rhodopseudomonas_palustris
RS_GCF_000005845.2__O__Enterobacteriaceae/Escherichia_coli
RS_GCF_000177635.2__O__Chrysiogenaceae/Desulfurispirillum_indicum
RS_GCF_001878485.1__O__Campylobacteraceae/Campylobacter_jejuni
RS_GCF_000217795.1__O__Thermodesulfatatoraceae/Thermodesulfatator_indicus
RS_GCF_000008625.1__O__Aquificaceae/Aquifex_aeolicus
RS_GCF_000482825.1__O__Thermodesulfovibrionaceae/Thermodesulfovibrio_islandicus
RS_GCF_000022565.1__O__Acidobacteriaceae/Acidobacterium_capsulatum
RS_GCF_000020145.1__O__Elusimicrobiaceae/Elusimicrobium_minutum
RS_GCF_001886815.1__O__Calditrichaceae/Caldithrix_abyssi
RS_GCF_000016085.1__O__Chlorobiaceae/Chlorobium_phaeovibrioides
RS_GCF_000146065.2__O__Cloacimonadaceae/Cloacimonas_acidaminovorans
RS_GCF_000010305.1__O__Gemmatimonadaceae/Gemmatimonas_aurantiaca
RS_GCF_000146505.1__O__Fibrobacteraceae/Fibrobacter_succinogenes
RS_GCF_000170755.1__O__Lentisphaeraceae/Lentisphaera_araneosa
RS_GCF_000172155.1__O__Verrucomicrobiaceae/Verrucomicrobium_spinosum
RS_GCF_000068585.1__O__Chlamydiaceae/Chlamydia_trachomatis
RS_GCF_000196115.1__O__Pirellulaceae/Rhodopirellula_baltica
RS_GCF_000604125.1__O__Treponemataceae/Treponema_pallidum
0.16
0.59
0.16
2.45
3.92
31.15
0.61
321.61
398.19
3.85
841.63
0.18
0.54
2.61
454.61
87.3
177.17
36.03
24.62
529.91
380.27
1.62
9.97
174.02
5.33
698.08
299.68
215.05
2072.43
2436.55
246.53
671.87
1117.52
2539.19
2226.03
2332.51
2057.68
1944.11
2321.33
2051.69
2679.39
1373.91
1953.25
2252.03
1914.2
2441.57
1862.43
2851.18
2129.06
2198.12
2024.51
3024.92
2434.72
2685.29
2363.11
3450.65
2476.01
2851.49
3012.18
3500.0
3000.0
2500.0
2000.0
1500.0
1000.0
500.0
0.0
