## Supplementary figures and images for "Antarctic cryptoendolithic bacterial lineages of pre-Cambrian origin as proxy for Mars colonization"

### Supplementary Fig. 1

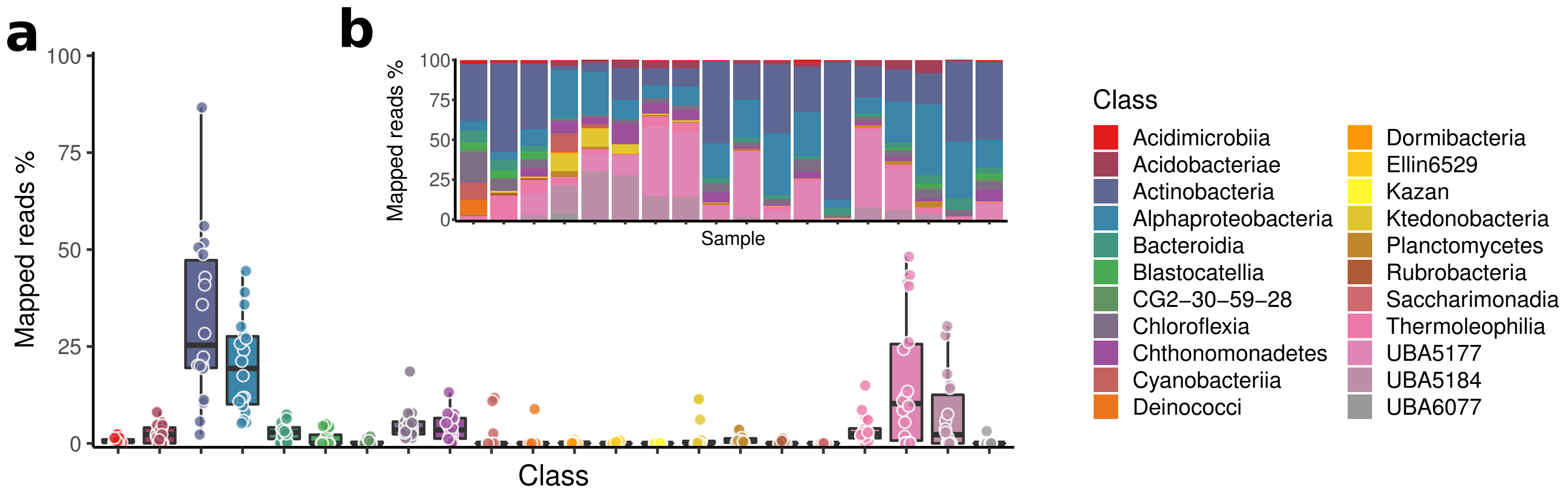

### Supplementary Fig. 2

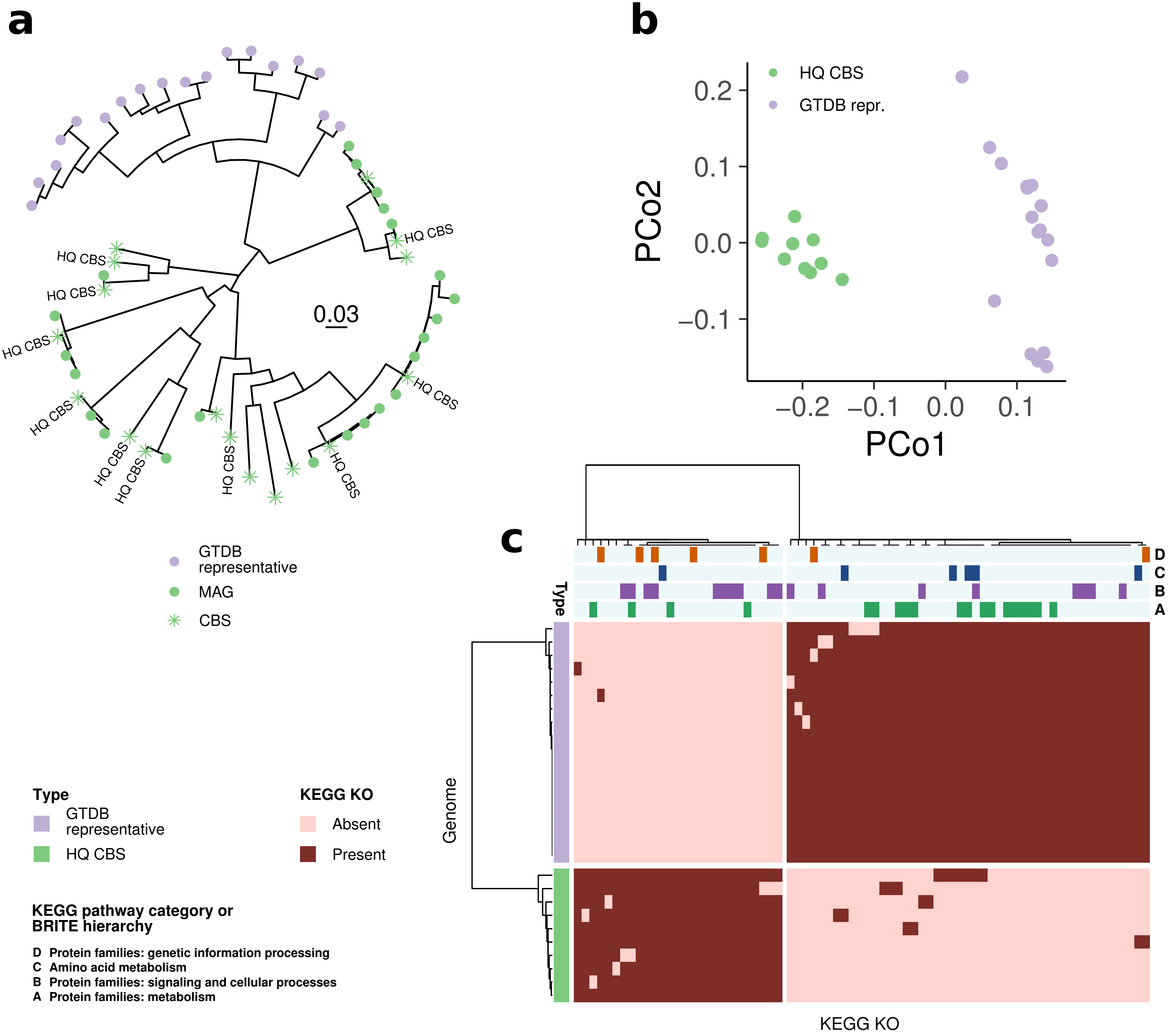
