## Supplementary Text for "Antarctic cryptoendolithic bacterial lineages of pre-Cambrian origin as proxy for Mars colonization"

### Supplementary Figure Captions

**Supplementary Figure 1. a**) Percentage of reads that could be mapped to the CBS representatives, grouped by Class. **b**) Per sample percentage of the reads that could be mapped to the CBS representatives, grouped by Class.

**Supplementary Figure 2.** Antarctic CBS form two distinct clades in the order Frankiales, with characteristic metabolic potential. **a)** Maximum-likelihood phylogenetic tree based on GTDB 120 core genes including representative genomes (violet) and Antarctic CBS (green). **b)** Principal Coordinate Analysis (Jaccard distance) of the metabolic potential of the GTDB representatives (circles) and Antarctic CBS (triangles). **c)** The Fisher's exact test (Bonferroni corrected $p<0.01$) highlights enriched functional categories. Genomes and KEGG orthologs are clustered according to the Hamming distance between the profiles. The top four KEGG categories significantly more present in the Antarctic CBS are highlighted in the upper bars.

### Supplementary Table Captions

**Supplementary Table 1.** Assembly statistics and taxonomic classification of the MAGs.

**Supplementary Table 2.** Abundance of CBS at phylum level, expressed as percentage of reads that could be mapped to the representative CBS. Median: median; Q1 and Q3: first and third quartile; IQR: interquartile range; Mean: mean; SD: standard deviation; #CBS: number of candidate bacterial species belonging to the phylum.

**Supplementary Table 3.** Increase in the number of bacterial species for each taxonomic Order provided by the data in the present study, compared to the data available in the GTDB database.

**Supplementary Table 4.** Sample metadata. Geographic coordinates of the sampling sites, accession numbers of the raw sequences, accession numbers and N50 of the assembled metagenomes on the JGI IMG/M portal.

**Supplementary Table 5.** Prevalence and taxonomic classification for each CBS representatives.

**Supplementary Table 6.** Summary of Bayesian divergence estimates. For each order we report the mean age of its origin (OO: the split of the order from the closest order) and the 95% CI (OO max and OO min), the origin of the oldest uniquely Antarctic clade (AOO1, the split of the Antarctic clade from a non-Antarctic lineage of the same order), and, where present, the origin of the second oldest antarctic clade (AOO2). See Supplementary Data 1.

**Supplementary Table 7.** Number of predicted proteins (NProts) and of proteins that had a match in the EggNOG database (NHitsOG) and that could be associated to a term in the Gene Ontology (NHitsGO) or had a match in the KEGG and COG databases (NHitsKEGG and NHitsCOG, respectively).
